# Cellular mechanisms of acquired drug resistance against regorafenib in colon cancer cells

**DOI:** 10.64898/2026.02.03.703469

**Authors:** Deepu sharma, Fayyaz Rasool, Binayak kumar, Rohit verma, Ashutosh singh, Sri Krishna Jayadev Magani

## Abstract

Colorectal cancer is the third most common cancer across the world. Acquired resistance to therapeutics is one of the major challenges in cancer cure. With the development of resistance to organometallic drugs there was switch for the usage of inhibitors of kinases which play a crucial role in cellular activities. Regorafenib is a multi-kinase inhibitor used as an oral anti-cancer drug for treating advanced colorectal cancer (CRC). With increasing reports of acquired resistance to regorafenib in long-term use it is inevitable to understand the mechanisms underlying in the development of resistance. To understand the molecular mechanism of acquired drug resistance towards regorafenib we have developed a regorafenib resistant HCT116 cell line (Reg-R-HCT116) and an integrated quantitative proteomic and phospho-proteomics approach is used to elucidate the molecular signaling mechanisms that help in drug tolerance.

Proteome and Phosphoproteome analysis revealed an extensive remodelling of signaling pathways associated with metabolism, protein synthesis and stress adaptations. This also revealed a large set of phosphorylated proteins as well as proteins that might be associated with aberrant activation or differential alteration of PI3K-AKT-mTOR, EIF2, HIF-1, Apoptosis inhibition, Glucose metabolism, amino acid metabolism and DNA-repair-associated signaling. Differential phosphorylation of downstream molecules of the mentioned pathways NDRG1, ACINUS and RICTOR and enhanced cell survival further confirmed their role in drug tolerance.

Targeted inhibition of both the mTOR complexes using Torin1 resensitized the cells to regorafenib. Combination treatment of regorafenib with torin1 showed a synergistic cytotoxicity and attenuated the expression of key survival proteins. These findings provide mechanistic insight into acquired resistance to regorafenib in colorectal cancer and identified mTOR/eIF2 signaling as one of the critical drivers of resistance phenotype. The results suggest combinatorial targeting of these pathways could be an effective strategy to overcome regorafenib resistance and improve clinical outcomes in CRC.

## 1. Introduction

Colorectal cancer (CRC) is the third most common cancer globally, with over 1.9 million new cases and 930,000 deaths reported in 2020 (World Health Organization. Colorectal cancer fact sheet, 2024) (Bray et al., 2024). It also known as bowel cancer, developed in the colon or rectum part of the large intestine. Tumour initiates on the luminal surface and progressively invades different layers of the colorectal walls. In advanced stages, tumour cells spread to lymph nodes and finally spread over the body through blood circulation (Centelles, 2012). Chemotherapy and targeted therapy are the primary modes of treatment for advanced-stage colorectal cancer (Cunningham et al., 2010) (DeVita & Chu, 2008). Targeted therapy includes antibody-based immunotherapy and synthetic, small molecule-based kinase inhibitors. Targeted therapy have become important in treating mCRC when combined with chemotherapy i.e BRAF inhibitors, Anti-EGFR and Anti-VEGFR agents, etc (Fernández Montes et al., 2023) (Koelblinger et al., 2018). Regorafenib is a synthetic, small molecule-based, multi-kinase inhibitor being used in advanced stages colorectal cancer. It has multiple membrane-bound and intra-cellular molecules targeted such as RET, VEGFR1,2 and 3, C-KIT, PDGFR-alpha, BRAF, CRAF, FGFR1 and TIE2 etc., which plays crucial roles in cell survival, cell proliferation and angiogenesis (Cheng et al., 2013) (Wilhelm et al., 2011) (Aprile et al., 2013). However, acquired resistance to these receptors and intracellular molecule inhibitors are often observed, which motivates our interest in examining potential resistance mechanisms to regorafenib in colorectal cancer (Goldstein et al., 2015)(Papadimitriou & Papadimitriou, 2021).

Few reports have shown regorafenib-resistance as a therapeutic challenge in colorectal cancer (Koelblinger et al., 2018) (Cai et al., 2018) (Imamura et al., 2016). Chemotherapy failure and disease recurrence mainly occur due to drug resistance, highlighting the importance of understanding the molecular basis of resistance in cancer. (Van Cutsem et al., 2016)(Gottesman, 2002) (Mansoori et al., 2017). By understanding the altered pathways and molecular targets responsible for resistance, novel strategies to reverse it can be developed (Gotwals et al., 2017). Although some resistance mechanisms are known, a comprehensive understanding to it is still lacking. Therefore, this study aims to fill this gap by using proteomic and phospho-proteomic analyses to identify novel molecular targets and signaling pathways involved in regorafenib resistance. Various experimental models have been employed to investigate drug resistance mechanisms, including cell lines, organoids, and genetically modified mice (Goodspeed et al., 2016) (Sales Amaral et al., 2019). Among these, drug-resistant cell line models stand out as essential tools for comprehending resistance development.

Proteomics is a powerful approach to study all proteins in a biological system and has widely used in identifying differentially expressed proteins linked to drug resistance. It enables the identification of biomarkers and therapeutic targets, as demonstrated in ovarian and breast cancers (Raivola, 2022) (Vinet M, 2019). Furthermore, it aids in personalizing treatment strategies and developing novel drug combinations (Wang et al., 2021). However, phospho-proteomics, a subset of proteomics, specifically examines protein phosphorylation, a key post-translational modification. Dysregulated phosphorylation is implicated in diseases including cancer, underscoring its significance in elucidating signaling pathways contributing to resistance (Johnson & Lewis, 2001) (Lee et al., 2023). Notably, phosphor-proteomics analyses in various cancers, including hepatocellular carcinoma and gastric cancer, have shown its role on adaptive resistance mechanisms and potential therapeutic targets (Dazert et al., 2016)(Chen et al., 2020)(Liu et al., 2017).

In this context, this study undertakes a comprehensive analysis of non-labelled proteomics and phospho-proteomics to unravel the mechanisms behind acquired drug resistance to regorafenib. By comparing drug-resistant and sensitive cells, the study aims to identify altered signaling pathways, molecular mechanisms, and potential biomarkers responsible for driving resistance. Ultimately, this study contributes to the broader understanding of drug resistance in colorectal cancer and provides insights that could guide the development of novel therapeutic strategies, ultimately improving patient outcomes.

The various signaling pathways, such as PI3K-AKT-mTOR proteins, play crucial roles in controlling the activity of several downstream molecules, which are responsible for controlling the activity of several other signaling pathways that regulate normal cellular functions like cell growth and proliferation. Alterations in PI3K-AKT-mTOR proteins and signaling activated by them have been reported as mediators for chemo-resistance (Liu et al., 2020). Alterations in PI3K-AKT-mTOR signaling, activation of JAK-STAT and RAF-MEK-ERK signaling have been implicated in therapy resistance (Chappell et al., 2011)(Cao et al., 2019).

So, in this study, we have developed a regorafenib resistance cell model and tried to explore the mechanisms of acquired therapy resistance in the same model through the study of the proteome profile as well as the phospho-proteome profile of resistance models. By analysis both the omics studies our spotlight shifts to key pathways including the profiling of the PI3K/Akt/mTOR pathway, the eIF2 pathway, HIF1 signaling, as well as the translation machinery involving ribosomal proteins, and energy metabolism, particularly hexokinases.

## 2. Materials and Methods

### 2.1 Cell lines and Development of Regorafenib-Resistant-HCT-116 cell line model

HCT116 cells, a human colorectal cancer cell line, were cultured in Dulbecco’s Modified Eagle Medium (DMEM) Gibco Thermo Fischer Scientific, containing 10% fetal bovine serum (FBS) Brazil origin, Gibco, Thermo Fischer Scientific and 1% antibiotics (100 U/mL penicillin G and 100 mg/mL streptomycin), in a humidified incubator with 5% CO2 at 37°C. These cells were continuously treated with regorafenib (BAY 73-4506), a multi-tyrosine kinase inhibitor, starting with an initial concentration of 10 nM and progressively increased the concentration till the cells could tolerate and survive. This continuous treatment was done for over 12 months. The development of resistance was assessed from time to time by MTT assay. Cell lines were regarded to be resistant to the drug if the fold resistance factor was ≥2 (Reg-R-HCT116) and used for further experiments. Cell Lines were authenticated by STR (Short Tandem Repeat) profiling.

### 2.2 Reagents and Antibodies

Primary antibodies Beta-actin antibody (1:5,000) (ImmunoTag, Cat No. ITT07018, S. Rabbit), PCNA (1:1000) (ImmunoTag, G-biosciences, Cat No. ITP0016, S. Rabbit), anti-Bcl-2 (1:1,000) (ImmunoTag, G-biosciences, Cat No. ITM3041, S. Mouse), cleaved PARP (1:1,000) (ImmunoTag, G-biosciences, Cat No. ITT07023, S. Rabbit), anti-P-NDRG1 thr346(1: 1000), P-ACINUS (1: 1000), ERK (1:1,000) Antibody - BD Biosciences, Mouse, Cat No. 610123, RPS6 (1:1000) Antibody – Cell signaling Technology (CST), Rabbit, Cat No. #2217,eIF4E (1:1000) Antibody – Cell signaling Technology (CST), Rabbit, Cat No. #2067, Rictor (1:1000) Antibody – Cell signaling Technology (CST), Rabbit, Cat No. #2114, p-4E-BP1 (Thr37/46) (1:1000) Antibody –CST, Rabbit, Cat No. #2855, Raptor (1:1000) Antibody - ImmunoTag, Rabbit, Cat No. ITT07527, p-RPS6 (Ser-235) (1:1000) Antibody - ImmunoTag, Rabbit, Cat No. ITP03354, EIF4A3 (1:1000) Antibody - ImmunoTag, Rabbit, Cat No. ITT12071, EIF4B (1:1000) Antibody - ImmunoTag, Rabbit, Cat No. ITT07324, EIF5 (1:1000) Antibody - ImmunoTag, Rabbit, Cat No. ITT09511, antibodies at 4°C for 10-12 hour. Goat anti-Rabbit IgG/HRP 1:5,000 ImmunoTag G-biosciences, Cat No. SE134, Goat anti-Mouse 1:5,000 BD Biosciences Cat No. 554002). RNeasy Mini Kit from Qiagen, Thermo Scientific Verso cDNA synthesis kit, 2xSYBR green master mix obtained from Promega, JC-1 staining (Invitrogen, REF M34152), carbonyl cyanide m-chlorophenyl hydrazine (CCCP, Invitrogen, REF M34152)

### 2.3 Measurement of cell viability

The cytotoxicity and drug tolerance of HCT-116 and Reg-R-HCT116 cells for Regorafenib was checked by MTT assay. 2×10^4^ cells per well were seeded and incubated overnight in flat bottom 96-well culture plates (Thermo Nunc). Cells were treated with a concentration gradient of regorafenib from 0.5 to 10 µM for 48 hrs. After treatment, the media was discarded, and the cells were incubated for 3 hours in fresh DMEM containing 0.5 mg/ml MTT (G-Biosciences India). The MTT containing media was discarded, and the formazan crystals were dissolved in 100µl DMSO, and absorbance was measured at 595nm wavelength in a Bio-Rad multi-plate reader. All the experiments were repeated thrice, and percentage viability was analysed using the formula Percentage viable cells = (mean OD of treated cells/mean OD of control cells) X 100.

### 2.4 Propidium Iodide (PI) staining and Microscopy

The cytotoxicity of regorafenib on HCT-116 and Reg-R-HCT116 cells was further confirmed by PI live /dead exclusion experiment. 5X 10^4^ cells were seeded in the 6-well plates and then treated with different concentrations of regorafenib for defined time points as reported by MTT assay. After drug treatment, the cells were washed with PBS and incubated with 20µg/ml propidium Iodide for 15 minutes at room temperature. The cells were then washed twice with PBS and imaged using a Nikon inverted fluorescence microscope with white light, DAPI and TRITC channels at 100X magnification.

### 2.5 JC-1 Staining for Mitochondrial Membrane Potential Loss:-

To understand the process of cell death, alterations in mitochondrial membrane potential (MMP) was examined through JC-1 staining (Invitrogen, REF M34152) on cells subjected to IC50 concentration of drug treatment. Cells treated with Regorafenib, Torin 1, and Reg/Tor1 for 48 hours were subjected to JC1 staining at a final concentration of 2 µM and left to incubate for 15–20 minutes. The cells were rinsed with PBS and were imaged using a Leica fluorescence microscope at 200X magnification. Flow cytometric analysis was also performed with the JC-1stained cells. Carbonyl cyanide m-chlorophenyl hydrazine (CCCP, Invitrogen, REF M34152) was used as a positive control.

### 2.6 Western Blot analysis

HCT116 and Reg-R-HCT116 cells were seeded into 6-well culture plates and were treated with respective drugs once cells were attached and attained proper morphology. Cells were harvested and lysed in a cell lysis buffer (RIPA buffer) with protease and phosphatase inhibitors. Lysates were quantified, and 30µg of total protein was resolved on SDS-PAGE. The resolved proteins were transferred on to the PVDF membrane. Protein transferred membranes were blocked with either 5% skim milk (for non-phosphorylated proteins) or 3% BSA fraction V (phosphorylated proteins) for 1 hour at room temperature. The blots were probed with respective primary antibodies for 1 hour and then washed with PBST. The blots were then incubated with respective secondary antibodies for 1 hr at room temperature. The unbound secondary antibody was washed with 1X TBST. Blots were detected by the enhanced chemiluminescence (ECL) detection reagent (BioRad) using FluorChem E system (Protein simple) to visualize the signals.

### 2.7 Real-time PCR

5 x10^4^ cells of HCT-116 and Reg-R-HCT-116 cells were seeded into each well in a 6-well culture plate and incubated overnight. Cells were treated with respective drugs for 24-48 hours as mentioned above. Total RNA was isolated using the RNeasy Mini Kit from Qiagen per the given protocol. The quantity and quality of RNA were measured by Nano-Drop. 100 ng of total RNA was used for cDNA synthesis using Thermo Scientific Verso cDNA synthesis kit using the manufacturer’s protocol. 2 µl of cDNA is used as a template for real-time qPCR. 2xSYBR green master mix obtained from Applied Biosystem was used. Experiments were done in triplicates. Reaction were prepared as per manufacturer’s protocol. The Ct-value corresponding to each reaction was recorded and the average of duplicate reaction was calculated. 18S rRNA gene was used as an internal control. Fold change in the expression of different genes was plotted using **GraphPad Prism 8**.0 (2019). Gene list and their primer sequences are listed in supplementary data S1.

### 2.8 Immunofluorescence Assay (IFA), Immunocytochemistry

To validate the western blotting data and visualize the intracellular localization of proteins HCT116 and Reg-R-HCT116 cells were seeded in chambered slides. After 24 hours of incubation, the cells were treated with regorafenib, while untreated cells were used as controls. After treatment with the drug the cells were fixed with 4% paraformaldehyde in PBS for 15 minutes and were permeabilized using 0.1% Triton-X 100 in 1X PBS for 10 minutes at room temperature. Subsequently, the cells were washed three times with 1X PBS and blocked with BSA (fraction V) for 1 hour at room temperature. Thus, blocked cells were incubated with primary antibodies [p-NDRG1 (1:250)] and [p-ACINUS (1:250)] at 4°C for 8-10 hours in a humidified chamber. The slides were washed thrice with 1X PBS and incubated with corresponding secondary antibodies (Alexa Flour 488 anti-rabbit IgG or Alexa Flour 596 anti-mouse IgG at a dilution of 1:2000) at room temperature for 1 hour in a humidified chamber. Subsequently, the slides were washed and counterstained with 4’, 6-diamidino-2-phenylindole (DAPI) at a concentration of 1 µg/ml. Finally, the slides were examined using a confocal microscope (Nikon Fluorescence microscope) at 200X magnification to detect the signals associated with the stained proteins and nuclei.

### 2.9 Visualization of Mitochondria by Confocal Analysis

Mito Tracker Deep Red was employed to observe the morphology and structural integrity of mitochondria. Cells were seeded in a glass bottom plate of 35mm diameter. A total of 6 × 10^4^ cells per well were plated and allowed to attach for 20-24 hours. These cells were then incubated for 15-20 minutes with 200 nM concentration of Mito Tracker Deep Red. Subsequently, the cells were visualized at 100X magnification using a Nikon Confocal Microscope.

### 2.10 Scratch and Wound Healing Assay

In a 12-well plate, 1 × 10^4^ cells per well were seeded for both Control and Resistant cell lines, achieving 60% confluency. Before treatment a scratch was carefully created at the center of each well using a 10 µL tip. Subsequently, the cells were treated with varying concentrations of drugs. Cell imaging was conducted at 0-, 24-, and 48 hours post-treatment and alterations in the scratch were examined. The microscopy analysis was performed using a Leica microscope equipped with a standard DIC filter at 100X magnification.

### 2.11 Liquid chromatography-tandem mass spectrometry (LC-MS/MS) and data analysis

To elucidate the mechanisms involved, a systematic quantitative comparison of the proteome and phosphor-proteome of drug-resistant cells Reg-R-HCT116 and parental cells HCT116 using mass spectrometry was performed. Three biological replicates each were used for proteome, and two biological replicates were used for phosphor-proteome analysis to meet the high reproducibility of sample preparation.

### 2.12 Sample preparation and Mass Spectrometric Analysis of Peptide Mixtures

Cells cultured to 80% confluence, washed 5 times with chilled 1X PBS (Himedia, India) and serum starved for 8 hours, and lysed using cell lysis buffer. Cell lysate prepared in GN lysis buffer with protease inhibitors (Roche) and phosphatase inhibitors (Roche) and boiled at 95°C for 10 min. The supernatant was collected after sonication and centrifugation. Collected supernatant (lysate) was stored at-80 °C. Protein quantification and quality check were performed using Bradford’s reagent and Comassie blue after running SDS-PAGE.

25 µl samples were reduced with 5 mM TCEP, alkylated with 50 mM iodoacetamide, and then digested with Trypsin (1:50, Trypsin/lysate ratio) for 16 h at 37 °C. Trypsin digested samples were cleaned using a C18 silica cartridge, dried, and resuspended in buffer A (5% acetonitrile, 0.1% formic acid).

All the experiment was performed using EASY-nLC 1000 system (Thermo Fisher Scientific) coupled to Thermo Fisher-QExactive equipped with a nano electrospray ion source.

Peptides were loaded and eluted using a gradient, analysed using a data-dependent top10 method, dynamically choosing the most abundant precursor ions from the survey scan.

### 2.13 Proteome data analysis

Generated raw files were analysed with Proteome Discoverer (v2.2) against UniProt HUMAN reference proteome database. For the Sequest search, the precursor tolerance 10ppm, fragment mass tolerances were set at 0.5 Da, trypsin specificity, Carbamidomethyl on cysteine as (fixed), oxidation of methionine and N-terminal acetylation were considered variable modifications for database search. Differential protein quantitation analysis was performed by Protti R (0.6.0) package. Briefly, after data normalization correlation and PCA analysis were carried out. The data filtration was done by using a filter of more than one identified peptide. Significant proteins identified with 0.05 FDR,1.2-fold change cut off. Lists of significantly altered proteins obtained can be subjected to further analysis using functional (Ashburner et al., 2000) or pathway enrichment analyses, as well as a network analysis tool based on the STRING dB package (Szklarczyk et al., 2019) and data annotation with Uniprot and KEGG databases.

### 2.14 Phospho-proteome data analysis

Raw files were analysed using Phosphomatics for global phospho-proteomics data. Raw data files containing phosphorylation sites and protein identification were uploaded. Control HCT116 and Regorafenib-resistant HCT116 groups were created, and the missing values are handled by filtering and median imputation. Log2 transformation was applied for normal distribution and median normalization method was used to normalize our data. Phoshositeplus and Signor databases were selected for analysis.

For Sequest HT and MS Amanda 2.0 searches, set precursor and fragment mass tolerances to 10 ppm and 0.5 Da, respectively, with Trypsin specificity and variable modifications. Both peptide spectrum match and protein false discovery rate were set to 0.05 FDR and determined using a percolator node. The Minora feature detector node of Proteome Discoverer 2.2 was used for relative protein quantification, with annotations assigned using Uniprot, Pfam, KEGG, and GO. High sensitivity-phospho site localization was achieved with the ptmRS node.

## 3. Results

### 3.1 Validation of Reg-RHCT116 cells for regorafenib tolerance

Regorafenib-resistant colorectal cancer cell lines were generated by continuous exposure to increasing concentrations of regorafenib. Control HCT116 and Reg-RHCT116 cells were treated with regorafenib with a gradient of increasing concentration from 1µm to 16µm, and IC50 values were evaluated. The IC50 values for control HCT116 cells were recorded around 2.7µm concentration of regorafenib. However, the cell death was observed beyond 3µm concentration. Reg-RHCT116 cells shows IC50 value around 12 µm concentration of the drug. Beyond 12µm cells started showing cell death gradually. Along with the drug tolerance capabilities of the cells, we could see a change in the morphology of resistant cells compared to parental HCT116 cells **Figure 1B**. The Reg-RHCT116 cells started to grow as individual colonies or patches, rather than showing a uniform monolayer, and shows elongated fusiform structure. These results indicated that continuous treatment with the drug had increased the tolerance capacity of the cells, thus conferring resistance. There is a 4-fold increase in IC50 concentration in resistant cells with respect to parental cells for regorafenib **Table 1**. Further the drug tolerance capabilities were evaluated by PI live dead exclusion assay. Control HCT-116 and REG-RHCT-116 cells were seeded into 6 well plates, treated with 3µM concentration of regorafenib for 48 hrs, and stained with Propidium Iodide as described in materials and methods. As shown in **Figure 1C**, untreated control (HCT-116) cells look healthier, and very few cells were stained red with PI, whereas the control cells treated with the drug showed a stressed phenotype, and comparatively the number of cells stained with PI increased, showing sensitivity of these cells towards the drug. However, the very few resistant cells picked up PI, and most of the cells looked healthier even during drug treatment.

**Figure 1.**
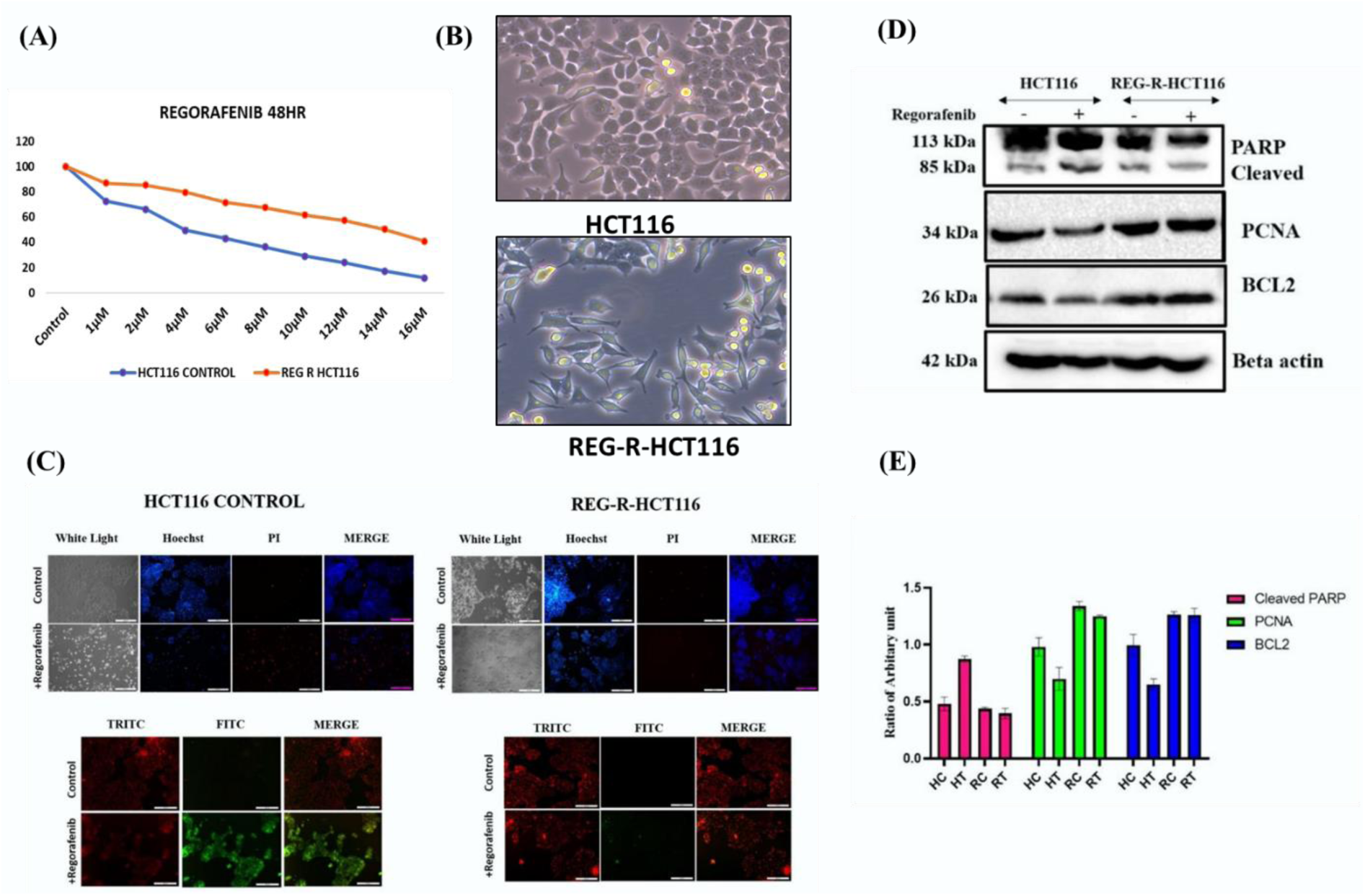
Increased tolerance of regorafenib in HCT116 cells and change in morphology of HCT116 cells. (A) Dose-response curves of regorafenib on HCT116 and Reg-R-HCT116 cells. The Effect of regorafenib on proliferation in HCT116 and Reg-R-HCT116 cells was done by MTT assay after 48 h of treatment. The data shown are mean ± SEM of three replicate wells. The HCT116 and Reg-R-HCT116 cells were treated with 1-16 µM of regorafenib. A graph was plotted with the inhibitor concentration (X-axis) vs cell death % (Y-axis) and the IC50 values were calculated. (B) Microscopic images of HCT116 and Reg-R-HCT116 cells showing a change in the morphology. (C) Propidium iodide exclusion assay to check live/dead cells to show the drug tolerance capability of Reg-R-HCT116 cells. JC-1 staining shows the loss of membrane potential in HCT116 cells with increased green fluorescence, whereas intact mitochondria in Reg-R-HCT116 cells. (D) Western blot analysis of the HCT116 sensitive and Reg-R-HCT116 Resistant cells. Anti-apoptotic protein BCL-2, apoptotic marker PARP cleavage and Proliferating marker PCNA levels were detected by western blot in both HCT116 sensitive and Reg-R-HCT116 Resistant cells. Beta-actin served as loading control. (E) The comparison of the quantified protein densitometric analysis using ImageJ software Cleaved PARP product, Bcl-2 and PCNA are shown to indicate the statistically significant difference (p<0.05).

**Table 1.** IC50 values of the Regorafenib in HCT116 and Reg-R- HCT116.

| Sl.no. | Cell Line | IC50 Values (μM) |
| --- | --- | --- |
| 1. | HCT116 | 2.7±0.6 |
| 2. | Reg-R-HCT116 | 12±0.5 |

The resistance was further confirmed by different markers for apoptosis. Loss of mitochondrial membrane potential a hall mark feature of apoptosis was checked using the JC-1 staining as described in the materials and methods. As shown in Figure 1C, with drug treatment an increase in green fluorescence was recorded in sensitive cells but the resistant cells appeared normal and exhibited red fluorescence further confirming the integrity of mitochondria and the drug tolerance capabilities.

Further, western blot analysis was performed to check the expression of apoptotic marker and cell survival-associated proteins. As shown in Figure 1D, the level of cleaved PARP in drug-treated HCT116 cells showed a significant increase, but the levels in Reg-R-HCT116 showed equal levels to that of control HCT116 cells, confirming the drug tolerance capability of Reg-R-HCT116 cells. Further, the expression of Proliferating Cell Nuclear Antigen (PCNA) levels was checked. The expression of PCNA was found to increase in Reg-R-HCT116 cells. Though we could find a slight decrease in PCNA levels in drug-treated Reg-R-HCT116 cells, they still have a higher expression than the HCT116 cells. A prominent reduction in the expression levels in drug-treated HCT116 cells supports the anticancer properties of regorafenib. Thus, the levels of PCNA suggest resistance development in Reg-R-HCT116 cells.

A significant increase in the expression of pro-survival Bcl-2 in Reg-R-HCT116 was recorded compared to HCT116. A decrease in Bcl-2 levels was noticed in drug-treated HCT 116 cells, thereby supporting the cell death induction by the drug in sensitive cells. Meanwhile, in REG-R-HCT116 cells, we could observe an increase in Bcl-2 levels even in the drug-treated cells, confirming the resistance of these cells to drug-induced cell death.

### 3.2 Comparative proteome profile analysis of HCT116 and Reg-R-HCT116 Cells

Further to understand the molecular alteration in drug resistant cells in comparison to sensitive HCT116 cells quantitative proteomic analysis was performed. The proteome profiling is aimed at identifying deregulated proteins and their abundance levels to characterize the signaling mechanism involved in Reg-R-HCT116 cells.

This proteome data obtained was matched against the NCBI database using proteome discoverer 2.2. A total of 6558 peptides representing 1466 proteins were identified. The proteins within a group were ranked according to the number of peptide sequences, the number of protein sequence motifs (PSMs), their protein scores, and the sequence coverage. Using an FDR cutoff of 0.05, 1466 proteins were found altered, out of which 198 proteins were upregulated (log 2-fold change ≥1) Table 2 and 47 proteins were found downregulated (log 2-fold change ≤1) Table 3. In Figure 2A, the Venn diagram analysis shows common and specific altered proteins in HCT116 and Reg-R-HCT116. 71 proteins were identified as specifically expressed only in Reg-R-HCT116 cells are shown in Table 4, and 48 proteins were found to be expressed only in Control cells are shown in Table 5, whereas 1347 proteins were found differentially expressed and expressed in both the samples.

**Figure 2.**
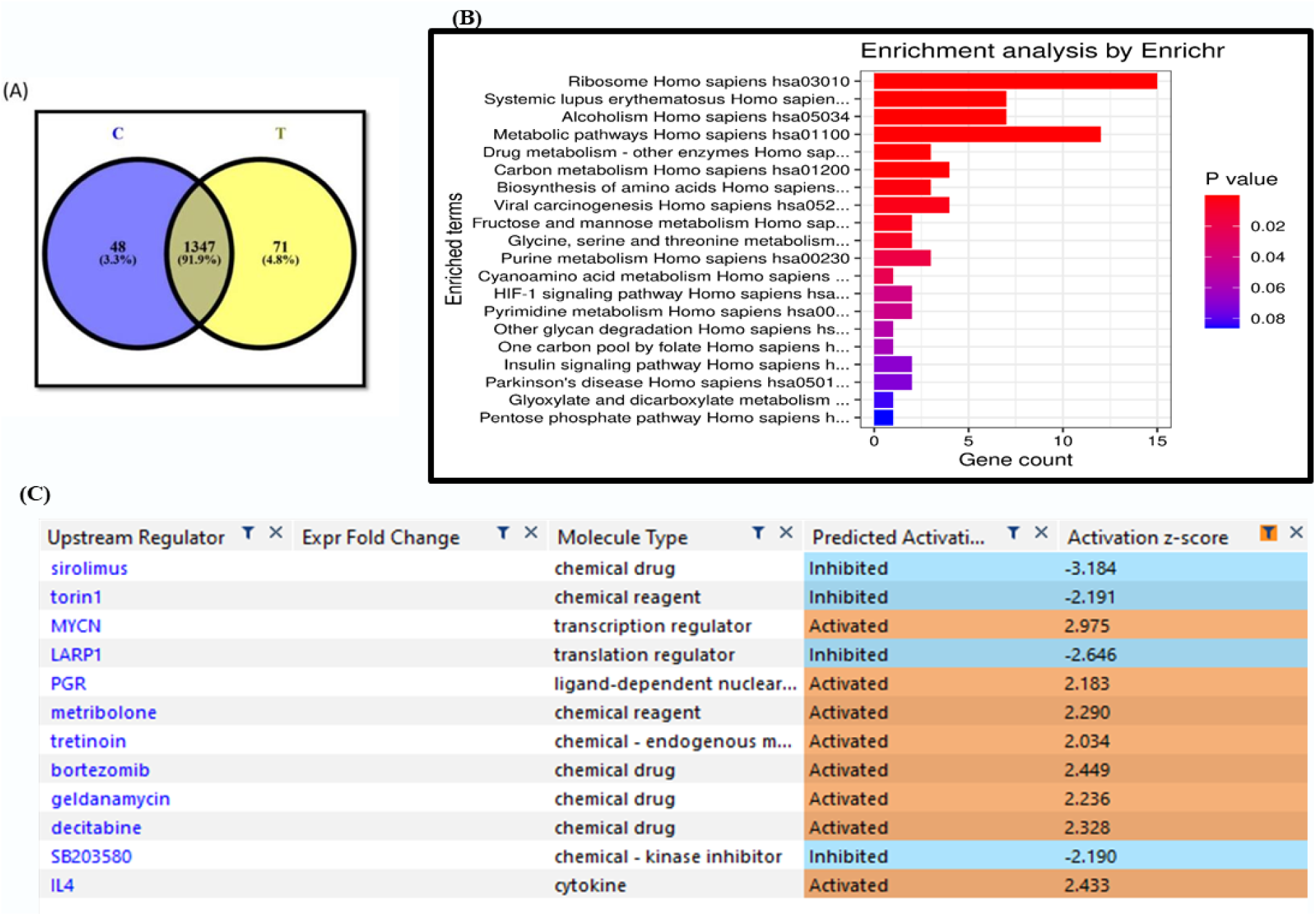
Analysis of altered proteome in Reg-R-HCT116 (A)Venn diagram shows total number of proteins found after analysis of C-HCT116 and Reg-R-HCT116 samples. (B) Pathways Enrichment Analysis. This graph shows the gene ontology (pathways) of the significant proteins found in proteomics data. (C) Represents top regulators of Ribosome associated pathway i.e. eIF2 signaling, with the activation and inhibition z-score value, generated by IPA.

**Table 2:**
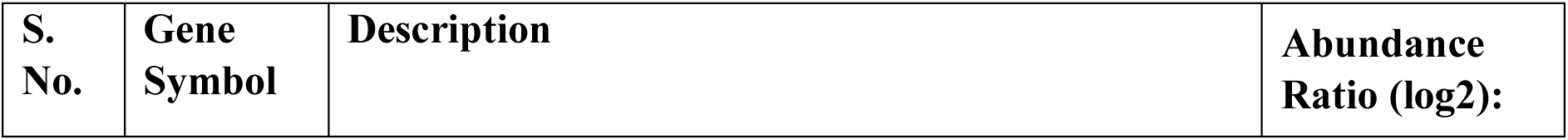
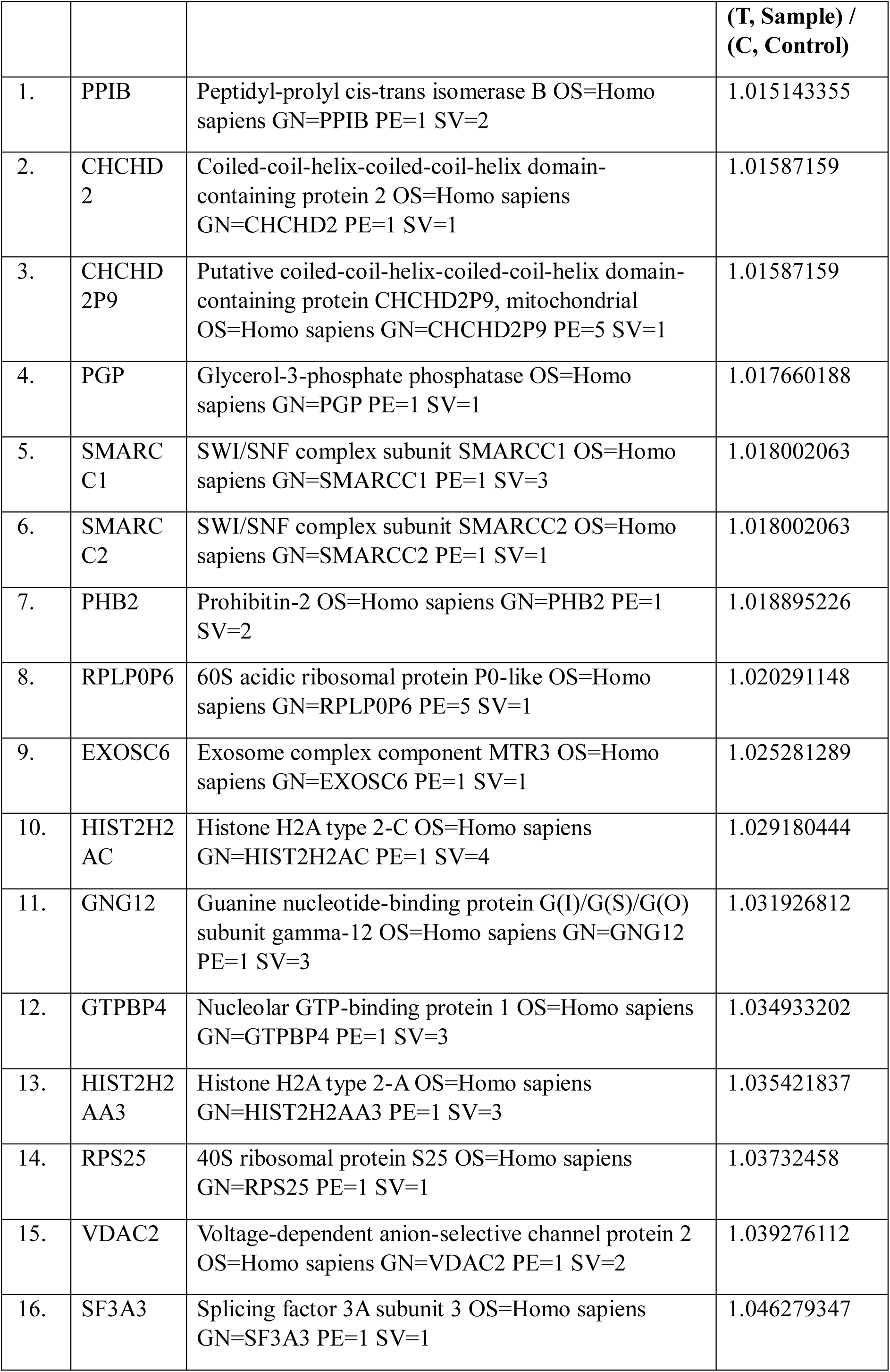
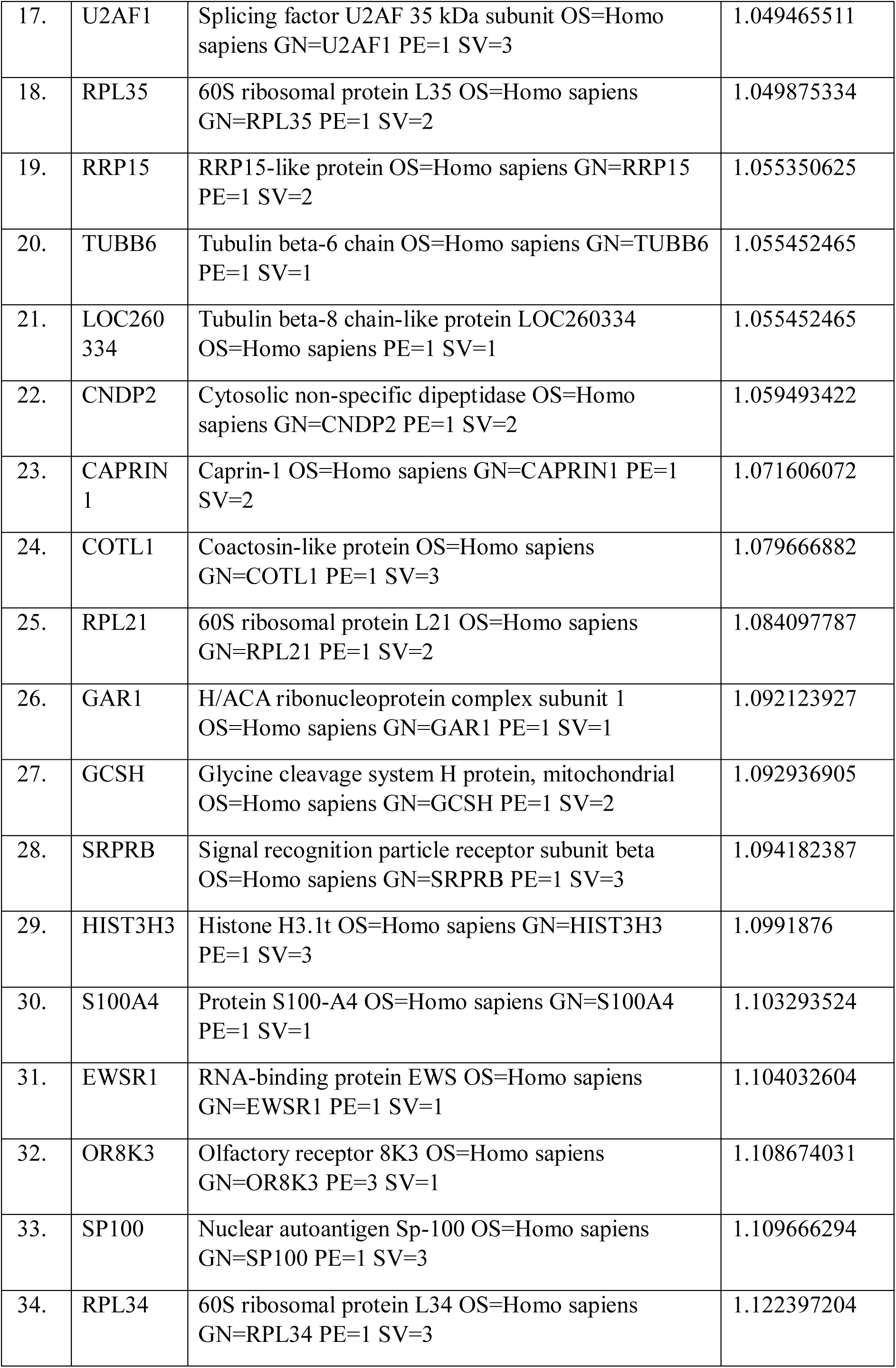
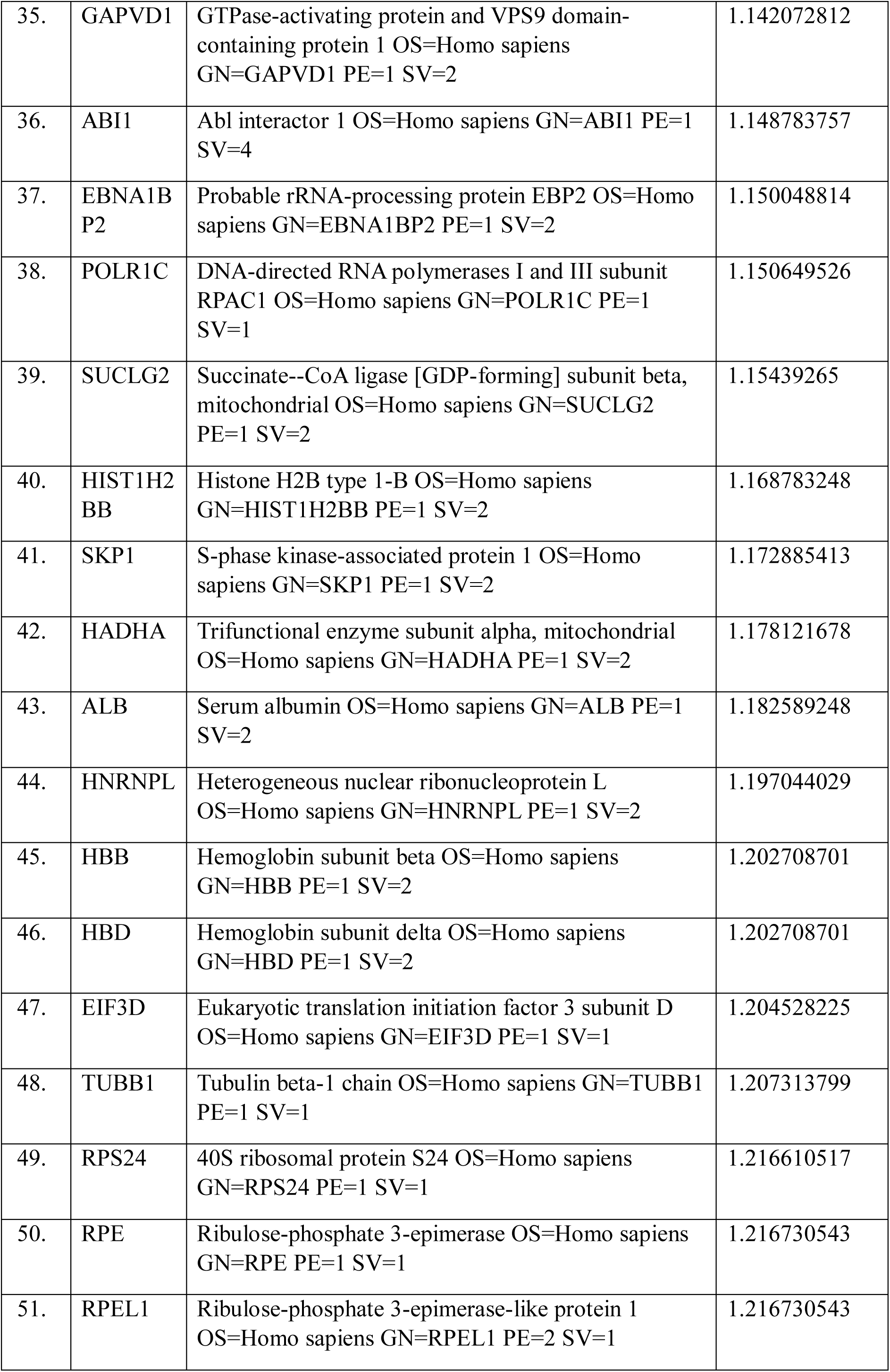
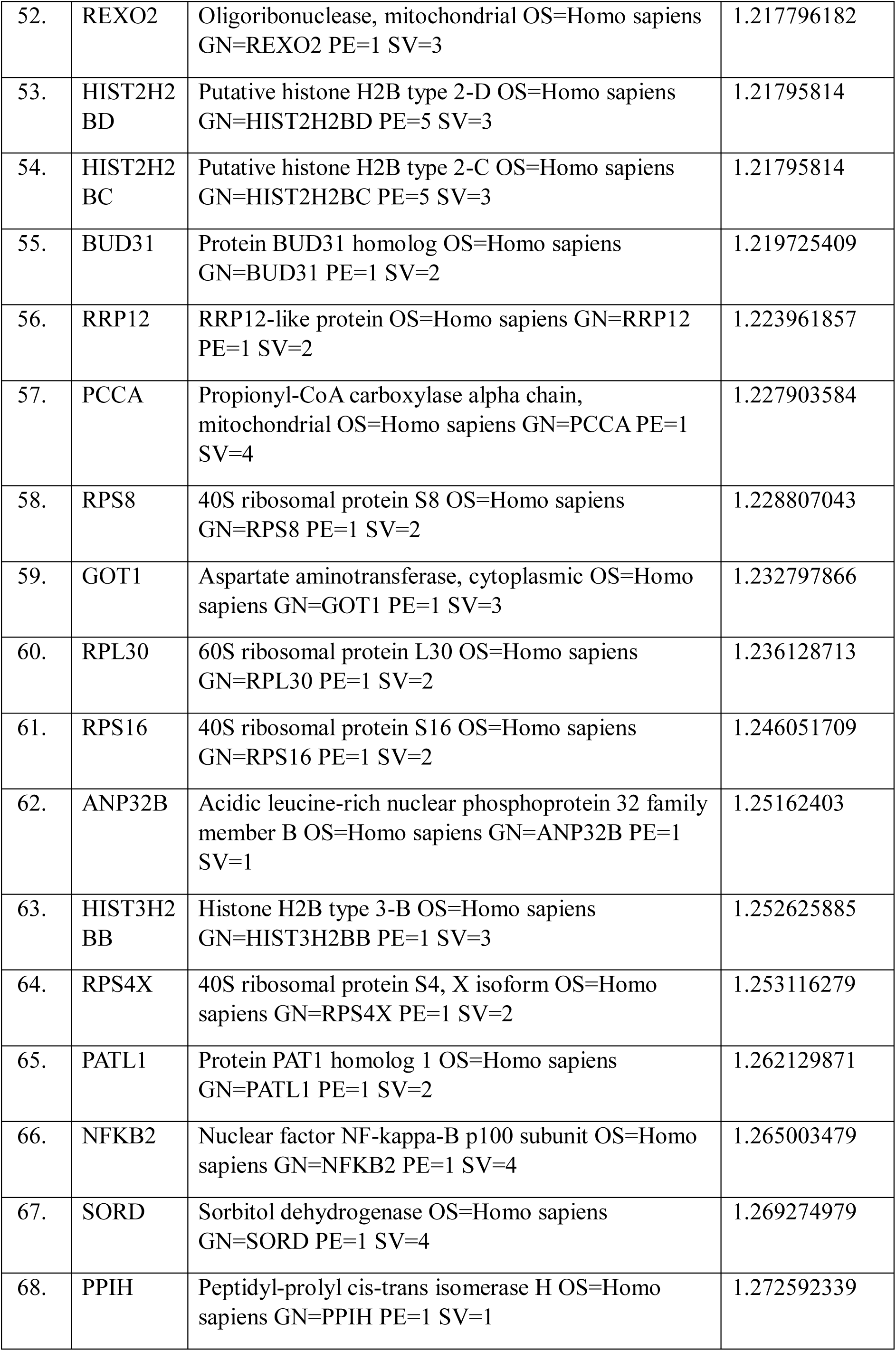
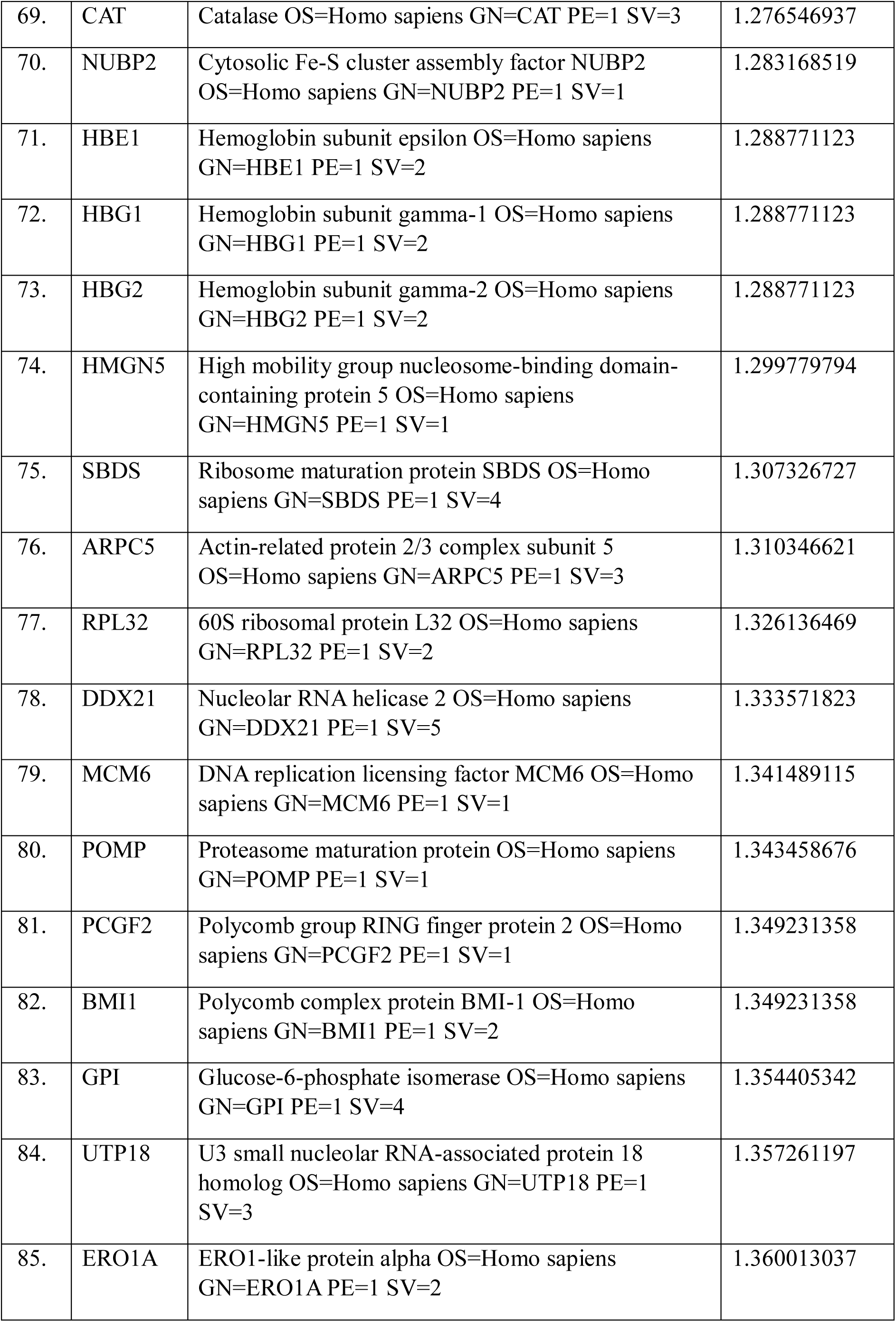
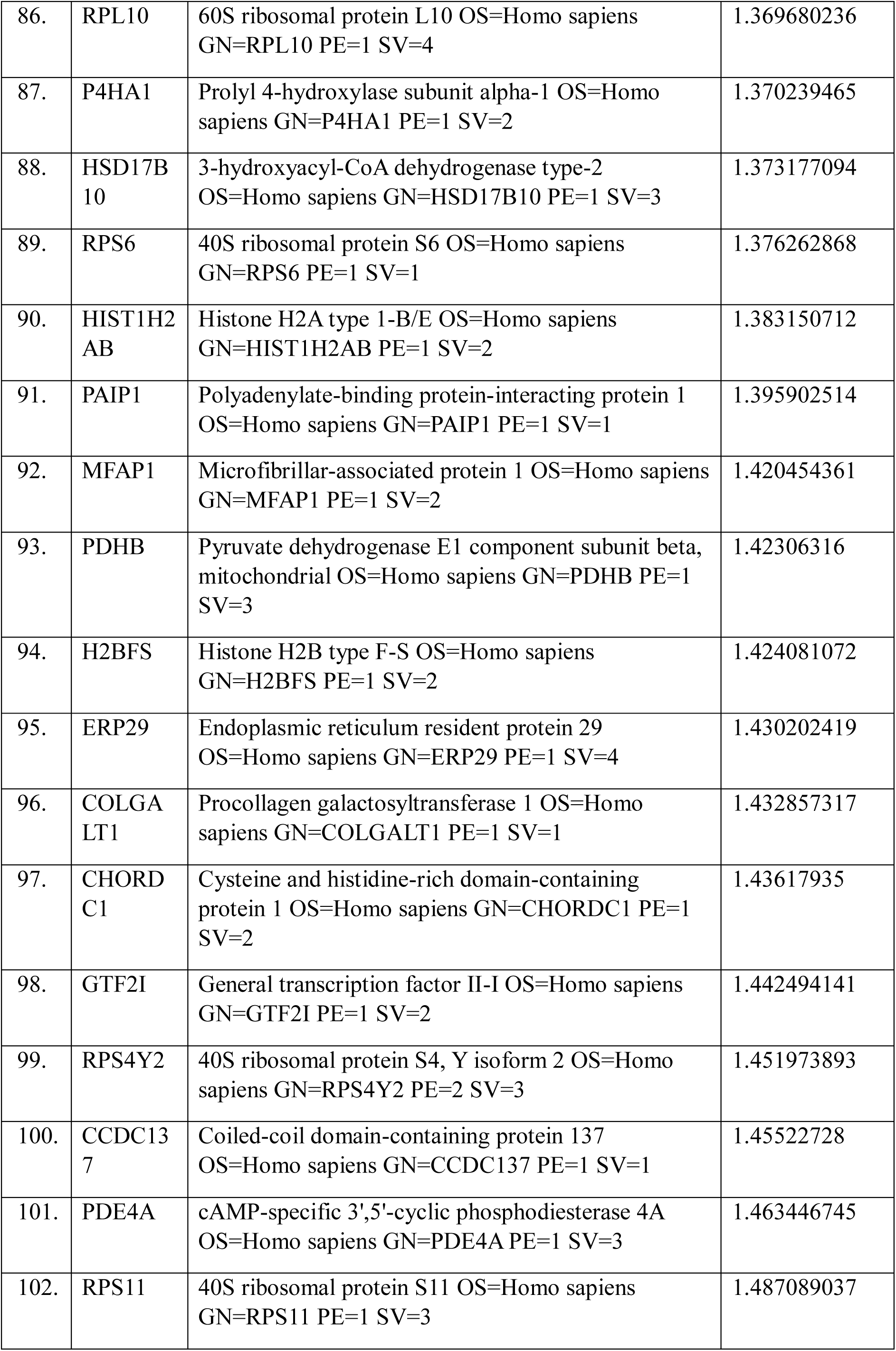
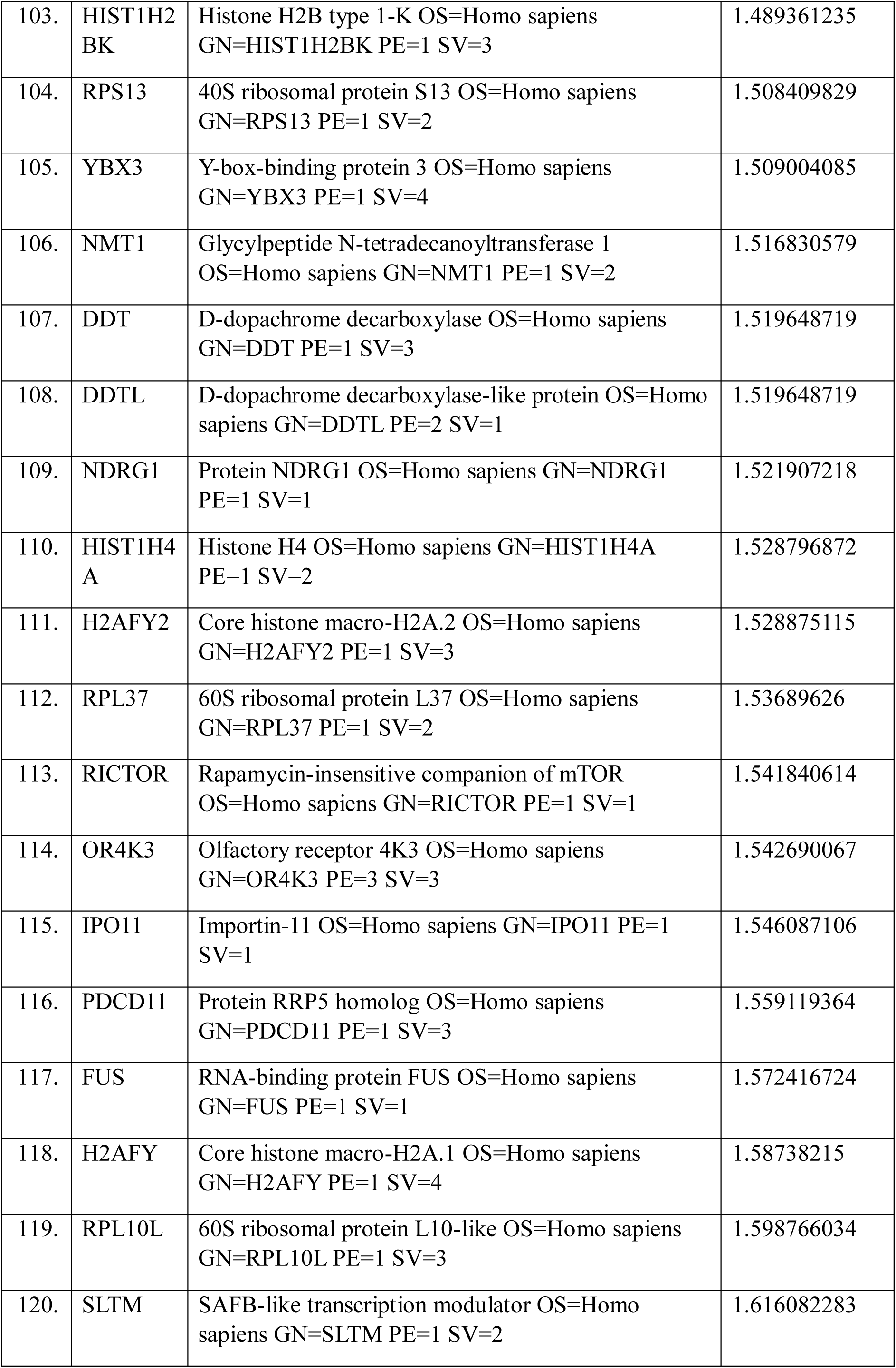
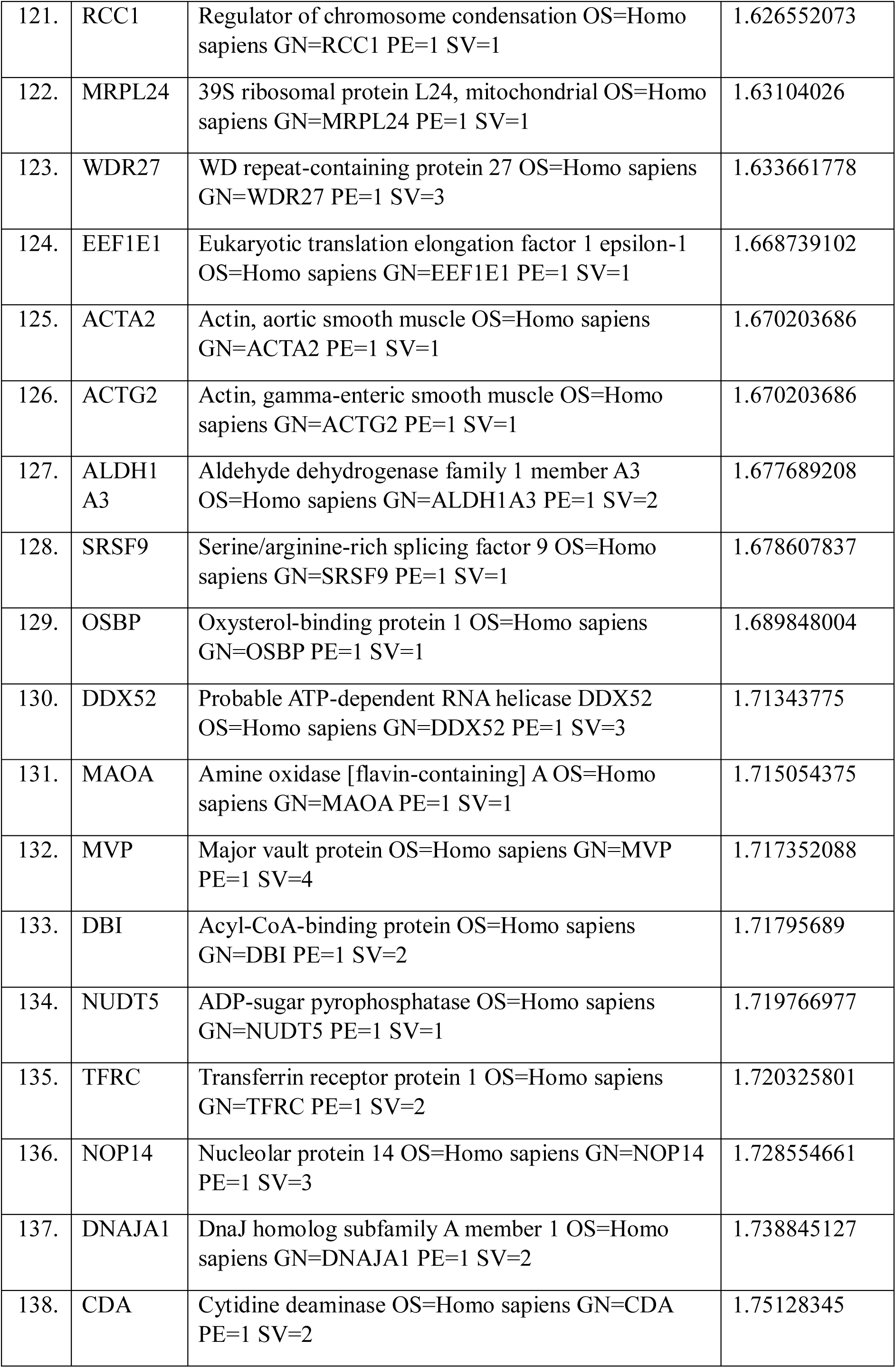
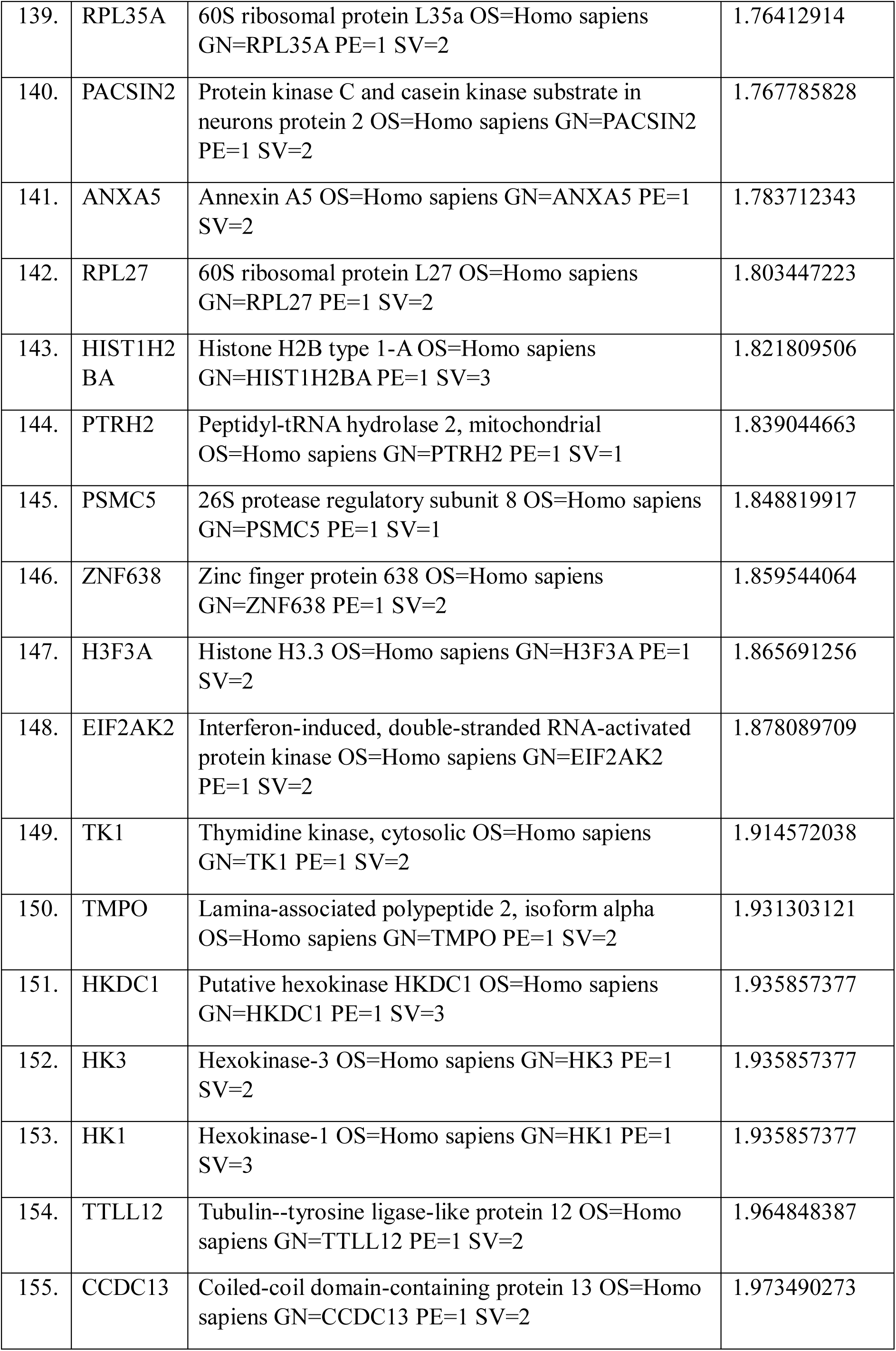
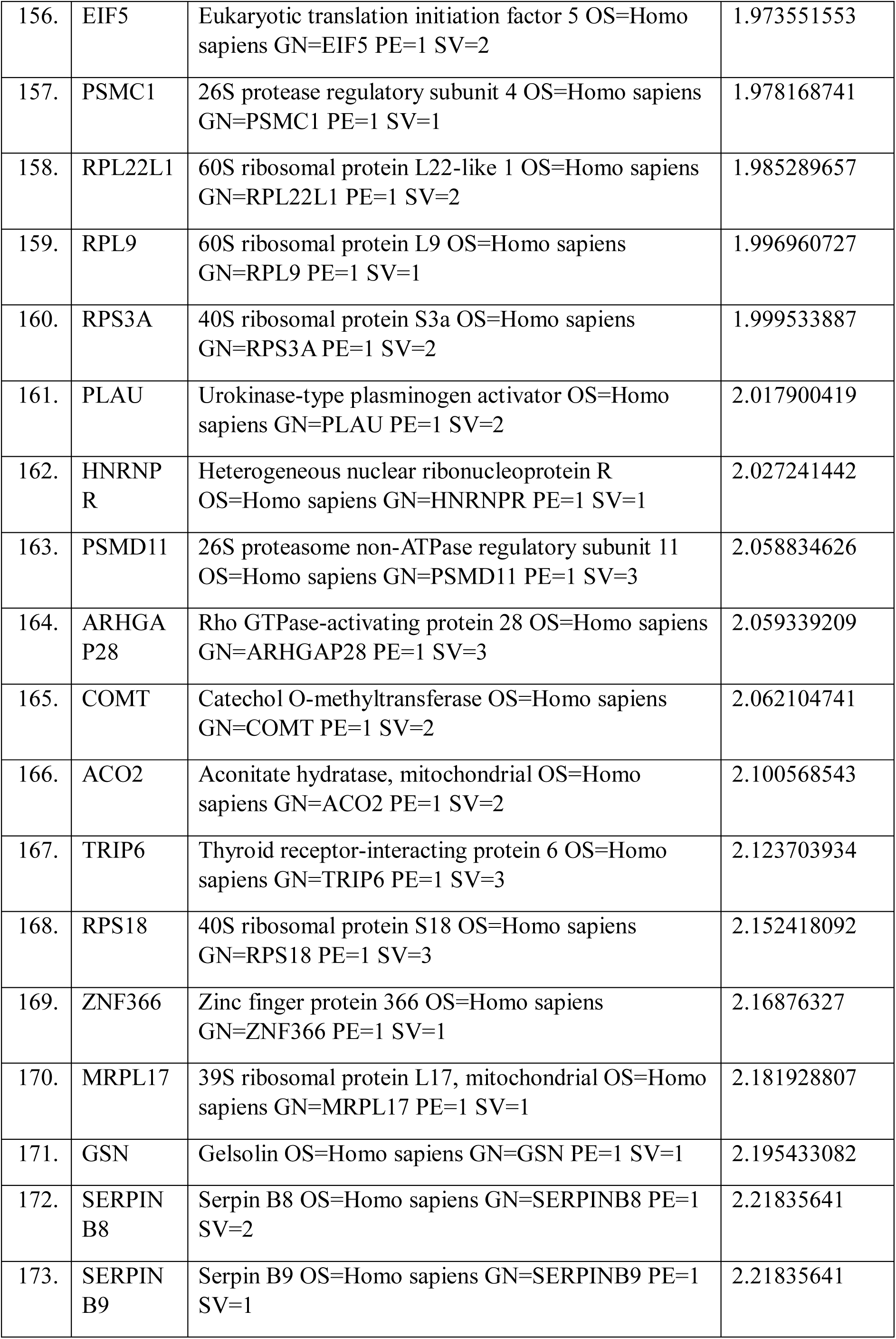
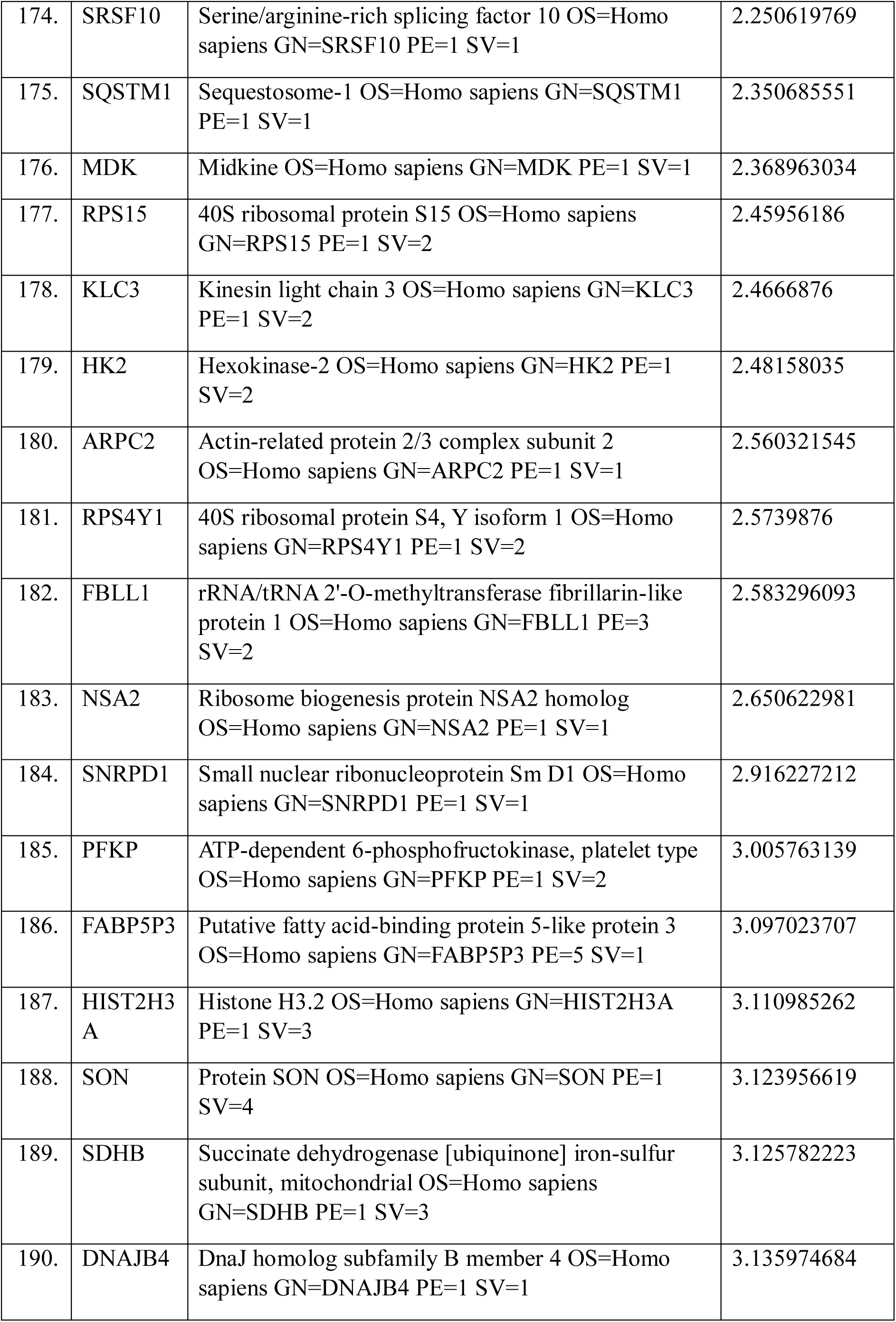
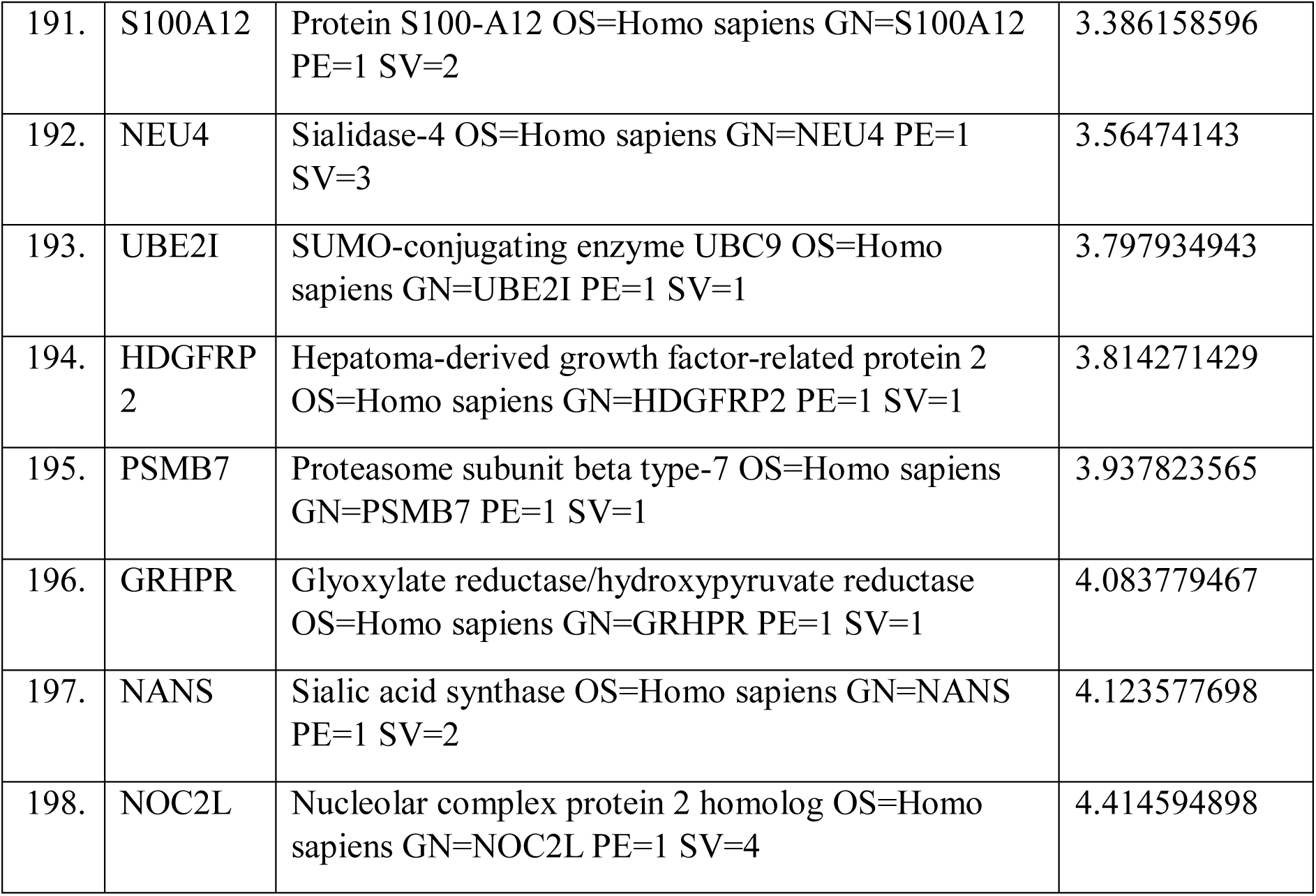
List of the genes found upregulated in Reg-R-HCT116 cells as compare to control in proteome data analysis keeping log2 Fold change > 1.

| S. No. | Gene Symbol | Description | Abundance Ratio (log2): |
| --- | --- | --- | --- |

|  |  |  | (T, Sample) /<br>(C, Control) |
| --- | --- | --- | --- |
| 1. | PPIB | Peptidyl-prolyl cis-trans isomerase B OS=Homo sapiens GN=PPIB PE=1 SV=2 | 1.015143355 |
| 2. | CHCHD2 | Coiled-coil-helix-coiled-coil-helix domain-containing protein 2 OS=Homo sapiens GN=CHCHD2 PE=1 SV=1 | 1.01587159 |
| 3. | CHCHD2P9 | Putative coiled-coil-helix-coiled-coil-helix domain-containing protein CHCHD2P9, mitochondrial OS=Homo sapiens GN=CHCHD2P9 PE=5 SV=1 | 1.01587159 |
| 4. | PGP | Glycerol-3-phosphate phosphatase OS=Homo sapiens GN=PGP PE=1 SV=1 | 1.017660188 |
| 5. | SMARCC1 | SWI/SNF complex subunit SMARCC1 OS=Homo sapiens GN=SMARCC1 PE=1 SV=3 | 1.018002063 |
| 6. | SMARCC2 | SWI/SNF complex subunit SMARCC2 OS=Homo sapiens GN=SMARCC2 PE=1 SV=1 | 1.018002063 |
| 7. | PHB2 | Prohibitin-2 OS=Homo sapiens GN=PHB2 PE=1 SV=2 | 1.018895226 |
| 8. | RPLP0P6 | 60S acidic ribosomal protein P0-like OS=Homo sapiens GN=RPLP0P6 PE=5 SV=1 | 1.020291148 |
| 9. | EXOSC6 | Exosome complex component MTR3 OS=Homo sapiens GN=EXOSC6 PE=1 SV=1 | 1.025281289 |
| 10. | HIST2H2AC | Histone H2A type 2-C OS=Homo sapiens GN=HIST2H2AC PE=1 SV=4 | 1.029180444 |
| 11. | GNG12 | Guanine nucleotide-binding protein G(I)/G(S)/G(O) subunit gamma-12 OS=Homo sapiens GN=GNG12 PE=1 SV=3 | 1.031926812 |
| 12. | GTPBP4 | Nucleolar GTP-binding protein 1 OS=Homo sapiens GN=GTPBP4 PE=1 SV=3 | 1.034933202 |
| 13. | HIST2H2AA3 | Histone H2A type 2-A OS=Homo sapiens GN=HIST2H2AA3 PE=1 SV=3 | 1.035421837 |
| 14. | RPS25 | 40S ribosomal protein S25 OS=Homo sapiens GN=RPS25 PE=1 SV=1 | 1.03732458 |
| 15. | VDAC2 | Voltage-dependent anion-selective channel protein 2 OS=Homo sapiens GN=VDAC2 PE=1 SV=2 | 1.039276112 |
| 16. | SF3A3 | Splicing factor 3A subunit 3 OS=Homo sapiens GN=SF3A3 PE=1 SV=1 | 1.046279347 |
| 17. | U2AF1 | Splicing factor U2AF 35 kDa subunit OS=Homo sapiens GN=U2AF1 PE=1 SV=3 | 1.049465511 |
| 18. | RPL35 | 60S ribosomal protein L35 OS=Homo sapiens GN=RPL35 PE=1 SV=2 | 1.049875334 |
| 19. | RRP15 | RRP15-like protein OS=Homo sapiens GN=RRP15 PE=1 SV=2 | 1.055350625 |
| 20. | TUBB6 | Tubulin beta-6 chain OS=Homo sapiens GN=TUBB6 PE=1 SV=1 | 1.055452465 |
| 21. | LOC260334 | Tubulin beta-8 chain-like protein LOC260334 OS=Homo sapiens PE=1 SV=1 | 1.055452465 |
| 22. | CNDP2 | Cytosolic non-specific dipeptidase OS=Homo sapiens GN=CNDP2 PE=1 SV=2 | 1.059493422 |
| 23. | CAPRIN1 | Caprin-1 OS=Homo sapiens GN=CAPRIN1 PE=1 SV=2 | 1.071606072 |
| 24. | COTL1 | Coactosin-like protein OS=Homo sapiens GN=COTL1 PE=1 SV=3 | 1.079666882 |
| 25. | RPL21 | 60S ribosomal protein L21 OS=Homo sapiens GN=RPL21 PE=1 SV=2 | 1.084097787 |
| 26. | GAR1 | H/ACA ribonucleoprotein complex subunit 1 OS=Homo sapiens GN=GAR1 PE=1 SV=1 | 1.092123927 |
| 27. | GCSH | Glycine cleavage system H protein, mitochondrial OS=Homo sapiens GN=GCSH PE=1 SV=2 | 1.092936905 |
| 28. | SRPRB | Signal recognition particle receptor subunit beta OS=Homo sapiens GN=SRPRB PE=1 SV=3 | 1.094182387 |
| 29. | HIST3H3 | Histone H3.1t OS=Homo sapiens GN=HIST3H3 PE=1 SV=3 | 1.0991876 |
| 30. | S100A4 | Protein S100-A4 OS=Homo sapiens GN=S100A4 PE=1 SV=1 | 1.103293524 |
| 31. | EWSR1 | RNA-binding protein EWS OS=Homo sapiens GN=EWSR1 PE=1 SV=1 | 1.104032604 |
| 32. | OR8K3 | Olfactory receptor 8K3 OS=Homo sapiens GN=OR8K3 PE=3 SV=1 | 1.108674031 |
| 33. | SP100 | Nuclear autoantigen Sp-100 OS=Homo sapiens GN=SP100 PE=1 SV=3 | 1.109666294 |
| 34. | RPL34 | 60S ribosomal protein L34 OS=Homo sapiens GN=RPL34 PE=1 SV=3 | 1.122397204 |
| 35. | GAPVD1 | GTPase-activating protein and VPS9 domain-containing protein 1 OS=Homo sapiens GN=GAPVD1 PE=1 SV=2 | 1.142072812 |
| 36. | ABI1 | Abl interactor 1 OS=Homo sapiens GN=ABI1 PE=1 SV=4 | 1.148783757 |
| 37. | EBNA1BP2 | Probable rRNA-processing protein EBP2 OS=Homo sapiens GN=EBNA1BP2 PE=1 SV=2 | 1.150048814 |
| 38. | POLR1C | DNA-directed RNA polymerases I and III subunit RPAC1 OS=Homo sapiens GN=POLR1C PE=1 SV=1 | 1.150649526 |
| 39. | SUCLG2 | Succinate--CoA ligase [GDP-forming] subunit beta, mitochondrial OS=Homo sapiens GN=SUCLG2 PE=1 SV=2 | 1.15439265 |
| 40. | HIST1H2BB | Histone H2B type 1-B OS=Homo sapiens GN=HIST1H2BB PE=1 SV=2 | 1.168783248 |
| 41. | SKP1 | S-phase kinase-associated protein 1 OS=Homo sapiens GN=SKP1 PE=1 SV=2 | 1.172885413 |
| 42. | HADHA | Trifunctional enzyme subunit alpha, mitochondrial OS=Homo sapiens GN=HADHA PE=1 SV=2 | 1.178121678 |
| 43. | ALB | Serum albumin OS=Homo sapiens GN=ALB PE=1 SV=2 | 1.182589248 |
| 44. | HNRNPL | Heterogeneous nuclear ribonucleoprotein L OS=Homo sapiens GN=HNRNPL PE=1 SV=2 | 1.197044029 |
| 45. | HBB | Hemoglobin subunit beta OS=Homo sapiens GN=HBB PE=1 SV=2 | 1.202708701 |
| 46. | HBD | Hemoglobin subunit delta OS=Homo sapiens GN=HBD PE=1 SV=2 | 1.202708701 |
| 47. | EIF3D | Eukaryotic translation initiation factor 3 subunit D OS=Homo sapiens GN=EIF3D PE=1 SV=1 | 1.204528225 |
| 48. | TUBB1 | Tubulin beta-1 chain OS=Homo sapiens GN=TUBB1 PE=1 SV=1 | 1.207313799 |
| 49. | RPS24 | 40S ribosomal protein S24 OS=Homo sapiens GN=RPS24 PE=1 SV=1 | 1.216610517 |
| 50. | RPE | Ribulose-phosphate 3-epimerase OS=Homo sapiens GN=RPE PE=1 SV=1 | 1.216730543 |
| 51. | RPEL1 | Ribulose-phosphate 3-epimerase-like protein 1 OS=Homo sapiens GN=RPEL1 PE=2 SV=1 | 1.216730543 |
| 52. | REXO2 | Oligoribonuclease, mitochondrial OS=Homo sapiens<br>GN=REXO2 PE=1 SV=3 | 1.217796182 |
| 53. | HIST2H2<br>BD | Putative histone H2B type 2-D OS=Homo sapiens<br>GN=HIST2H2BD PE=5 SV=3 | 1.21795814 |
| 54. | HIST2H2<br>BC | Putative histone H2B type 2-C OS=Homo sapiens<br>GN=HIST2H2BC PE=5 SV=3 | 1.21795814 |
| 55. | BUD31 | Protein BUD31 homolog OS=Homo sapiens<br>GN=BUD31 PE=1 SV=2 | 1.219725409 |
| 56. | RRP12 | RRP12-like protein OS=Homo sapiens GN=RRP12<br>PE=1 SV=2 | 1.223961857 |
| 57. | PCCA | Propionyl-CoA carboxylase alpha chain,<br>mitochondrial OS=Homo sapiens GN=PCCA PE=1<br>SV=4 | 1.227903584 |
| 58. | RPS8 | 40S ribosomal protein S8 OS=Homo sapiens<br>GN=RPS8 PE=1 SV=2 | 1.228807043 |
| 59. | GOT1 | Aspartate aminotransferase, cytoplasmic OS=Homo<br>sapiens GN=GOT1 PE=1 SV=3 | 1.232797866 |
| 60. | RPL30 | 60S ribosomal protein L30 OS=Homo sapiens<br>GN=RPL30 PE=1 SV=2 | 1.236128713 |
| 61. | RPS16 | 40S ribosomal protein S16 OS=Homo sapiens<br>GN=RPS16 PE=1 SV=2 | 1.246051709 |
| 62. | ANP32B | Acidic leucine-rich nuclear phosphoprotein 32 family<br>member B OS=Homo sapiens GN=ANP32B PE=1<br>SV=1 | 1.25162403 |
| 63. | HIST3H2<br>BB | Histone H2B type 3-B OS=Homo sapiens<br>GN=HIST3H2BB PE=1 SV=3 | 1.252625885 |
| 64. | RPS4X | 40S ribosomal protein S4, X isoform OS=Homo<br>sapiens GN=RPS4X PE=1 SV=2 | 1.253116279 |
| 65. | PATL1 | Protein PAT1 homolog 1 OS=Homo sapiens<br>GN=PATL1 PE=1 SV=2 | 1.262129871 |
| 66. | NFKB2 | Nuclear factor NF-kappa-B p100 subunit OS=Homo<br>sapiens GN=NFKB2 PE=1 SV=4 | 1.265003479 |
| 67. | SORD | Sorbitol dehydrogenase OS=Homo sapiens<br>GN=SORD PE=1 SV=4 | 1.269274979 |
| 68. | PPIH | Peptidyl-prolyl cis-trans isomerase H OS=Homo<br>sapiens GN=PPIH PE=1 SV=1 | 1.272592339 |
| 69. | CAT | Catalase OS=Homo sapiens GN=CAT PE=1 SV=3 | 1.276546937 |
| 70. | NUBP2 | Cytosolic Fe-S cluster assembly factor NUBP2<br>OS=Homo sapiens GN=NUBP2 PE=1 SV=1 | 1.283168519 |
| 71. | HBE1 | Hemoglobin subunit epsilon OS=Homo sapiens<br>GN=HBE1 PE=1 SV=2 | 1.288771123 |
| 72. | HBG1 | Hemoglobin subunit gamma-1 OS=Homo sapiens<br>GN=HBG1 PE=1 SV=2 | 1.288771123 |
| 73. | HBG2 | Hemoglobin subunit gamma-2 OS=Homo sapiens<br>GN=HBG2 PE=1 SV=2 | 1.288771123 |
| 74. | HMGN5 | High mobility group nucleosome-binding domain-containing protein 5 OS=Homo sapiens<br>GN=HMGN5 PE=1 SV=1 | 1.299779794 |
| 75. | SBDS | Ribosome maturation protein SBDS OS=Homo<br>sapiens GN=SBDS PE=1 SV=4 | 1.307326727 |
| 76. | ARPC5 | Actin-related protein 2/3 complex subunit 5<br>OS=Homo sapiens GN=ARPC5 PE=1 SV=3 | 1.310346621 |
| 77. | RPL32 | 60S ribosomal protein L32 OS=Homo sapiens<br>GN=RPL32 PE=1 SV=2 | 1.326136469 |
| 78. | DDX21 | Nucleolar RNA helicase 2 OS=Homo sapiens<br>GN=DDX21 PE=1 SV=5 | 1.333571823 |
| 79. | MCM6 | DNA replication licensing factor MCM6 OS=Homo<br>sapiens GN=MCM6 PE=1 SV=1 | 1.341489115 |
| 80. | POMP | Proteasome maturation protein OS=Homo sapiens<br>GN=POMP PE=1 SV=1 | 1.343458676 |
| 81. | PCGF2 | Polycomb group RING finger protein 2 OS=Homo<br>sapiens GN=PCGF2 PE=1 SV=1 | 1.349231358 |
| 82. | BMI1 | Polycomb complex protein BMI-1 OS=Homo<br>sapiens GN=BMI1 PE=1 SV=2 | 1.349231358 |
| 83. | GPI | Glucose-6-phosphate isomerase OS=Homo sapiens<br>GN=GPI PE=1 SV=4 | 1.354405342 |
| 84. | UTP18 | U3 small nucleolar RNA-associated protein 18<br>homolog OS=Homo sapiens GN=UTP18 PE=1<br>SV=3 | 1.357261197 |
| 85. | ERO1A | ERO1-like protein alpha OS=Homo sapiens<br>GN=ERO1A PE=1 SV=2 | 1.360013037 |
| 86. | RPL10 | 60S ribosomal protein L10 OS=Homo sapiens<br>GN=RPL10 PE=1 SV=4 | 1.369680236 |
| 87. | P4HA1 | Prolyl 4-hydroxylase subunit alpha-1 OS=Homo sapiens<br>GN=P4HA1 PE=1 SV=2 | 1.370239465 |
| 88. | HSD17B10 | 3-hydroxyacyl-CoA dehydrogenase type-2<br>OS=Homo sapiens GN=HSD17B10 PE=1 SV=3 | 1.373177094 |
| 89. | RPS6 | 40S ribosomal protein S6 OS=Homo sapiens<br>GN=RPS6 PE=1 SV=1 | 1.376262868 |
| 90. | HIST1H2AB | Histone H2A type 1-B/E OS=Homo sapiens<br>GN=HIST1H2AB PE=1 SV=2 | 1.383150712 |
| 91. | PAIP1 | Polyadenylate-binding protein-interacting protein 1<br>OS=Homo sapiens GN=PAIP1 PE=1 SV=1 | 1.395902514 |
| 92. | MFAP1 | Microfibrillar-associated protein 1 OS=Homo sapiens<br>GN=MFAP1 PE=1 SV=2 | 1.420454361 |
| 93. | PDHB | Pyruvate dehydrogenase E1 component subunit beta,<br>mitochondrial OS=Homo sapiens GN=PDHB PE=1<br>SV=3 | 1.42306316 |
| 94. | H2BFS | Histone H2B type F-S OS=Homo sapiens<br>GN=H2BFS PE=1 SV=2 | 1.424081072 |
| 95. | ERP29 | Endoplasmic reticulum resident protein 29<br>OS=Homo sapiens GN=ERP29 PE=1 SV=4 | 1.430202419 |
| 96. | COLGALT1 | Procollagen galactosyltransferase 1 OS=Homo sapiens<br>GN=COLGALT1 PE=1 SV=1 | 1.432857317 |
| 97. | CHORDC1 | Cysteine and histidine-rich domain-containing<br>protein 1 OS=Homo sapiens GN=CHORDC1 PE=1<br>SV=2 | 1.43617935 |
| 98. | GTF2I | General transcription factor II-I OS=Homo sapiens<br>GN=GTF2I PE=1 SV=2 | 1.442494141 |
| 99. | RPS4Y2 | 40S ribosomal protein S4, Y isoform 2 OS=Homo sapiens<br>GN=RPS4Y2 PE=2 SV=3 | 1.451973893 |
| 100. | CCDC137 | Coiled-coil domain-containing protein 137<br>OS=Homo sapiens GN=CCDC137 PE=1 SV=1 | 1.45522728 |
| 101. | PDE4A | cAMP-specific 3',5'-cyclic phosphodiesterase 4A<br>OS=Homo sapiens GN=PDE4A PE=1 SV=3 | 1.463446745 |
| 102. | RPS11 | 40S ribosomal protein S11 OS=Homo sapiens<br>GN=RPS11 PE=1 SV=3 | 1.487089037 |
| 103. | HIST1H2BK | Histone H2B type 1-K OS=Homo sapiens<br>GN=HIST1H2BK PE=1 SV=3 | 1.489361235 |
| 104. | RPS13 | 40S ribosomal protein S13 OS=Homo sapiens<br>GN=RPS13 PE=1 SV=2 | 1.508409829 |
| 105. | YBX3 | Y-box-binding protein 3 OS=Homo sapiens<br>GN=YBX3 PE=1 SV=4 | 1.509004085 |
| 106. | NMT1 | Glycylpeptide N-tetradecanoyltransferase 1<br>OS=Homo sapiens GN=NMT1 PE=1 SV=2 | 1.516830579 |
| 107. | DDT | D-dopachrome decarboxylase OS=Homo sapiens<br>GN=DDT PE=1 SV=3 | 1.519648719 |
| 108. | DDTL | D-dopachrome decarboxylase-like protein OS=Homo sapiens<br>GN=DDTL PE=2 SV=1 | 1.519648719 |
| 109. | NDRG1 | Protein NDRG1 OS=Homo sapiens GN=NDRG1<br>PE=1 SV=1 | 1.521907218 |
| 110. | HIST1H4A | Histone H4 OS=Homo sapiens GN=HIST1H4A<br>PE=1 SV=2 | 1.528796872 |
| 111. | H2AFY2 | Core histone macro-H2A.2 OS=Homo sapiens<br>GN=H2AFY2 PE=1 SV=3 | 1.528875115 |
| 112. | RPL37 | 60S ribosomal protein L37 OS=Homo sapiens<br>GN=RPL37 PE=1 SV=2 | 1.53689626 |
| 113. | RICTOR | Rapamycin-insensitive companion of mTOR<br>OS=Homo sapiens GN=RICTOR PE=1 SV=1 | 1.541840614 |
| 114. | OR4K3 | Olfactory receptor 4K3 OS=Homo sapiens<br>GN=OR4K3 PE=3 SV=3 | 1.542690067 |
| 115. | IPO11 | Importin-11 OS=Homo sapiens GN=IPO11 PE=1<br>SV=1 | 1.546087106 |
| 116. | PDCD11 | Protein RRP5 homolog OS=Homo sapiens<br>GN=PDCD11 PE=1 SV=3 | 1.559119364 |
| 117. | FUS | RNA-binding protein FUS OS=Homo sapiens<br>GN=FUS PE=1 SV=1 | 1.572416724 |
| 118. | H2AFY | Core histone macro-H2A.1 OS=Homo sapiens<br>GN=H2AFY PE=1 SV=4 | 1.58738215 |
| 119. | RPL10L | 60S ribosomal protein L10-like OS=Homo sapiens<br>GN=RPL10L PE=1 SV=3 | 1.598766034 |
| 120. | SLTM | SAFB-like transcription modulator OS=Homo sapiens<br>GN=SLTM PE=1 SV=2 | 1.616082283 |
| 121. | RCC1 | Regulator of chromosome condensation OS=Homo sapiens GN=RCC1 PE=1 SV=1 | 1.626552073 |
| 122. | MRPL24 | 39S ribosomal protein L24, mitochondrial OS=Homo sapiens GN=MRPL24 PE=1 SV=1 | 1.63104026 |
| 123. | WDR27 | WD repeat-containing protein 27 OS=Homo sapiens GN=WDR27 PE=1 SV=3 | 1.633661778 |
| 124. | EEF1E1 | Eukaryotic translation elongation factor 1 epsilon-1 OS=Homo sapiens GN=EEF1E1 PE=1 SV=1 | 1.668739102 |
| 125. | ACTA2 | Actin, aortic smooth muscle OS=Homo sapiens GN=ACTA2 PE=1 SV=1 | 1.670203686 |
| 126. | ACTG2 | Actin, gamma-enteric smooth muscle OS=Homo sapiens GN=ACTG2 PE=1 SV=1 | 1.670203686 |
| 127. | ALDH1A3 | Aldehyde dehydrogenase family 1 member A3 OS=Homo sapiens GN=ALDH1A3 PE=1 SV=2 | 1.677689208 |
| 128. | SRSF9 | Serine/arginine-rich splicing factor 9 OS=Homo sapiens GN=SRSF9 PE=1 SV=1 | 1.678607837 |
| 129. | OSBP | Oxysterol-binding protein 1 OS=Homo sapiens GN=OSBP PE=1 SV=1 | 1.689848004 |
| 130. | DDX52 | Probable ATP-dependent RNA helicase DDX52 OS=Homo sapiens GN=DDX52 PE=1 SV=3 | 1.71343775 |
| 131. | MAOA | Amine oxidase [flavin-containing] A OS=Homo sapiens GN=MAOA PE=1 SV=1 | 1.715054375 |
| 132. | MVP | Major vault protein OS=Homo sapiens GN=MVP PE=1 SV=4 | 1.717352088 |
| 133. | DBI | Acyl-CoA-binding protein OS=Homo sapiens GN=DBI PE=1 SV=2 | 1.71795689 |
| 134. | NUDT5 | ADP-sugar pyrophosphatase OS=Homo sapiens GN=NUDT5 PE=1 SV=1 | 1.719766977 |
| 135. | TFRC | Transferrin receptor protein 1 OS=Homo sapiens GN=TFRC PE=1 SV=2 | 1.720325801 |
| 136. | NOP14 | Nucleolar protein 14 OS=Homo sapiens GN=NOP14 PE=1 SV=3 | 1.728554661 |
| 137. | DNAJA1 | DnaJ homolog subfamily A member 1 OS=Homo sapiens GN=DNAJA1 PE=1 SV=2 | 1.738845127 |
| 138. | CDA | Cytidine deaminase OS=Homo sapiens GN=CDA PE=1 SV=2 | 1.75128345 |
| 139. | RPL35A | 60S ribosomal protein L35a OS=Homo sapiens<br>GN=RPL35A PE=1 SV=2 | 1.76412914 |
| 140. | PACSIN2 | Protein kinase C and casein kinase substrate in<br>neurons protein 2 OS=Homo sapiens GN=PACSIN2<br>PE=1 SV=2 | 1.767785828 |
| 141. | ANXA5 | Annexin A5 OS=Homo sapiens GN=ANXA5 PE=1<br>SV=2 | 1.783712343 |
| 142. | RPL27 | 60S ribosomal protein L27 OS=Homo sapiens<br>GN=RPL27 PE=1 SV=2 | 1.803447223 |
| 143. | HIST1H2<br>BA | Histone H2B type 1-A OS=Homo sapiens<br>GN=HIST1H2BA PE=1 SV=3 | 1.821809506 |
| 144. | PTRH2 | Peptidyl-tRNA hydrolase 2, mitochondrial<br>OS=Homo sapiens GN=PTRH2 PE=1 SV=1 | 1.839044663 |
| 145. | PSMC5 | 26S protease regulatory subunit 8 OS=Homo sapiens<br>GN=PSMC5 PE=1 SV=1 | 1.848819917 |
| 146. | ZNF638 | Zinc finger protein 638 OS=Homo sapiens<br>GN=ZNF638 PE=1 SV=2 | 1.859544064 |
| 147. | H3F3A | Histone H3.3 OS=Homo sapiens GN=H3F3A PE=1<br>SV=2 | 1.865691256 |
| 148. | EIF2AK2 | Interferon-induced, double-stranded RNA-activated<br>protein kinase OS=Homo sapiens GN=EIF2AK2<br>PE=1 SV=2 | 1.878089709 |
| 149. | TK1 | Thymidine kinase, cytosolic OS=Homo sapiens<br>GN=TK1 PE=1 SV=2 | 1.914572038 |
| 150. | TMPO | Lamina-associated polypeptide 2, isoform alpha<br>OS=Homo sapiens GN=TMPO PE=1 SV=2 | 1.931303121 |
| 151. | HKDC1 | Putative hexokinase HKDC1 OS=Homo sapiens<br>GN=HKDC1 PE=1 SV=3 | 1.935857377 |
| 152. | HK3 | Hexokinase-3 OS=Homo sapiens GN=HK3 PE=1<br>SV=2 | 1.935857377 |
| 153. | HK1 | Hexokinase-1 OS=Homo sapiens GN=HK1 PE=1<br>SV=3 | 1.935857377 |
| 154. | TTLL12 | Tubulin--tyrosine ligase-like protein 12 OS=Homo<br>sapiens GN=TTLL12 PE=1 SV=2 | 1.964848387 |
| 155. | CCDC13 | Coiled-coil domain-containing protein 13 OS=Homo<br>sapiens GN=CCDC13 PE=1 SV=2 | 1.973490273 |
| 156. | EIF5 | Eukaryotic translation initiation factor 5 OS=Homo sapiens GN=EIF5 PE=1 SV=2 | 1.973551553 |
| 157. | PSMC1 | 26S protease regulatory subunit 4 OS=Homo sapiens GN=PSMC1 PE=1 SV=1 | 1.978168741 |
| 158. | RPL22L1 | 60S ribosomal protein L22-like 1 OS=Homo sapiens GN=RPL22L1 PE=1 SV=2 | 1.985289657 |
| 159. | RPL9 | 60S ribosomal protein L9 OS=Homo sapiens GN=RPL9 PE=1 SV=1 | 1.996960727 |
| 160. | RPS3A | 40S ribosomal protein S3a OS=Homo sapiens GN=RPS3A PE=1 SV=2 | 1.999533887 |
| 161. | PLAU | Urokinase-type plasminogen activator OS=Homo sapiens GN=PLAU PE=1 SV=2 | 2.017900419 |
| 162. | HNRNP R | Heterogeneous nuclear ribonucleoprotein R OS=Homo sapiens GN=HNRNPR PE=1 SV=1 | 2.027241442 |
| 163. | PSMD11 | 26S proteasome non-ATPase regulatory subunit 11 OS=Homo sapiens GN=PSMD11 PE=1 SV=3 | 2.058834626 |
| 164. | ARHGA P28 | Rho GTPase-activating protein 28 OS=Homo sapiens GN=ARHGAP28 PE=1 SV=3 | 2.059339209 |
| 165. | COMT | Catechol O-methyltransferase OS=Homo sapiens GN=COMT PE=1 SV=2 | 2.062104741 |
| 166. | ACO2 | Aconitate hydratase, mitochondrial OS=Homo sapiens GN=ACO2 PE=1 SV=2 | 2.100568543 |
| 167. | TRIP6 | Thyroid receptor-interacting protein 6 OS=Homo sapiens GN=TRIP6 PE=1 SV=3 | 2.123703934 |
| 168. | RPS18 | 40S ribosomal protein S18 OS=Homo sapiens GN=RPS18 PE=1 SV=3 | 2.152418092 |
| 169. | ZNF366 | Zinc finger protein 366 OS=Homo sapiens GN=ZNF366 PE=1 SV=1 | 2.16876327 |
| 170. | MRPL17 | 39S ribosomal protein L17, mitochondrial OS=Homo sapiens GN=MRPL17 PE=1 SV=1 | 2.181928807 |
| 171. | GSN | Gelsolin OS=Homo sapiens GN=GSN PE=1 SV=1 | 2.195433082 |
| 172. | SERPIN B8 | Serpin B8 OS=Homo sapiens GN=SERPINB8 PE=1 SV=2 | 2.21835641 |
| 173. | SERPIN B9 | Serpin B9 OS=Homo sapiens GN=SERPINB9 PE=1 SV=1 | 2.21835641 |
| 174. | SRSF10 | Serine/arginine-rich splicing factor 10 OS=Homo sapiens GN=SRSF10 PE=1 SV=1 | 2.250619769 |
| 175. | SQSTM1 | Sequestosome-1 OS=Homo sapiens GN=SQSTM1 PE=1 SV=1 | 2.350685551 |
| 176. | MDK | Midkine OS=Homo sapiens GN=MDK PE=1 SV=1 | 2.368963034 |
| 177. | RPS15 | 40S ribosomal protein S15 OS=Homo sapiens GN=RPS15 PE=1 SV=2 | 2.45956186 |
| 178. | KLC3 | Kinesin light chain 3 OS=Homo sapiens GN=KLC3 PE=1 SV=2 | 2.4666876 |
| 179. | HK2 | Hexokinase-2 OS=Homo sapiens GN=HK2 PE=1 SV=2 | 2.48158035 |
| 180. | ARPC2 | Actin-related protein 2/3 complex subunit 2 OS=Homo sapiens GN=ARPC2 PE=1 SV=1 | 2.560321545 |
| 181. | RPS4Y1 | 40S ribosomal protein S4, Y isoform 1 OS=Homo sapiens GN=RPS4Y1 PE=1 SV=2 | 2.5739876 |
| 182. | FBLL1 | rRNA/tRNA 2'-O-methyltransferase fibrillarin-like protein 1 OS=Homo sapiens GN=FBLL1 PE=3 SV=2 | 2.583296093 |
| 183. | NSA2 | Ribosome biogenesis protein NSA2 homolog OS=Homo sapiens GN=NSA2 PE=1 SV=1 | 2.650622981 |
| 184. | SNRPD1 | Small nuclear ribonucleoprotein Sm D1 OS=Homo sapiens GN=SNRPD1 PE=1 SV=1 | 2.916227212 |
| 185. | PFKP | ATP-dependent 6-phosphofructokinase, platelet type OS=Homo sapiens GN=PFKP PE=1 SV=2 | 3.005763139 |
| 186. | FABP5P3 | Putative fatty acid-binding protein 5-like protein 3 OS=Homo sapiens GN=FABP5P3 PE=5 SV=1 | 3.097023707 |
| 187. | HIST2H3A | Histone H3.2 OS=Homo sapiens GN=HIST2H3A PE=1 SV=3 | 3.110985262 |
| 188. | SON | Protein SON OS=Homo sapiens GN=SON PE=1 SV=4 | 3.123956619 |
| 189. | SDHB | Succinate dehydrogenase [ubiquinone] iron-sulfur subunit, mitochondrial OS=Homo sapiens GN=SDHB PE=1 SV=3 | 3.125782223 |
| 190. | DNAJB4 | DnaJ homolog subfamily B member 4 OS=Homo sapiens GN=DNAJB4 PE=1 SV=1 | 3.135974684 |
| 191. | S100A12 | Protein S100-A12 OS=Homo sapiens GN=S100A12 PE=1 SV=2 | 3.386158596 |
| 192. | NEU4 | Sialidase-4 OS=Homo sapiens GN=NEU4 PE=1 SV=3 | 3.56474143 |
| 193. | UBE2I | SUMO-conjugating enzyme UBC9 OS=Homo sapiens GN=UBE2I PE=1 SV=1 | 3.797934943 |
| 194. | HDGFRP2 | Hepatoma-derived growth factor-related protein 2 OS=Homo sapiens GN=HDGFRP2 PE=1 SV=1 | 3.814271429 |
| 195. | PSMB7 | Proteasome subunit beta type-7 OS=Homo sapiens GN=PSMB7 PE=1 SV=1 | 3.937823565 |
| 196. | GRHPR | Glyoxylate reductase/hydroxypyruvate reductase OS=Homo sapiens GN=GRHPR PE=1 SV=1 | 4.083779467 |
| 197. | NANS | Sialic acid synthase OS=Homo sapiens GN=NANS PE=1 SV=2 | 4.123577698 |
| 198. | NOC2L | Nucleolar complex protein 2 homolog OS=Homo sapiens GN=NOC2L PE=1 SV=4 | 4.414594898 |

**Table 3:** List of the genes found downregulated in Reg-R-HCT116 cells as compare to control in proteome data analysis keeping log2 Fold change >-1.

| S. No. | Accession | Description | <u>Abundance Ratio (log2): (T, Sample) / (C, Control)</u> |
| --- | --- | --- | --- |
| 1. | EGFR | Epidermal growth factor receptor OS=Homo sapiens GN=EGFR PE=1 SV=2 | -4.39444187 |
| 2. | LRSAM1 | E3 ubiquitin-protein ligase LRSAM1 OS=Homo sapiens GN=LRSAM1 PE=1 SV=1 | -3.65710023 |
| 3. | PRPF3 | U4/U6 small nuclear ribonucleoprotein Prp3 OS=Homo sapiens GN=PRPF3 PE=1 SV=2 | -3.32283236 |
| 4. | EXOSC3 | Exosome complex component RRP40 OS=Homo sapiens GN=EXOSC3 PE=1 SV=3 | -2.94152692 |
| 5. | HMGN1 | Non-histone chromosomal protein HMG-14 OS=Homo sapiens GN=HMGN1 PE=1 SV=3 | -2.89575786 |
| 6. | TMA16 | Translation machinery-associated protein 16 OS=Homo sapiens GN=TMA16 PE=1 SV=2 | -2.78285811 |
| 7. | CYB5B | Cytochrome b5 type B OS=Homo sapiens GN=CYB5B PE=1 SV=2 | -2.36124711 |
| 8. | ZGRF1 | Protein ZGRF1 OS=Homo sapiens<br>GN=ZGRF1 PE=1 SV=3 | -2.35710332 |
| 9. | HSP90AA4P | Putative heat shock protein HSP 90-alpha A4<br>OS=Homo sapiens GN=HSP90AA4P PE=5<br>SV=1 | -2.34354435 |
| 10. | EPCAM | Epithelial cell adhesion molecule OS=Homo<br>sapiens GN=EPCAM PE=1 SV=2 | -2.32764078 |
| 11. | MBNL2 | Muscleblind-like protein 2 OS=Homo sapiens<br>GN=MBNL2 PE=1 SV=2 | -2.0794873 |
| 12. | EPB41L2 | Band 4.1-like protein 2 OS=Homo sapiens<br>GN=EPB41L2 PE=1 SV=1 | -2.07532883 |
| 13. | SAFB2 | Scaffold attachment factor B2 OS=Homo<br>sapiens GN=SAFB2 PE=1 SV=1 | -2.06314749 |
| 14. | PYM1 | Partner of Y14 and mago OS=Homo sapiens<br>GN=PYM1 PE=1 SV=1 | -1.96933242 |
| 15. | ADRM1 | Proteasomal ubiquitin receptor ADRM1<br>OS=Homo sapiens GN=ADRM1 PE=1 SV=2 | -1.8993794 |
| 16. | CANX | Calnexin OS=Homo sapiens GN=CANX<br>PE=1 SV=2 | -1.82963878 |
| 17. | PSME2 | Proteasome activator complex subunit 2<br>OS=Homo sapiens GN=PSME2 PE=1 SV=4 | -1.8288157 |
| 18. | FAM234B | Protein FAM234B OS=Homo sapiens<br>GN=FAM234B PE=1 SV=1 | -1.73311155 |
| 19. | DIS3 | Exosome complex exonuclease RRP44<br>OS=Homo sapiens GN=DIS3 PE=1 SV=2 | -1.55730442 |
| 20. | TIMM9 | Mitochondrial import inner membrane<br>translocase subunit Tim9 OS=Homo sapiens<br>GN=TIMM9 PE=1 SV=1 | -1.51621167 |
| 21. | PGAM5 | Serine/threonine-protein phosphatase PGAM5,<br>mitochondrial OS=Homo sapiens<br>GN=PGAM5 PE=1 SV=2 | -1.51312653 |
| 22. | NUCKS1 | Nuclear ubiquitous casein and cyclin-<br>dependent kinase substrate 1 OS=Homo<br>sapiens GN=NUCKS1 PE=1 SV=1 | -1.41178142 |
| 23. | CCAR1 | Cell division cycle and apoptosis regulator<br>protein 1 OS=Homo sapiens GN=CCAR1<br>PE=1 SV=2 | -1.40641669 |
| 24. | CLPTM1 | Cleft lip and palate transmembrane protein 1<br>OS=Homo sapiens GN=CLPTM1 PE=1 SV=1 | -1.37552017 |
| 25. | PAFAH1B3 | Platelet-activating factor acetylhydrolase IB<br>subunit gamma OS=Homo sapiens<br>GN=PAFAH1B3 PE=1 SV=1 | -1.34897617 |
| 26. | PDLIM1 | PDZ and LIM domain protein 1 OS=Homo<br>sapiens GN=PDLIM1 PE=1 SV=4 | -1.32372168 |
| 27. | YBX1 | Nuclease-sensitive element-binding protein 1<br>OS=Homo sapiens GN=YBX1 PE=1 SV=3 | -1.32029193 |
| 28. | KRT13 | Keratin, type I cytoskeletal 13 OS=Homo<br>sapiens GN=KRT13 PE=1 SV=4 | -1.25814494 |
| 29. | EIF3B | Eukaryotic translation initiation factor 3<br>subunit B OS=Homo sapiens GN=EIF3B<br>PE=1 SV=3 | -1.25750912 |
| 30. | PYGB | Glycogen phosphorylase, brain form<br>OS=Homo sapiens GN=PYGB PE=1 SV=5 | -1.24920029 |
| 31. | MBNL1 | Muscleblind-like protein 1 OS=Homo sapiens<br>GN=MBNL1 PE=1 SV=2 | -1.23190677 |
| 32. | CBX3 | Chromobox protein homolog 3 OS=Homo<br>sapiens GN=CBX3 PE=1 SV=4 | -1.23136009 |
| 33. | PFDN6 | Prefoldin subunit 6 OS=Homo sapiens<br>GN=PFDN6 PE=1 SV=1 | -1.21650905 |
| 34. | XRN2 | 5'-3' exoribonuclease 2 OS=Homo sapiens<br>GN=XRN2 PE=1 SV=1 | -1.20880472 |
| 35. | HMGN2 | Non-histone chromosomal protein HMG-17<br>OS=Homo sapiens GN=HMGN2 PE=1 SV=3 | -1.17073426 |
| 36. | C1QBP | Complement component 1 Q subcomponent-<br>binding protein, mitochondrial OS=Homo<br>sapiens GN=C1QBP PE=1 SV=1 | -1.13560446 |
| 37. | ZNF326 | DBIRD complex subunit ZNF326 OS=Homo<br>sapiens GN=ZNF326 PE=1 SV=2 | -1.11976928 |
| 38. | TAGLN3 | Transgelin-3 OS=Homo sapiens<br>GN=TAGLN3 PE=1 SV=2 | -1.11896065 |
| 39. | MKI67 | Proliferation marker protein Ki-67 OS=Homo<br>sapiens GN=MKI67 PE=1 SV=2 | -1.11589293 |
| 40. | HMGN4 | High mobility group nucleosome-binding domain-containing protein 4 OS=Homo sapiens GN=HMGN4 PE=1 SV=3 | -1.10004428 |
| 41. | USP39 | U4/U6.U5 tri-snRNP-associated protein 2 OS=Homo sapiens GN=USP39 PE=1 SV=2 | -1.09254256 |
| 42. | PPP1R14B | Protein phosphatase 1 regulatory subunit 14B OS=Homo sapiens GN=PPP1R14B PE=1 SV=3 | -1.07737359 |
| 43. | UQCRH | Cytochrome b-c1 complex subunit 6, mitochondrial OS=Homo sapiens GN=UQCRH PE=1 SV=2 | -1.06268956 |
| 44. | PCK2 | Phosphoenolpyruvate carboxykinase [GTP], mitochondrial OS=Homo sapiens GN=PCK2 PE=1 SV=3 | -1.04139658 |
| 45. | EIF5B | Eukaryotic translation initiation factor 5B OS=Homo sapiens GN=EIF5B PE=1 SV=4 | -1.03788263 |
| 46. | FKBP3 | Peptidyl-prolyl cis-trans isomerase FKBP3 OS=Homo sapiens GN=FKBP3 PE=1 SV=1 | -1.01627714 |
| 47. | RBMX | RNA-binding motif protein, X chromosome OS=Homo sapiens GN=RBMX PE=1 SV=3 | -1.00071537 |

**Table 4:** Genes specifically found only in Reg-R-HCT116 cell line in the proteome data.

| S. No. | Gene Symbol | Gene Name |
| --- | --- | --- |
| 1 | DKK1 | Dickkopf-related protein 1 OS=Homo sapiens GN=DKK1 PE=1 SV=1 |
| 2 | PYCR2 | Pyrroline-5-carboxylate reductase 2 OS=Homo sapiens GN=PYCR2 PE=1 SV=1 |
| 3 | DBNL | Drebrin-like protein OS=Homo sapiens GN=DBNL PE=1 SV=1 |
| 4 | CSE1L | Exportin-2 OS=Homo sapiens GN=CSE1L PE=1 SV=3 |
| 5 | IK | Protein Red OS=Homo sapiens GN=IK PE=1 SV=3 |
| 6 | PUF60 | Poly(U)-binding-splicing factor PUF60 OS=Homo sapiens GN=PUF60 PE=1 SV=1 |
| 7 | METAP2 | Methionine aminopeptidase 2 OS=Homo sapiens GN=METAP2 PE=1 SV=1 |
| 8 | KRI1 | Protein KRI1 homolog OS=Homo sapiens GN=KRI1 PE=1 SV=3 |
| 9 | NUP107 | Nuclear pore complex protein Nup107 OS=Homo sapiens<br>GN=NUP107 PE=1 SV=1 |
| 10 | YARS2 | Tyrosine--tRNA ligase, mitochondrial OS=Homo sapiens<br>GN=YARS2 PE=1 SV=2 |
| 11 | FMR1 | Synaptic functional regulator FMR1 OS=Homo sapiens GN=FMR1<br>PE=1 SV=1 |
| 12 | MRPL55 | 39S ribosomal protein L55, mitochondrial OS=Homo sapiens<br>GN=MRPL55 PE=1 SV=1 |
| 13 | SDF2L1 | Stromal cell-derived factor 2-like protein 1 OS=Homo sapiens<br>GN=SDF2L1 PE=1 SV=2 |
| 14 | HLA-A | HLA class I histocompatibility antigen, A-11 alpha chain OS=Homo sapiens<br>GN=HLA-A PE=1 SV=1 |
| 15 | SRP54 | Signal recognition particle 54 kDa protein OS=Homo sapiens<br>GN=SRP54 PE=1 SV=1 |
| 16 | RPAP3 | RNA polymerase II-associated protein 3 OS=Homo sapiens<br>GN=RPAP3 PE=1 SV=2 |
| 17 | SNCG | Gamma-synuclein OS=Homo sapiens GN=SNCG PE=1 SV=2 |
| 18 | RFC4 | Replication factor C subunit 4 OS=Homo sapiens GN=RFC4 PE=1<br>SV=2 |
| 19 | THOP1 | Thimet oligopeptidase OS=Homo sapiens GN=THOP1 PE=1 SV=2 |
| 20 | TAF15 | TATA-binding protein-associated factor 2N OS=Homo sapiens<br>GN=TAF15 PE=1 SV=1 |
| 21 | RAP1B | Ras-related protein Rap-1b OS=Homo sapiens GN=RAP1B PE=1<br>SV=1 |
| 22 | RAP1B | Ras-related protein Rap-1b-like protein OS=Homo sapiens PE=2<br>SV=1 |
| 23 | RAP1A | Ras-related protein Rap-1A OS=Homo sapiens GN=RAP1A PE=1<br>SV=1 |
| 24 | EIF4EBP1 | Eukaryotic translation initiation factor 4E-binding protein 1<br>OS=Homo sapiens GN=EIF4EBP1 PE=1 SV=3 |
| 25 | HECTD1 | E3 ubiquitin-protein ligase HECTD1 OS=Homo sapiens<br>GN=HECTD1 PE=1 SV=3 |
| 26 | CD2AP | CD2-associated protein OS=Homo sapiens GN=CD2AP PE=1 SV=1 |
| 27 | QARS | Glutamine--tRNA ligase OS=Homo sapiens GN=QARS PE=1 SV=1 |
| 28 | UQCRC2 | Cytochrome b-c1 complex subunit 2, mitochondrial OS=Homo sapiens GN=UQCRC2 PE=1 SV=3 |
| 29 | UBQLN2 | Ubiquilin-2 OS=Homo sapiens GN=UBQLN2 PE=1 SV=2 |
| 30 | NQO1 | NAD(P)H dehydrogenase [quinone] 1 OS=Homo sapiens GN=NQO1 PE=1 SV=1 |
| 31 | RGPD2 | RANBP2-like and GRIP domain-containing protein 2 OS=Homo sapiens GN=RGPD2 PE=2 SV=1 |
| 32 | RGPD3 | RanBP2-like and GRIP domain-containing protein 3 OS=Homo sapiens GN=RGPD3 PE=3 SV=2 |
| 33 | RGPD5 | RANBP2-like and GRIP domain-containing protein 5/6 OS=Homo sapiens GN=RGPD5 PE=1 SV=3 |
| 34 | RGPD1 | RANBP2-like and GRIP domain-containing protein 1 OS=Homo sapiens GN=RGPD1 PE=2 SV=1 |
| 35 | RANBP2 | E3 SUMO-protein ligase RanBP2 OS=Homo sapiens GN=RANBP2 PE=1 SV=2 |
| 36 | RGPD8 | RANBP2-like and GRIP domain-containing protein 8 OS=Homo sapiens GN=RGPD8 PE=1 SV=2 |
| 37 | EMG1 | Ribosomal RNA small subunit methyltransferase NEP1 OS=Homo sapiens GN=EMG1 PE=1 SV=4 |
| 38 | BID | BH3-interacting domain death agonist OS=Homo sapiens GN=BID PE=1 SV=1 |
| 39 | VDAC3 | Voltage-dependent anion-selective channel protein 3 OS=Homo sapiens GN=VDAC3 PE=1 SV=1 |
| 40 | CHMP1B | Charged multivesicular body protein 1b OS=Homo sapiens GN=CHMP1B PE=1 SV=1 |
| 41 | LARP4 | La-related protein 4 OS=Homo sapiens GN=LARP4 PE=1 SV=3 |
| 42 | TF | Serotransferrin OS=Homo sapiens GN=TF PE=1 SV=3 |
| 43 | AKR1A1 | Alcohol dehydrogenase [NADP(+)] OS=Homo sapiens GN=AKR1A1 PE=1 SV=3 |
| 44 | LAP3 | Cytosol aminopeptidase OS=Homo sapiens GN=LAP3 PE=1 SV=3 |
| 45 | PXN | Paxillin OS=Homo sapiens GN=PXN PE=1 SV=3 |
| 46 | MRPL41 | 39S ribosomal protein L41, mitochondrial OS=Homo sapiens GN=MRPL41 PE=1 SV=1 |
| 47 | RBM25 | RNA-binding protein 25 OS=Homo sapiens GN=RBM25 PE=1 SV=3 |
| 48 | UBE2N | Ubiquitin-conjugating enzyme E2 N OS=Homo sapiens<br>GN=UBE2N PE=1 SV=1 |
| 49 | UQCRQ | Cytochrome b-c1 complex subunit 8 OS=Homo sapiens<br>GN=UQCRQ PE=1 SV=4 |
| 50 | TBCB | Tubulin-folding cofactor B OS=Homo sapiens GN=TBCB PE=1<br>SV=2 |
| 51 | PHF6 | PHD finger protein 6 OS=Homo sapiens GN=PHF6 PE=1 SV=1 |
| 52 | CCDC12 | Coiled-coil domain-containing protein 12 OS=Homo sapiens<br>GN=CCDC12 PE=1 SV=1 |
| 53 | KIF5A | Kinesin heavy chain isoform 5A OS=Homo sapiens GN=KIF5A<br>PE=1 SV=2 |
| 54 | KIF5C | Kinesin heavy chain isoform 5C OS=Homo sapiens GN=KIF5C<br>PE=1 SV=1 |
| 55 | TRIP10 | Cdc42-interacting protein 4 OS=Homo sapiens GN=TRIP10 PE=1<br>SV=3 |
| 56 | AKAP8 | A-kinase anchor protein 8 OS=Homo sapiens GN=AKAP8 PE=1<br>SV=1 |
| 57 | GLG1 | Golgi apparatus protein 1 OS=Homo sapiens GN=GLG1 PE=1<br>SV=2 |
| 58 | TIAL1 | Nucleolysin TIAR OS=Homo sapiens GN=TIAL1 PE=1 SV=1 |
| 59 | NLN | Neurolysin, mitochondrial OS=Homo sapiens GN=NLN PE=1<br>SV=1 |
| 60 | MAGEB2 | Melanoma-associated antigen B2 OS=Homo sapiens GN=MAGEB2<br>PE=1 SV=3 |
| 61 | CDC73 | Parafibromin OS=Homo sapiens GN=CDC73 PE=1 SV=1 |
| 62 | CRTC2 | CREB-regulated transcription coactivator 2 OS=Homo sapiens<br>GN=CRTC2 PE=1 SV=2 |
| 63 | TTC4 | Tetratricopeptide repeat protein 4 OS=Homo sapiens GN=TTC4<br>PE=1 SV=3 |
| 64 | ALDH1B1 | Aldehyde dehydrogenase X, mitochondrial OS=Homo sapiens<br>GN=ALDH1B1 PE=1 SV=3 |
| 65 | DKK2 | Dickkopf-related protein 2 OS=Homo sapiens GN=DKK2 PE=2<br>SV=1 |
| 66 | PLOD1 | Procollagen-lysine,2-oxoglutarate 5-dioxygenase 1 OS=Homo<br>sapiens GN=PLOD1 PE=1 SV=2 |
| 67 | IWS1 | Protein IWS1 homolog OS=Homo sapiens GN=IWS1 PE=1 SV=2 |
| 68 | MRPS33 | 28S ribosomal protein S33, mitochondrial OS=Homo sapiens GN=MRPS33 PE=1 SV=1 |
| 69 | TOR1AIP2 | Torsin-1A-interacting protein 2 OS=Homo sapiens GN=TOR1AIP2 PE=1 SV=1 |
| 70 | TJP1 | Tight junction protein ZO-1 OS=Homo sapiens GN=TJP1 PE=1 SV=3 |
| 71 | OLA1 | Obg-like ATPase 1 OS=Homo sapiens GN=OLA1 PE=1 SV=2 |

**Table 5:**
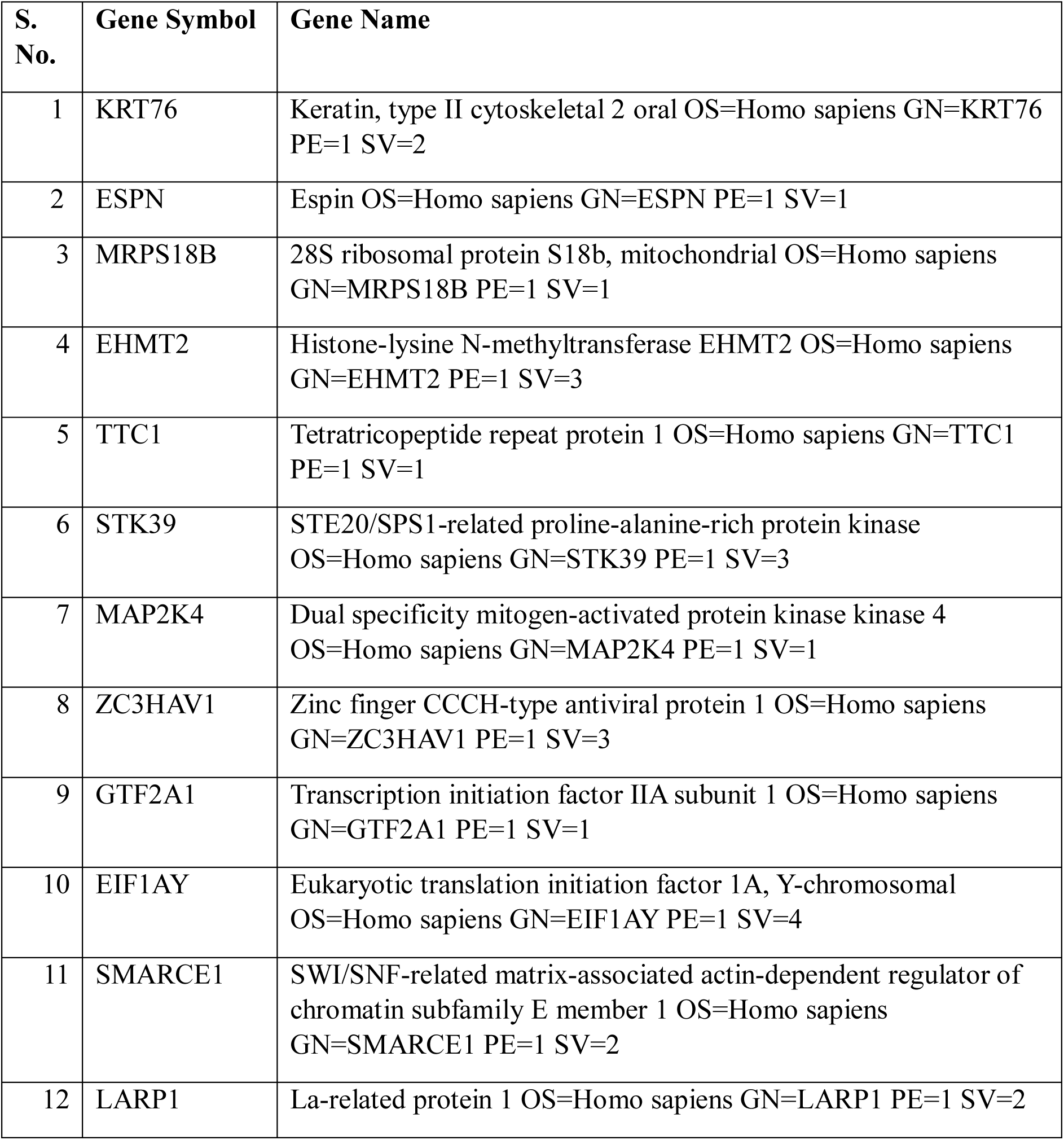
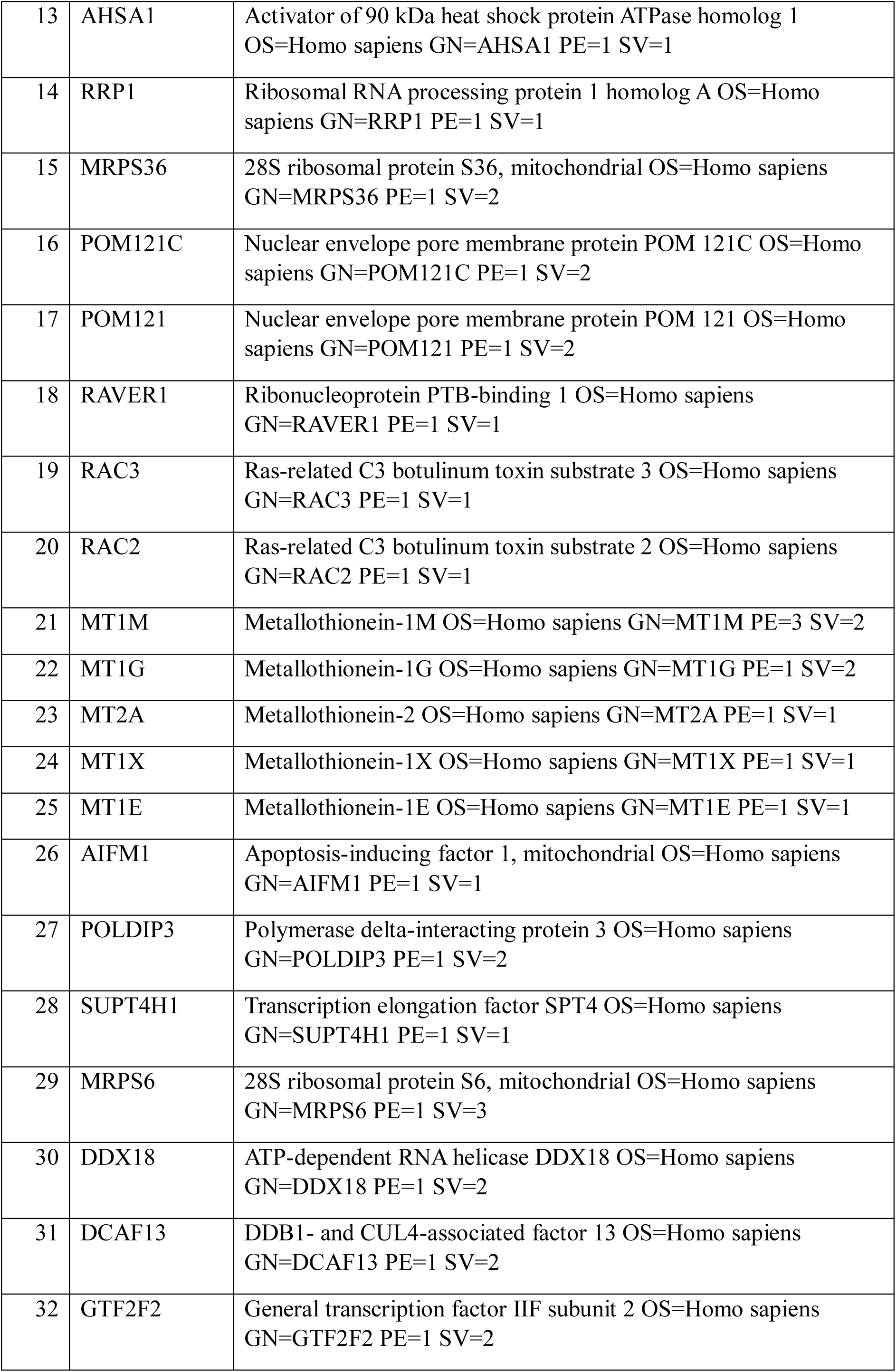
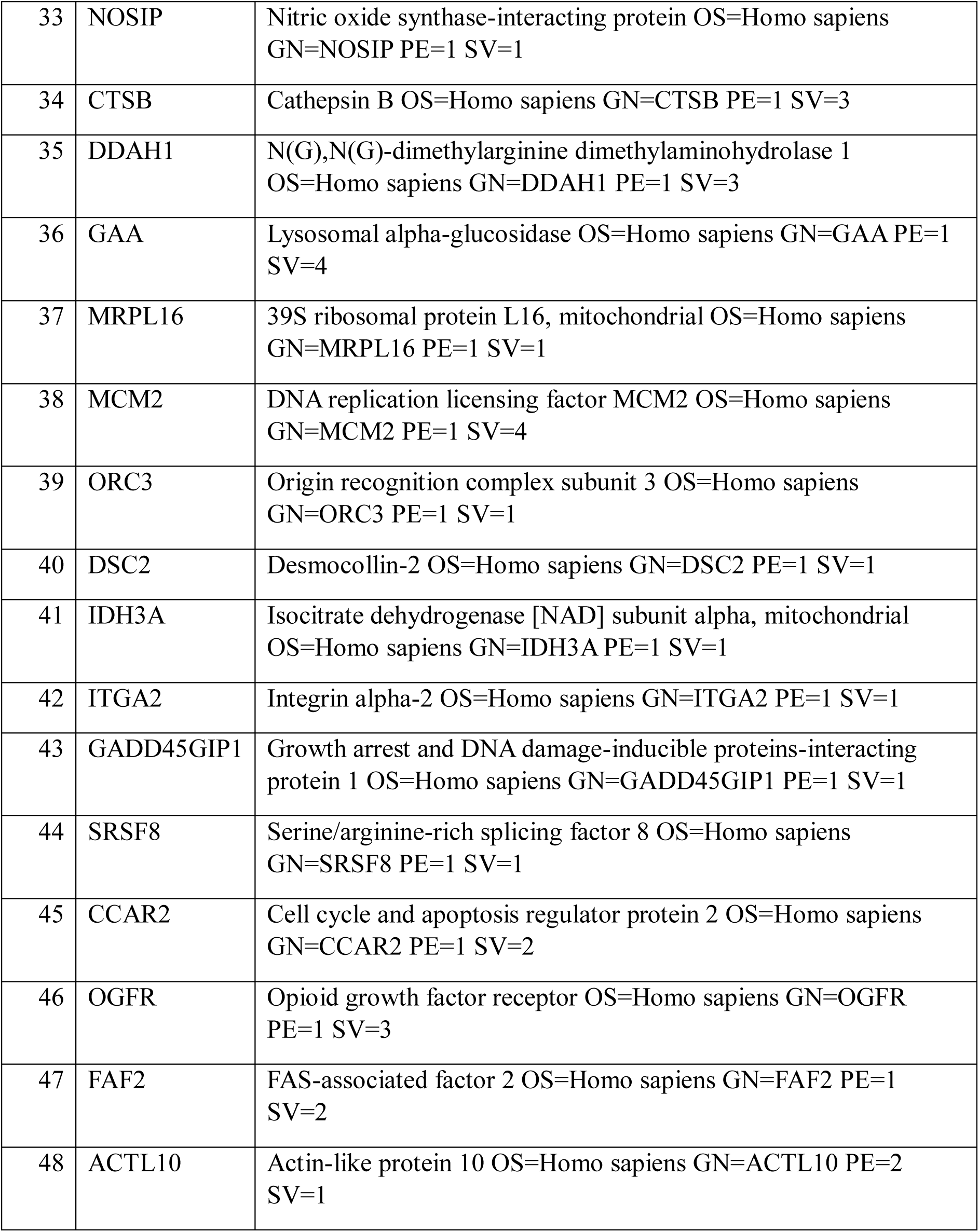
Genes specifically found only in Control-HCT116 cell line in the proteome data.

| S. No. | Gene Symbol | Gene Name |
| --- | --- | --- |
| 1 | KRT76 | Keratin, type II cytoskeletal 2 oral OS=Homo sapiens GN=KRT76 PE=1 SV=2 |
| 2 | ESPN | Espin OS=Homo sapiens GN=ESPN PE=1 SV=1 |
| 3 | MRPS18B | 28S ribosomal protein S18b, mitochondrial OS=Homo sapiens GN=MRPS18B PE=1 SV=1 |
| 4 | EHMT2 | Histone-lysine N-methyltransferase EHMT2 OS=Homo sapiens GN=EHMT2 PE=1 SV=3 |
| 5 | TTC1 | Tetratricopeptide repeat protein 1 OS=Homo sapiens GN=TTC1 PE=1 SV=1 |
| 6 | STK39 | STE20/SPS1-related proline-alanine-rich protein kinase OS=Homo sapiens GN=STK39 PE=1 SV=3 |
| 7 | MAP2K4 | Dual specificity mitogen-activated protein kinase kinase 4 OS=Homo sapiens GN=MAP2K4 PE=1 SV=1 |
| 8 | ZC3HAV1 | Zinc finger CCCH-type antiviral protein 1 OS=Homo sapiens GN=ZC3HAV1 PE=1 SV=3 |
| 9 | GTF2A1 | Transcription initiation factor IIA subunit 1 OS=Homo sapiens GN=GTF2A1 PE=1 SV=1 |
| 10 | EIF1AY | Eukaryotic translation initiation factor 1A, Y-chromosomal OS=Homo sapiens GN=EIF1AY PE=1 SV=4 |
| 11 | SMARCE1 | SWI/SNF-related matrix-associated actin-dependent regulator of chromatin subfamily E member 1 OS=Homo sapiens GN=SMARCE1 PE=1 SV=2 |
| 12 | LARP1 | La-related protein 1 OS=Homo sapiens GN=LARP1 PE=1 SV=2 |
| 13 | AHSA1 | Activator of 90 kDa heat shock protein ATPase homolog 1<br>OS=Homo sapiens GN=AHSA1 PE=1 SV=1 |
| 14 | RRP1 | Ribosomal RNA processing protein 1 homolog A OS=Homo sapiens GN=RRP1 PE=1 SV=1 |
| 15 | MRPS36 | 28S ribosomal protein S36, mitochondrial OS=Homo sapiens GN=MRPS36 PE=1 SV=2 |
| 16 | POM121C | Nuclear envelope pore membrane protein POM 121C OS=Homo sapiens GN=POM121C PE=1 SV=2 |
| 17 | POM121 | Nuclear envelope pore membrane protein POM 121 OS=Homo sapiens GN=POM121 PE=1 SV=2 |
| 18 | RAVER1 | Ribonucleoprotein PTB-binding 1 OS=Homo sapiens GN=RAVER1 PE=1 SV=1 |
| 19 | RAC3 | Ras-related C3 botulinum toxin substrate 3 OS=Homo sapiens GN=RAC3 PE=1 SV=1 |
| 20 | RAC2 | Ras-related C3 botulinum toxin substrate 2 OS=Homo sapiens GN=RAC2 PE=1 SV=1 |
| 21 | MT1M | Metallothionein-1M OS=Homo sapiens GN=MT1M PE=3 SV=2 |
| 22 | MT1G | Metallothionein-1G OS=Homo sapiens GN=MT1G PE=1 SV=2 |
| 23 | MT2A | Metallothionein-2 OS=Homo sapiens GN=MT2A PE=1 SV=1 |
| 24 | MT1X | Metallothionein-1X OS=Homo sapiens GN=MT1X PE=1 SV=1 |
| 25 | MT1E | Metallothionein-1E OS=Homo sapiens GN=MT1E PE=1 SV=1 |
| 26 | AIFM1 | Apoptosis-inducing factor 1, mitochondrial OS=Homo sapiens GN=AIFM1 PE=1 SV=1 |
| 27 | POLDIP3 | Polymerase delta-interacting protein 3 OS=Homo sapiens GN=POLDIP3 PE=1 SV=2 |
| 28 | SUPT4H1 | Transcription elongation factor SPT4 OS=Homo sapiens GN=SUPT4H1 PE=1 SV=1 |
| 29 | MRPS6 | 28S ribosomal protein S6, mitochondrial OS=Homo sapiens GN=MRPS6 PE=1 SV=3 |
| 30 | DDX18 | ATP-dependent RNA helicase DDX18 OS=Homo sapiens GN=DDX18 PE=1 SV=2 |
| 31 | DCAF13 | DDB1- and CUL4-associated factor 13 OS=Homo sapiens GN=DCAF13 PE=1 SV=2 |
| 32 | GTF2F2 | General transcription factor IIF subunit 2 OS=Homo sapiens GN=GTF2F2 PE=1 SV=2 |
| 33 | NOSIP | Nitric oxide synthase-interacting protein OS=Homo sapiens<br>GN=NOSIP PE=1 SV=1 |
| 34 | CTSB | Cathepsin B OS=Homo sapiens GN=CTSB PE=1 SV=3 |
| 35 | DDAH1 | N(G),N(G)-dimethylarginine dimethylaminohydrolase 1<br>OS=Homo sapiens GN=DDAH1 PE=1 SV=3 |
| 36 | GAA | Lysosomal alpha-glucosidase OS=Homo sapiens GN=GAA PE=1<br>SV=4 |
| 37 | MRPL16 | 39S ribosomal protein L16, mitochondrial OS=Homo sapiens<br>GN=MRPL16 PE=1 SV=1 |
| 38 | MCM2 | DNA replication licensing factor MCM2 OS=Homo sapiens<br>GN=MCM2 PE=1 SV=4 |
| 39 | ORC3 | Origin recognition complex subunit 3 OS=Homo sapiens<br>GN=ORC3 PE=1 SV=1 |
| 40 | DSC2 | Desmocollin-2 OS=Homo sapiens GN=DSC2 PE=1 SV=1 |
| 41 | IDH3A | Isocitrate dehydrogenase [NAD] subunit alpha, mitochondrial<br>OS=Homo sapiens GN=IDH3A PE=1 SV=1 |
| 42 | ITGA2 | Integrin alpha-2 OS=Homo sapiens GN=ITGA2 PE=1 SV=1 |
| 43 | GADD45GIP1 | Growth arrest and DNA damage-inducible proteins-interacting<br>protein 1 OS=Homo sapiens GN=GADD45GIP1 PE=1 SV=1 |
| 44 | SRSF8 | Serine/arginine-rich splicing factor 8 OS=Homo sapiens<br>GN=SRSF8 PE=1 SV=1 |
| 45 | CCAR2 | Cell cycle and apoptosis regulator protein 2 OS=Homo sapiens<br>GN=CCAR2 PE=1 SV=2 |
| 46 | OGFR | Opioid growth factor receptor OS=Homo sapiens GN=OGFR<br>PE=1 SV=3 |
| 47 | FAF2 | FAS-associated factor 2 OS=Homo sapiens GN=FAF2 PE=1<br>SV=2 |
| 48 | ACTL10 | Actin-like protein 10 OS=Homo sapiens GN=ACTL10 PE=2<br>SV=1 |

The proteins found upregulated in drug resistant cells are RPL27, RPS18, RPS3A, RPS4X, RPS13, H2BFS, RPS11, RPS24, RPL32, RPS15, RPS16, RPL30, RPL35A, RPL22L1, RPS8, RPL9, SBDS, ERO1A, P4HA1, UBQLN4, UBQLN1, CDA, H2AFY, H3F3A, PDHB, PSMC5, HEXOKINASE 2, SQSTM1, HISTH2BK, HISTH2BB HISTH2BC, HISTH2BD, HISTH2AB, HIST1H4A, HNRNPR, SNRPD1, NFKB2, COMT, RPS6, NDRG1, SORD, TMPO, PFKP, OSBP, DDX52, DDX21, EIF5, EWSR1, VDAC2, GPI, and GOT.

However, some of the important protein which are found downregulated are EPCAM, YBX1, PRSS1, LGALS1, STMN1, TARS, SAFB2, SERBP1, UQCRH, UBA52, ADH5, DPYSL2, TMSB4X, NUCKS1, HIF0, PPP1R14B, DBN1, PHGDH, CORO1B, ADRM1, PFDN6, Galectin-1 and GSTO1.

Further we also found a decrease in expression of TUBA1C; PARP1; TUBAL3; LMNA; SPTAN1; LMNB2; LMNB1 proteins. Interstingly, we also see increased expression of proteins related to chromosome condensation (Figure 3C) like H3C13, H3-4, H4C6, H2AC20, H4C6, H2AC18, H2BS1, H2AC8, H2BC12 etc. The decreased expression of Lamins and increased genes related to chromatin condensation further confirmed crippled apoptosis, a feature of drug-resistant cells.

**Figure 3:**
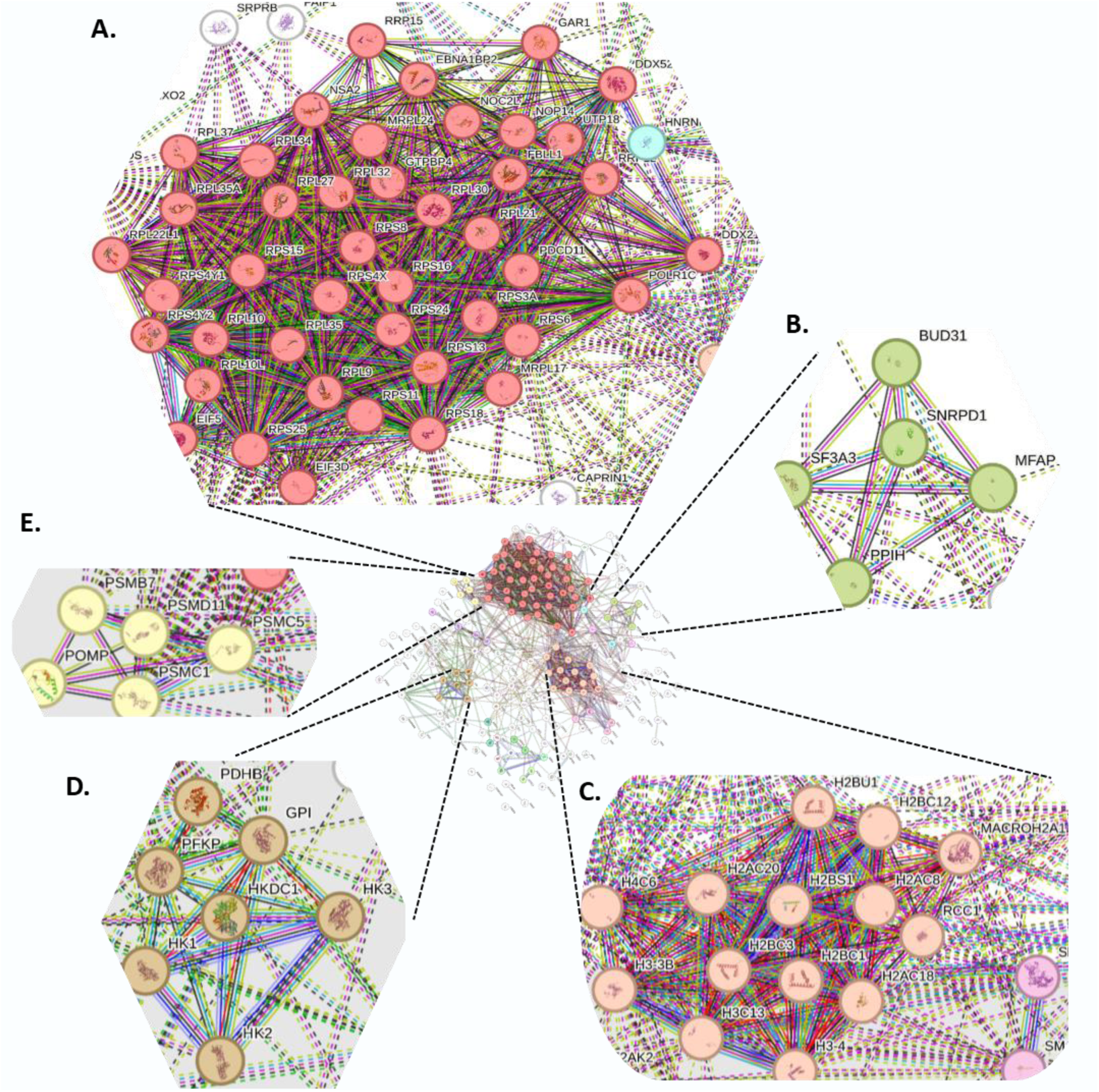
Proteome-wide protein-protein interactions were discovered in Reg-R-HCT116 cells. The STRING tool was used to analyse the protein-protein interactions discovered in the proteome data. Color classification: A) Red circles represent proteins involved in ribosome-associated pathways B) light green circles represent spliceosome mediated pathways C) Peach circles represent the genes involved in chromosome condensation during prophase. D) light brown circles represent protein processing in glycolysis/glucogenesis. E) light yellow circles represent proteins associated with proteasome assembly and degradation. The thick lines represent strong interactions, whereas the dotted lines represent expected interactions.

**Figure 4.**
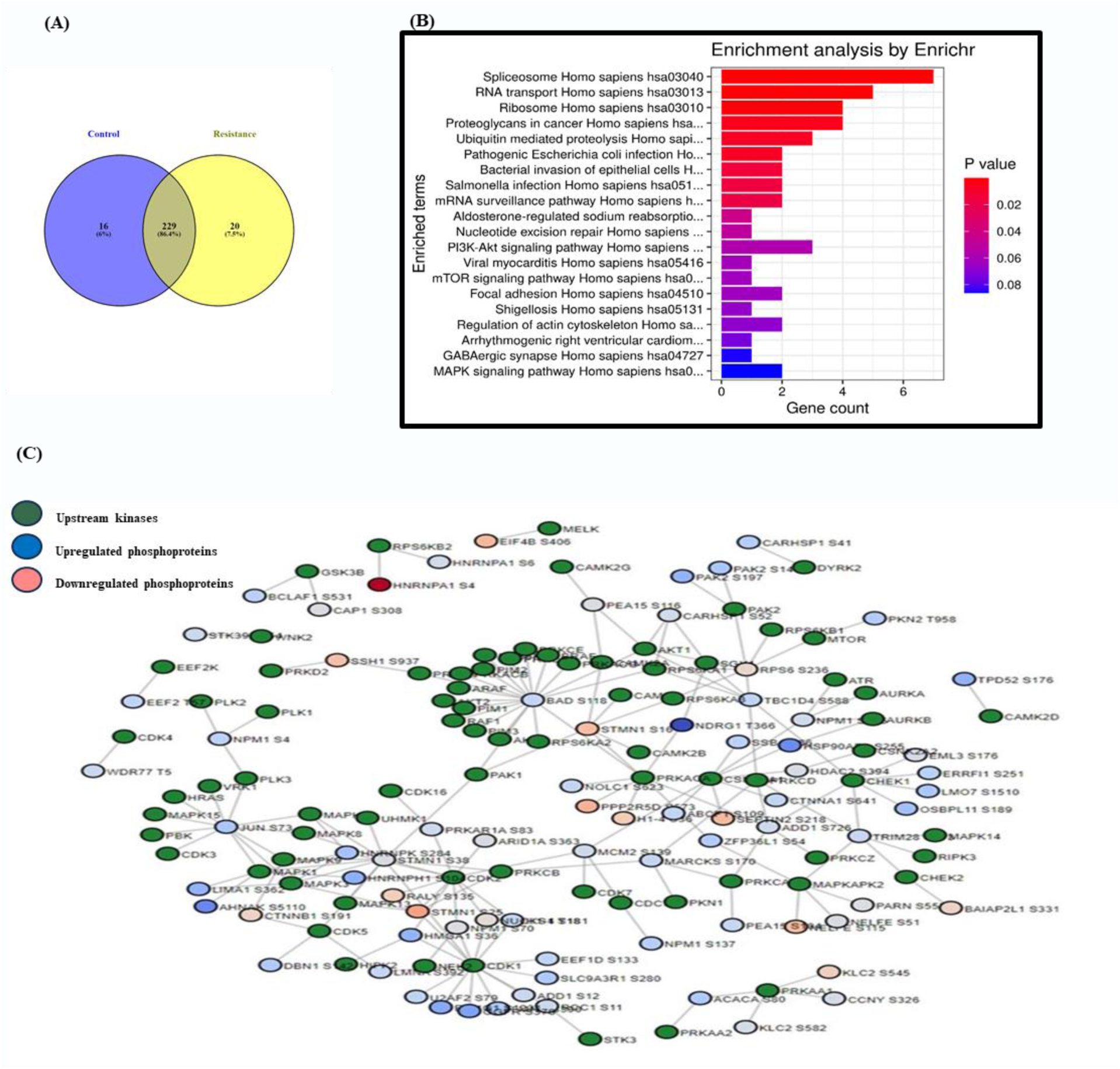
Phospho-proteomics data analysis. (A) Venn diagram shows total number of phosphorylated proteins found after analysis of HCT116 and Reg-R-HCT116 samples. (B) Pathways enrichment analysis. This graph shows the gene ontology pathways of the significant Proteins we found in Phospho-proteomics data. (C) Network analysis of phosphorylated proteins and their upstream kinase in Reg-R-HCT116 using Phosphomatics tool. The blue circles indicate upregulated phosphorylation, red indicates downregulated phosphorylation and green circles indicate their upstream kinases.

### 3.3 Pathway analysis

Further, to check the functional relevance of the differentially expressed proteins, pathway analysis was performed by EnrichR (Figure 2B), we found signaling pathways such as Ribosome, metabolic pathways, drug metabolism, HIF-1 signaling, insulin signaling, pentose phosphate pathway to be enriched in our resistant cell line. As most of the proteins upregulated are ribosomal proteins which are associated to mTOR and eIF2 signaling, we look for the upstream regulator of eIF2 signaling through Ingenuity pathway analysis (IPA). We see in Figure 2C Sirolimus (Rapamycin), Torin1 and LARP1 highly inhibits eIF2 signaling with high negative z-score value.

### 3.4 Protein-Protein Interaction Network

The differentially expressed proteins and their interacting partner’s network were generated using String tool. Figure 3 represents a network of differentially expressed proteins with their interactome. Color classification: A) Red circles represent proteins involved in ribosome-associated pathways B) light green circles represent spliceosome mediated pathways C) peach circles represent the genes involved in chromosome condensation during prophase. D) light brown circles represent protein processing in glycolysis/glucogenesis. E) light yellow circles represent proteins associated with proteasome assembly and degradation. The thick lines represent strong interactions, whereas the dotted lines represent expected interactions.

In the protein-protein network analysis through string and EnrichR we see that Ribosome associated pathways which comes under eIF2 signaling is highly upregulated. Some of the genes related to eIF2 signaling which are highly expressed with log2FC > 1.5 are listed in Table 5. Apart from proteins related to eIF2 signaling other proteins associated to PI3K/Akt/mTOR pathway are also highly upregulated like NDRG1, Rictor, NFKB2, AKT1S1, LAMB3, GNG12, GNG5, CDC37, PPP2R1A, ITGB1, ITGB3, ITGB4, RICTOR, RPS6 and MDK. (Table 1).

### 3.5 Differentially expressed phosphoproteins in Reg-R-HCT116

The phosphomatics V2B provides a suite of tools for the downstream analysis of global phospho-proteomics data acquired by LC-MS/MS. Regorafenib being a multi kinase inhibitor phospho-proteomics analysis was performed to check the exact mechanistic pathways or changes in phosphorylation status of different proteins involved in developing resistance. The obtained raw data was normalized followed by PCA, Correlation and distribution patterns analysis as shown in (Supplementary Data). This data was then analysed for differential phosphorylation of proteins. As represented in Figure 3 a total of 265 differentially phosphorylated proteins are identified. 229 differentially phosphorylated proteins are found common in both HCT116 and Reg-R-HCT116 cells. 20 proteins with differential phosphorylation are found specifically only in Reg-R-HCT116 cells whereas 16 are found specifically only in HCT116 cells. Further to identify the number of specific phosphorylation sites the phosphopeptide-spectral matches with a false discovery rate (FDR) of 0.05. were used. This analysis identified 348 unique phosphor-peptides corresponding to the 265 differentially phosphorylated proteins. The list of differentially phosphorylated proteins and their fold changes are represented in Table 6 and Table 7.

**Table 6.** The list of significantly hypo-phosphorylated proteins and their fold change in Reg-R-HCT116 are represented in this table.

| S.No. | Gene Name | Abundance Ratio (log2): (T, Sample) / (C, Sample) | Positions in Master Proteins |
| --- | --- | --- | --- |
| 1 | BAZ1A | -5.70919505 | Q9NRL2 [1410-1423] |
| 2 | PLEC | -5.493908098 | Q15149 [2789-2808] |
| 3 | RSBN1 | -4.645359292 | Q5VWQ0 [61-83] |
| 4 | ESYT2 | -4.623447544 | A0FGR8 [661-682] |
| 5 | SAFB | -4.409141379 | Q15424 [571-585] |
| 6 | SRRM2 | -3.806552757 | Q9UQ35 [846-870] |
| 7 | TNIK | -3.767903119 | Q9UKE5 [762-785] |
| 8 | STMN1 | -3.710283707 | P16949 [15-29] |
| 9 | KLC2 | -3.644068599 | Q9H0B6 [555-562] |
| 10 | HNRNPM | -3.533955969 | P52272 [525-531] |
| 11 | PLEC | -3.510776357 | Q15149 [4384-4401] |
| 12 | PDAP1 | -3.481862331 | Q13442 [173-181] |
| 13 | SLK | -3.294265072 | Q9H2G2 [1083-1091] |
| 14 | SNRNP70 | -3.273080801 | P08621 [210-231] |
| 15 | PHLDB2 | -3.227475698 | Q86SQ0 [925-954] |
| 16 | TCOF1 | -3.190762826 | Q13428 [1348-1360] |
| 17 | PPP1R12A | -3.184146759 | O14974 [694-709] |
| 18 | RBM17 | -3.171937774 | Q96I25 [150-166] |
| 19 | CAP1 | -3.170949636 | Q01518 [288-313] |
| 20 | TBC1D10B | -3.157142059 | Q4KMP7 [14-29] |
| 21 | SRRM2 | -3.108499292 | Q9UQ35 [2578-2591] |
| 22 | TRA2B | -3.056427892 | P62995 [189-217] |
| 23 | MTA1 | -3.045438934 | Q13330 [634-645] |
| 24 | SRRM2 | -3.016422226 | Q9UQ35 [2393-2406] |
| 25 | PHLDB1 | -3.005742099 | Q86UU1 [561-573] |
| 26 | TNIK | -2.971638214 | Q9UKE5 [762-785] |
| 27 | HNRNPM | -2.89754623 | P52272 [571-578];[584-591] |
| 28 | PRRC2A | -2.855125361 | P48634 [453-466] |
| 29 | TNIK | -2.83244977 | Q9UKE5 [699-712] |
| 30 | ABLIM1 | -2.790943695 | O14639 [585-598] |
| 31 | IRF2BP1 | -2.772192413 | Q8IU81 [382-403] |
| 32 | C9orf142 | -2.695904563 | Q9BUH6 [138-150] |
| 33 | ENSA | -2.678119275 | O43768 [107-121] |
| 34 | ACIN1 | -2.656965431 | Q9UKV3 [823-847] |
| 35 | ARHGAP29 | -2.641941608 | Q52LW3 [1142-1156] |
| 36 | HNRNPA2B1 | -2.641513959 | P22626 [326-350] |
| 37 | CAP1 | -2.476551722 | Q01518 [288-312] |
| 38 | SRRM2 | -2.437560214 | Q9UQ35 [2405-2418] |
| 39 | CUL4A | -2.360368856 | Q13619 [2-17] |
| 40 | BCLAF1 | -2.343058865 | Q9NYF8 [638-653] |
| 41 | KRT19 | -2.291935745 | P08727 [25-51] |
| 42 | ZC3H13 | -2.28222931 | Q5T200 [875-888] |
| 43 | DDA1 | -2.133372303 | Q9BW61 [89-102] |
| 44 | TRIM28 | -2.07272547 | Q13263 [33-71] |
| 45 | NAB2 | -2.064146603 | Q15742 [1-20] |
| 46 | ARID1A | -1.976825117 | O14497 [1880-1906] |
| 47 | AJUBA | -1.910962337 | Q96IF1 [256-275] |
| 48 | FNBP4 | -1.88334821 | Q8N3X1 [7-23] |
| 49 | EIF3G | -1.856988437 | O75821 [33-57] |
| 50 | RPS6 | -1.74892771 | P62753 [234-249] |
| 51 | SEPT9 | -1.728249211 | Q9UHD8 [77-91] |
| 52 | SRRM2 | -1.688727413 | Q9UQ35 [919-926] |
| 53 | LMNB1 | -1.678459663 | P20700 [15-29] |
| 54 | BAG3 | -1.674532646 | O95817 [122-139] |
| 55 | CD97 | -1.640693363 | P48960 [827-835] |
| 56 | RPL34 | -1.605139309 | P49207 [10-21] |
| 57 | IRF2BP1 | -1.571495804 | Q8IU81 [382-403] |
| 58 | NAB2 | -1.333638137 | Q15742 [1-21] |
| 59 | CCDC86 | -1.110285753 | Q9H6F5 [10-27] |
| 60 | RBMX | -1.091575761 | P38159 [204-223] |

**Table 7.** The list of significantly hyper-phosphorylated proteins and their fold change in Reg-R-HCT116 are represented in this table.

| S.No. | Gene Name | Abundance Ratio (log2): (T, Sample) / (C, Sample) | Positions in Master Proteins |
| --- | --- | --- | --- |
| 1 | NDRG1 | 1.104480729 | Q92597 [364-388] |
| 2 | DOCK5 | 1.159387299 | Q9H7D0 [1742-1759] |
| 3 | SLTM | 1.160100493 | Q9NWH9 [549-564] |
| 4 | MISP | 1.160586237 | Q8IVT2 [469-496] |
| 5 | NEDD4L | 1.360447145 | Q96PU5 [446-467] |
| 6 | UBAP2L | 1.466460449 | Q14157 [439-459] |
| 7 | SP100 | 1.573028523 | P23497 [350-386] |
| 8 | PKN2 | 1.574592639 | Q16513 [581-594] |
| 9 | SUPT5H | 1.593351813 | O00267 [653-668] |
| 10 | LAD1 | 1.640971738 | O00515 [266-282] |
| 11 | RPS8 | 1.651295969 | P62241 [125-139] |
| 12 | ACIN1 | 1.803670732 | Q9UKV3 [873-904] |
| 13 | EPPK1 | 1.857109174 | P58107 [2713-2732] |
| 14 | PKP2 | 1.861917315 | Q99959 [147-158] |
| 15 | AHNAK | 1.883628215 | Q09666 [5820-5831] |
| 16 | AHNAK | 1.913701635 | Q09666 [2386-2399] |
| 17 | SNTB2 | 1.914538917 | Q13425 [90-115] |
| 18 | CAV1 | 1.932364135 | Q03135 [31-47] |
| 19 | PROSER2 | 2.015806732 | Q86WR7 [41-52] |
| 20 | PALLD | 2.183322634 | Q8WX93 [632-654] |
| 21 | SRRM2 | 2.36046604 | Q9UQ35 [1690-1699] |
| 22 | RANBP2 | 2.557192546 | P49792 [1140-1185] |
| 23 | NDRG1 | 2.57670983 | Q92597 [362-388] |
| 24 | CTTN | 2.747879413 | Q14247 [397-428] |
| 25 | EIF4ENIF1 | 2.777705424 | Q9NRA8 [562-574] |
| 26 | AHNAK | 2.790344874 | Q09666 [488-499] |
| 27 | SRRM1 | 2.872127073 | Q8IYB3 [614-623] |
| 28 | MISP | 2.882596858 | Q8IVT2 [469-481] |
| 29 | UBE2O | 2.92568529 | Q9C0C9 [826-849] |
| 30 | MVP | 3.009837665 | Q14764 [861-893] |
| 31 | RPS3 | 3.314812633 | P23396 [202-243] |
| 32 | AHNAK | 3.37944935 | Q09666 [2386-2399] |
| 33 | THOC2 | 5.024831246 | Q8NI27 [1283-1297] |

The phosphor-proteomic analysis identified a panel of hypo-phosphorylated proteins with an abundance ratio (log₂) less than –1. These include chromatin regulators such as BAZ1A, ARID1A, TRIM28, MTA1, and ZC3H13; cytoskeletal and structural proteins including PLEC, KLC2, STMN1, CAP1, ABLIM1, KRT19, SEPT9, LMNB1, and BAG3; RNA-binding and splicing factors such as RSBN1, SAFB, SRRM2, HNRNPM, HNRNPA2B1, RBM17, RBMX, TRA2B, PRRC2A, ACIN1, and SNRNP70; signaling and regulatory proteins including TNIK, SLK, PPP1R12A, TBC1D10B, ENSA, NAB2, IRF2BP1, AJUBA, and ARHGAP29; and translation-related factors including EIF3G, RPS6, and RPL34. Additional hypo-phosphorylated proteins detected were ESYT2, PHLDB1, PHLDB2, TCOF1, PDAP1, FNBP4, DDA1, CUL4A, BCLAF1, CD97, CCDC86, and C9orf142. Collectively, the downregulation of phosphorylation across these diverse functional groups underscores potential alterations in chromatin remodelling, RNA processing, cytoskeletal organization, and signaling pathways that may contribute to the observed phenotype.

In contrast to the hypo-phosphorylated proteins identified, a set of hyperphosphorylated proteins with an abundance ratio (log₂) greater than 1, representing diverse functional categories were identified. Several of these proteins were associated with cytoskeletal organization and adhesion, including MISP, LAD1, EPPK1, PKP2, AHNAK, CTTN, CAV1, PALLD, and SNTB2, suggesting enhanced regulation of actin Remodelling and cell junction dynamics. Proteins involved in RNA binding, transcription, and splicing regulation included SLTM, SUPT5H, SRRM2, SRRM1, and THOC2, pointing to possible alterations in mRNA processing and transcriptional control. Members linked to signaling and ubiquitin regulation were NDRG1, NEDD4L, PKN2, UBE2O, PROSER2, and DOCK5, implicating pathways related to stress signaling, protein degradation, and cell signaling networks. Proteins associated with nuclear pore complex and transport included RANBP2 and EIF4ENIF1, indicating potential changes in nucleocytoplasmic trafficking. Finally, ribosomal and translational regulators such as RPS8 and RPS3, along with structural/scaffold proteins including UBAP2L and MVP, were also hyperphosphorylated, suggesting modulation of translation and stress granule dynamics

### 3.6 KEGG pathways analysis of differentially expressed Phosphoproteins in Reg-R-HCT116

Based on network analysis by KEGG and pathway analysis using Enrichr, we found signaling pathways such as Spliceosome, Ribosome, proteoglycans in cancer, mTOR signaling, PI3K/AKT signaling, regulation of actin cytoskeleton in our KEGG pathway analysis we found this signaling enriched and upregulated in our phospho-proteome analysis.

### 3.7 Validation of proteins expressed specifically in Reg-R-HCT116 cells by RT-PCR

The differentially expressed proteins like DKK1, CRTC, VDAC3, TJP, RAP1, FMR1, NUP and HECTD1 recorded in the proteome analysis of Reg-R-HCT116 cells are further validated to confirm the observed expression pattern. These are validated at the transcriptional level by real-time-qPCR. The qPCR data, as shown in Figure 5, further confirmed the increased expression of these genes at the transcriptional level as well. The VDAC3 gene showed the highest expression change of a 5-fold increase, whereas the CRTC gene exhibited the lowest expression change of around 1.5-fold increase. All other genes exhibited a change ranging from a 2-2.5-fold increase and have also shown the same pattern of gene expression at the transcriptional level.

**Figure 5.**
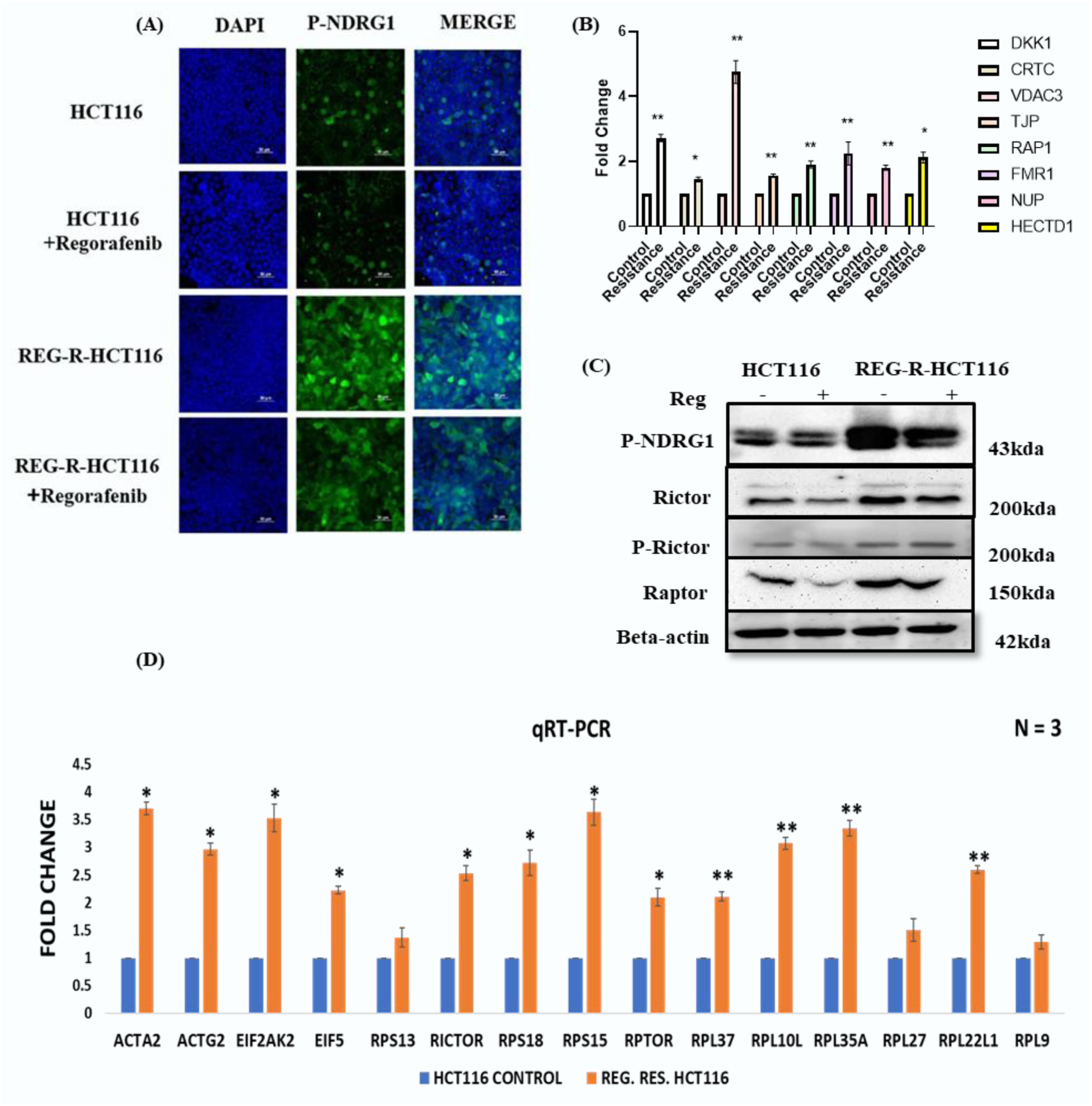
(A)- HCT116 Control and HCT116 Regorafenib Resistance cells were cultured and seeded for Immunocytochemistry to check the expression of p-NDRG1. (B) Graphical representation of Validation of gene expression of cells DKK1, CRTC, VDAC3, TJP, RAP1, FMR1, NUP, HECTD1 upregulated proteins we found in our proteome data of Reg-R-HCT116 cell line in comparison to drug sensitive HCT116 cells. (C) Western Blot analysis showing increased expression of p-NDRG1, Rictor, p-Rictor and Raptor in resistant cells as compared to control HCT116 cells (D) Graphical representation of Validation of gene expression of cells ACTA1, ACTG2, EIF2AK2, EIF5, RPA13, RICTOR, RPS18, RPS15, RPTOR, RPL37, RPL10L, RPL35A, RPL27, RPL22L1, RPL9. *p < 0.05, **p < 0.01, ***p < 0.001.

Further, the Phospho-proteome data was validated to rule out false positives. The validation of hyperphosphorylated proteins is done by immunocytochemistry and western blot analysis. According to the results observed in Phospho-proteome data of Reg-R-HCT116 cells exhibited hyperphosphorylation of NDRG1 and ACINUS in comparison to HCT116 cells. This is first evaluated by immunocytochemistry. As shown in Figure 5(A), HCT116 cells showed the basal level intensity of green fluorescence showing phosphorylation of NDRG1 protein and a decrease in intensity of green fluorescence was noticed in regorafenib-treated HCT116 cells. In contrast, Reg-R-HCT116 cells exhibited a higher green fluorescence intensity confirming hyperphosphorylation of NDRG1. A slight decrease in fluorescence intensity was observed in regorafenib-treated Reg-R-HCT116 cells, but this intensity was higher than the fluorescence intensity observed in control HCT116 cells. Thus, showing hyperphosphorylation of NDRG1. Similar phosphorylation status is reported even in the case of ACINUS protein **Supplementary 2.** The Reg-R-HCT 116 exhibited higher phosphorylation even in the presence of the drug.

This was further validated by western blots analysis, and the fold change was quantitated by densitometric analysis. The HCT116 cells showed, a basal level expression of phosphorylated NDRG1 and ACINUS, but upon exposure to regorafenib, a decrease in expression was observed **Supplementary 3 (A)**. In contrast, the Reg-R-HCT116 cells showed a significant increase in phosphorylated NDRG1 and ACINUS expression. Regorafenib-treated Reg-R-HCT116 cells showed a slight decrease in these, but the expression levels are higher than in HCT116 cells. This result is supported by the densitometric analysis of the western blots, as shown in **Supplementary 3 (B)**.

The cleavage of ACINUS during apoptosis induces chromatin condensation. The phosphorylation of this protein reduces the chromatin condensation during apoptosis. The upstream AKT regulates the phosphorylation of ACINUS (Hu et al., 2005). This observation further points to the activation of AKT signaling in Reg-R-HCT116 cells.

Additionally, as reported in the literature, the phosphorylation of NDRG1 is regulated by both AKT and the mTORC2 complex. Moreover, NDRG1 serves as a substrate for AKT and is involved in vesicle trafficking, and role in drug resistance as well (zhang et al., 2015) (Valluri et al., 2023) (Villodre et al., 2022) (Weiler et al., 2014) indicating that the increased expression of phosphorylated NDRG1 and ACINUS, along with the activation of upstream molecules, is linked to the PI3K/AKT/mTOR pathway. Notably, AKT is a key downstream component of the PI3K pathway, further establishing its involvement in these cellular processes.

Elevated SGK1 predicts resistance of breast cancer cells to AKT inhibitors (Sommer et al., 2013), mTOR Regulation of N-Myc Downstream Regulated 1 (NDRG1) Phosphorylation in Clear Cell Renal Cell Carcinoma (Valluri et al., 2023).

In accordance with the finding’s upregulation and phosphorylation of several proteins suggested the involvement and activation of upstream molecules associated with the PI3K/AKT/MTOR pathways. For instance, we observed an upregulation of NFKB in our proteome data, known to be regulated by AKT. This indicates the involvement of AKT in the activation of NFKB. Furthermore, we identified p-NDRG1, a substrate for AKT and regulated by the mTORC2 complex, which is known to participate in vesicular trafficking.

Moreover, our Phosphoproteome data highlighted the phosphorylation of Met and Rictor. Met has been associated with the activation of VEGFR-3 signaling. While Rictor is known to enhance this activation, further contributing to the activation of downstream pathways such as mTOR signaling. Being a part of mTORC2 it also regulates the phosphorylation of AKT at ser473 (Sarbassov et al., 2005).

We conducted western blotting experiments to verify these findings, as depicted in Supplementary Figure. We compared the expression levels of key upstream molecules in both the cell lines with or without regorafenib treatment. In Reg-R-HCT116 cells, we examined the expression levels of phosphorylated AKT for its activation. Additionally, we checked the expression levels of Rictor, p-Rictor, and Raptor to evaluate the activation of mTOR signaling Supplementary Figure. shows a significant increase in the expression of Rictor protein in Reg-R-HCT116 cells even in the presence of regorafenib. Whereas in the control cells, we could see a decrease in expression in drug-treated cells. A similar expression pattern was noticed in phosphorylated Rictor and Raptor proteins also. The densitometric analysis plotted in 3.5 B also supports the same.

Other AKT-associated proteins recorded in our data, such as VEGF3, NFKB and phosphorylation of AKT, were also evaluated. In Figure 3.6A, we could see an increase in VEGF3 expression in drug-resistant cells even in the presence of regorafenib. A similar pattern was noticed in the expression of NFKB. A significant increase in phosphorylation of AKT at ser473 was noticed in Reg-R-HCT116 cells in comparison to HCT116 cells, whereas a slight increase in phosphorylation of thr 308 was noitced. The same was confirmed by densitometric analysis (Data not shown). Which further validated our proteome and phosphor-proteome and confirmed the activation of PI3K/AKT/MTOR signaling in Reg-R-HCT116 cells. These results pointed out at the involvement of mTOR and eIF2 signaling pathways. This was first confirmed through q-RT-PCR analysis of the genes associated with mTOR and eIF2 signaling pathways in the HCT116 control and Reg-R-HCT116 cell lines. The gene expression levels observed to be elevated in proteome profiling within the mTOR/eIF2 pathway were also supported by the qRT-PCR results, thereby validating our proteome profiling findings Figure 5D. The Fold increase of all the eIF2 and mTOR signaling related genes through q-RT-PCR analysis is shown in **S4.** ACTA2, EIF2AK2, RPS15, RPL10L, RPL35A has fold change above three. However, ACTG2, EIF5, RICTOR, RPS18, RPTOR, RPL37, and RPL22L1 shows fold change value above 2. So, the increase in eIF2 and mTOR signaling were validated through q-RT-PCR analysis. Once the hyperactivation of mTOR and eIF2 signaling was confirmed, the impact of inhibition of this pathway on resistance was further evaluated. Both rapamycin and Torin 1 are potent inhibitors of mTOR as well as eIF2 signaling pathway whereas LARP1 only inhibits eIF2 signaling by inhibiting the translation. Rapamycin specifically targets mTORC1 whereas Torin1 targets the kinase activity of both mTORC1 as well as mTORC2.

### 3.8 Cytotoxicity Assay:-

After validation of upregulated eIF2 and mTOR signaling, the impact of inhibition of this signaling was assessed on both HCT116 and Reg-R-HCT116 cells. After 48 hours of incubation, the cytotoxic effects of Torin1 and its combinations with regorafenib were investigated in HCT116 and Reg-R-HCT116 cells Figure 6 (A&B). In the HCT116 cell line as well as Reg-R-HCT116 cells, the IC50 values for Torin1 were found to be around 5-6 µM. Torin1 exhibits a nearly equivalent cytotoxic impact on both control and resistant cell lines. The combination study of Regorafenib was also assessed using Torin1 on both control and resistant cell lines showing synergistic effects on both the cell line, specifically on the resistant cell line, as indicated in Figure 6 (C&D).

**Figure 6:**
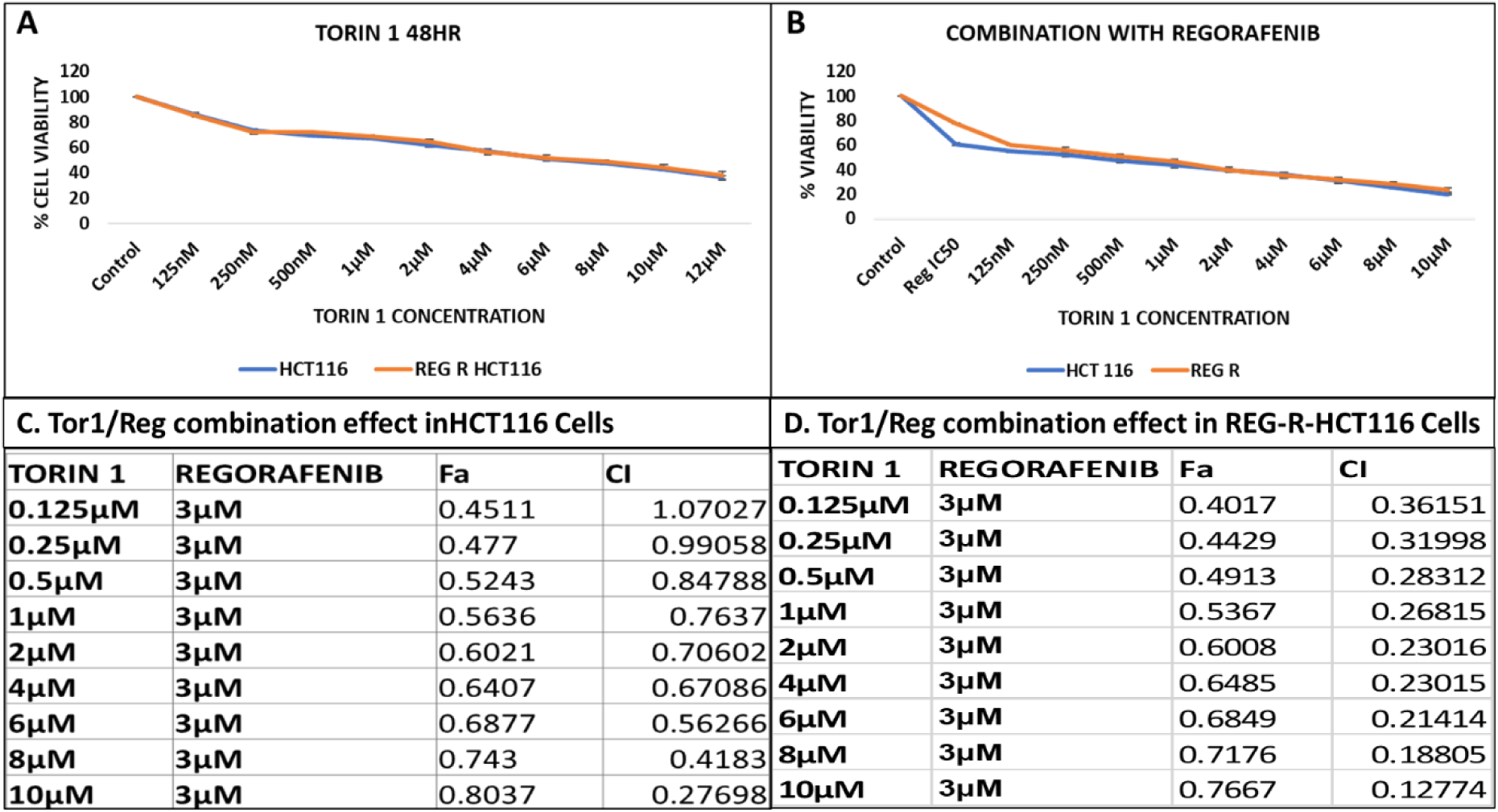
The cytotoxic effects of Torin 1 and its combination with Regorafenib was assessed against Control HCT116 and Reg-R-HCT116 cells. The X-axis represents the drug concentration, while the Y-axis indicates the percentage of cell viability. Figures A and B depict the Cell Viability Graph of HCT116 cells and Reg-R-HCT116 cells following treatment with (A) Torin 1, and (B) escalating doses of Torin 1 with an IC50 of Regorafenib (3uM) (Δ Tor/Reg), as determined by the MTT assay. The presented values represent means ± standard deviation of experiments conducted in triplicates, and statistical significance was determined. All data indicate significant p-values. Figure (C and D) Shows the synergistic impact of the Δ Tor/Reg pairing in Control HCT116 and Reg-R-HCT116 cell lines. Various drug treatments are presented with the Combination Index (CI) and Fraction affected of viable cells (Fa).

Combination index (CI) expressing Synergism was evaluated according to Chou and Talalay using Compusyn Software ver.1.0 (Combosyn, lnc., Paramus, NJ, 07652, USA). Fa indicates fraction affected, corresponding to the fraction of cell viability (%); CI means combination index, where CI < 1 indicates synergism, CI > 1 indicates antagonism, and CI = 1 indicates additive effect.

The reported CI index values in the figure suggest that the observed combination yielded values are less than 1, signifying a synergistic effect of the combination.

### 3.9 qRT-PCR analysis:-

Further, Real-time quantitative polymerase chain reaction (qPCR) was used to evaluate the gene expression profiles of PI3K/Akt/mTOR and eIF2 signaling-related genes in cells subjected to treatment with Torin1, and its combinations with regorafenib was evaluated at transcriptional level by qRT-PCR. Torin1 stands out as a potent allosteric ATP-competitive inhibitor of mTOR kinase, effectively targeting both complexes. Importantly, Torin 1 exerts no influence on the stability of both mTORC1 and mTORC2. It exhibits selectivity for mTOR over closely related kinases (Sun, 2013)(Thoreen et al., 2009). In Real-time qPCR analysis, both HCT116 and Reg-R-HCT116 cells were subjected to treatment with the IC50 concentration of Torin1 (6µM). Notably, in both HCT116 cell line and Reg-R-HCT116 cell line, Torin1 downregulates the expression of genes related to eIF2 signaling, such as EIF5, RPS15, and RPL27, while maintaining the expression levels of mTOR, Raptor, and Rictor, indicating effective inhibition of mTOR-related kinase activity Figure 7 (A&B).

**Figure 7:**
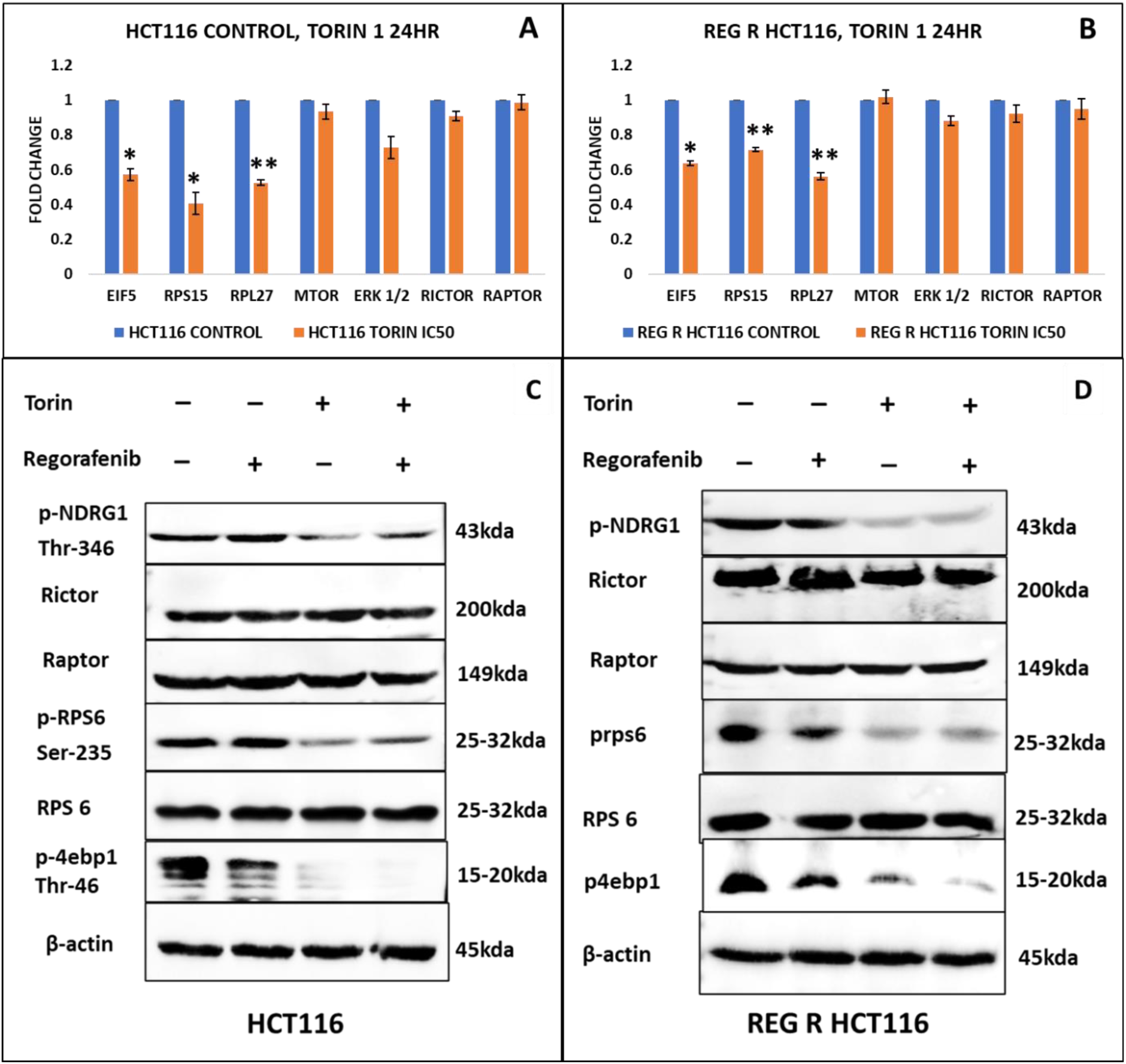
The graph shows gene expression profiles of EIF5, RPS15, RPL27, mTOR, ERK1/2, RICTOR, and RAPTOR, with gene names on the X-axis and fold change on the Y-axis. (A, B) Torin 1 treatment (6 µM) significantly reduces EIF5, RPS15, and RPL27 expression in both HCT116 and Reg-R-HCT116 cells, while mTOR, ERK1/2, RICTOR, and RAPTOR remain unchanged. (C, D) Western blot analysis at IC50 (6 µM) shows reduced levels of phospho-NDRG1 (Thr346), phospho-RPS6 (Ser235), and phospho-4E-BP1 (Thr46) in both cell lines, with no significant change in RICTOR, RAPTOR, RPS6, and eIF4E expression. Statistical significance is indicated as p < 0.05, ** p < 0.01, *** p < 0.001.

In conclusion, the same IC50 value in torin1 (6 μM) in both control and resistant cells, emphasizes its broader inhibition capabilities, effectively targeting cellular processes in both cell lines. Further, to evaluate the inhibition of kinase activity of both mTOR1 and mTOR2 by torin1 western blot analysis was performed.

### 3.10 Western Blot Analysis

Western Blot analysis was conducted to examine the mTOR-related kinase activity and proteins involved in PI3K/Akt/mTOR signaling, by evaluating the impact of Torin 1 on both HCT116 and Reg-R-HCT116 cells. mTORC1, known for its role in regulating protein translation by phosphorylating S6 Kinase (S6K) and eIF4E-binding protein 1 (4E-BP1), however mTORC2 regulates SGK1 which further phosphorylates NDRG1.

The phosphorylation status of key proteins, including 4E-BP1, phospho-4E-BP1, RPS6, phospho-RPS6, Rictor, Raptor, and phospho-NDRG1 was assessed upon treatment with Torin 1 (6 μM) on both control and resistant cell lines. In the HCT116 cell line, while the expression levels of Rictor and Raptor remained stable, there was a notable reduction in the phosphorylation levels of 4E-BP1, RPS6, and NDRG1. This suggests that Torin 1 inhibits cell growth and protein translation by suppressing both mTOR1 and mTOR2 kinase activity. Similar results were observed in Reg-R-HCT116 cells, indicating that blocking the ATP-dependent kinase activity of mTOR has a significant impact on both mTORC1 and mTORC2 signaling Figure 8 (C&D).

**Figure 8:**
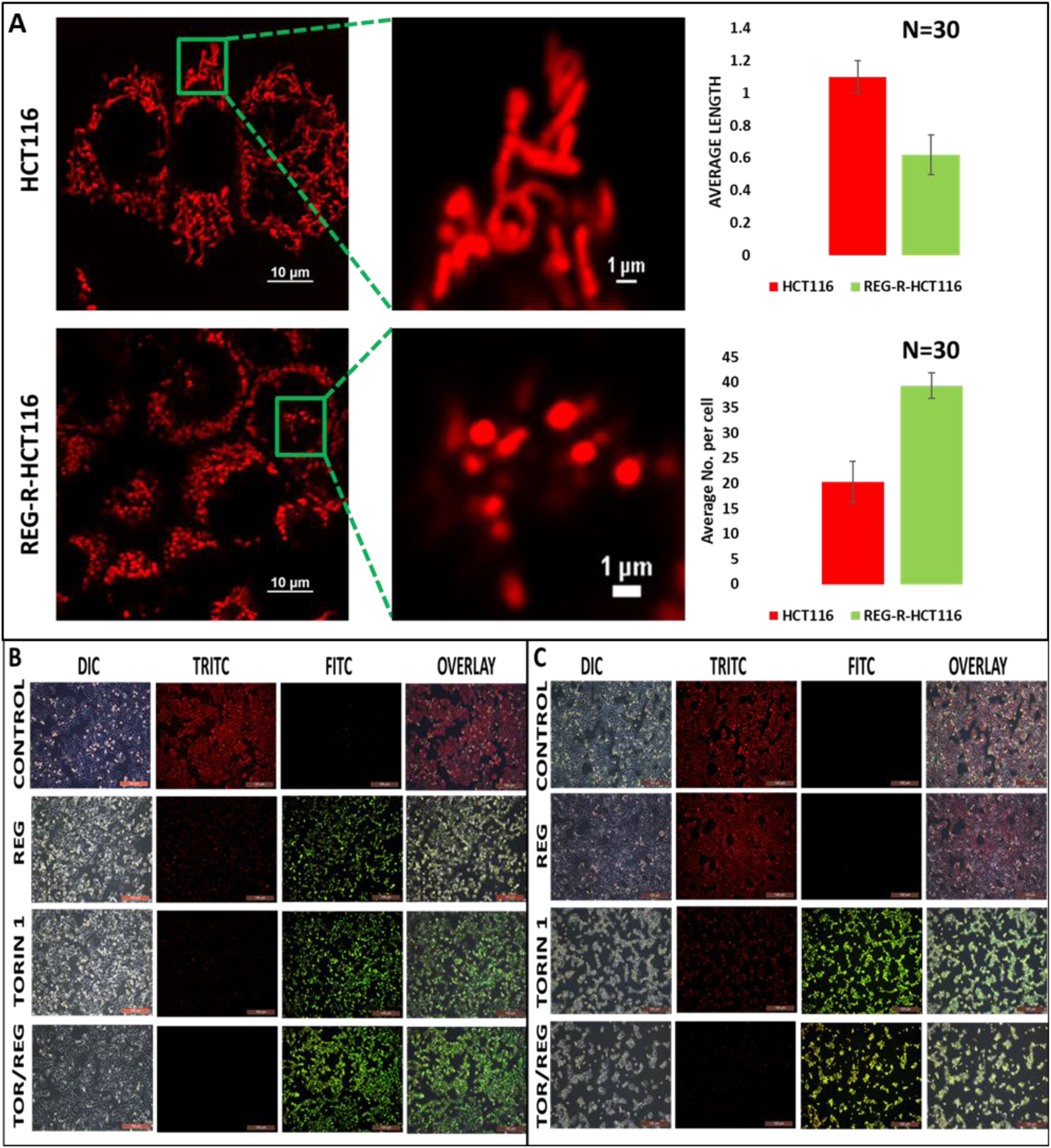
Shows mitochondrial membrane potential (MMP) loss using JC-1 staining on (A) HCT116 cells, (B) REG-R-HCT116 cells. Both treated with IC50 concentration of regorafenib, Torin 1, and Tor1/Reg combination imaged under a fluorescence microscope with 20× magnification. Scale bar 100 µm. Red—Intact mitochondria and Green— MMP loss.

### 3.11 Visualization of Mitochondria by Confocal Analysis:-

Mito Tracker Deep Red was employed to observe the morphology and integrity of mitochondria. The continuous proliferation of cancer cells requires ongoing alterations in energy metabolism to fuel cellular growth and division efficiently. It is established that mTORC1 facilitates mitochondrial fission by regulating the translation of the mitochondrial fission factor MTFP1, via 4E-BP mediation. Suppression of mTORC1 function, whether through pharmaceutical means or genetic techniques, results in mitochondrial hyper fusion, branching, and circularization (de la Cruz López et al., 2019). Because of the well-documented involvement of mTORC1 in regulating mitochondrial dynamics and cell viability, we aimed to investigate changes in mitochondrial structure in both our Control and Resistant cell lines. Our study using confocal imaging revealed a notable mitochondrial fission pattern in the Reg-R-HCT116 cell line, marked by decreased mitochondrial size and increased number as compared to the control HCT116 cell line Figure 8 (A). The observed results indicate an increased energy requirement in the resistant cell line, leading to mitochondrial fission. Additionally, mitochondrial fission serves as a mechanism utilized during hypoxic conditions to regulate reactive oxygen species (ROS) production at a physiologically low level and maintain cellular integrity by diminishing respiratory activity (Fuhrmann & Brüne, 2017).

### 3.12 Utilizing JC-1 Staining for Assessment of MMP Depletion:-

To further confirm the occurrence of cell death and the role of mitochondria in this process, we conducted JC-1 staining. Both HCT116 control and resistant cells were subjected to IC50 concentrations of Regorafenib, and Torin1. The disruption of mitochondrial membrane potential (MMP) is a significant feature of apoptosis. JC-1, a lipophilic cationic dye, was used to evaluate MMP loss. In healthy cells, this dye accumulates in the mitochondria, which are negatively charged, leading to the formation of J-aggregates emitting red fluorescence. Conversely, in apoptotic cells undergoing a decline in mitochondrial membrane potential, JC-1 penetrates the mitochondria but fails to form stable J-aggregates due to decreased negative charge. Consequently, the dye remains in a monomeric state, emitting green fluorescence (Sivandzade et al., 2019). Fluorescence microscopy was utilized to visualize JC-1 staining, using both TRITC and FITC channels. The resultant images were merged to analyse the spatial distribution of J-aggregates in comparison to J-monomers.

As depicted in the Figure 8 (B&C), both the HCT116 control and Reg-R-HCT116 cells were treated with the IC50 concentration of Regorafenib, Torin 1 and IC25 of both Regorafenib and Torin1 (1.5 µM and 3 µM respectively). HCT116 cells induce MMP loss against all the drug treatment, whereas Reg-R-HCT116 cells exhibited elevated MMP loss, particularly with Torin 1 and the combination of Regorafenib and Torin 1. This observation suggests that Torin 1 prominently triggers MMP loss in resistant cells, which is a characteristic sign of apoptosis-associated cell death induced by chemotherapeutic agents.

### 3.13 Torin 1 effectively inhibited cell migration and wound healing:-

The wound-healing assay offers a simple and reliable approach to evaluate cell migration, cell-to-cell interactions, and the wound-healing capacity of cells, providing valuable insights into their metastatic potential. As illustrated in the Figure 9, both HCT116 and REG-HCT116-R control cells demonstrated substantial migration and wound-healing abilities, achieving approximately 95% closure within 48 hours. Treatment of HCT116 cells with Regorafenib and Torin 1 at their respective IC50 concentrations leads to a notable decrease in their wound healing abilities. Conversely, Rapamycin does not exert a significant effect on wound healing in these cells. Interestingly, Reg-R-HCT116 cells exhibit wound-healing properties upon Torin 1 treatment, indicating a differential response compared to HCT116 cells. However, neither Rapamycin nor Regorafenib shows significant wound-healing effects in Reg-R-HCT116 cells. These findings suggest that Torin 1 is a potent inhibitor of migratory properties in the regorafenib resistant HCT116 cell line.

**Figure 9:**
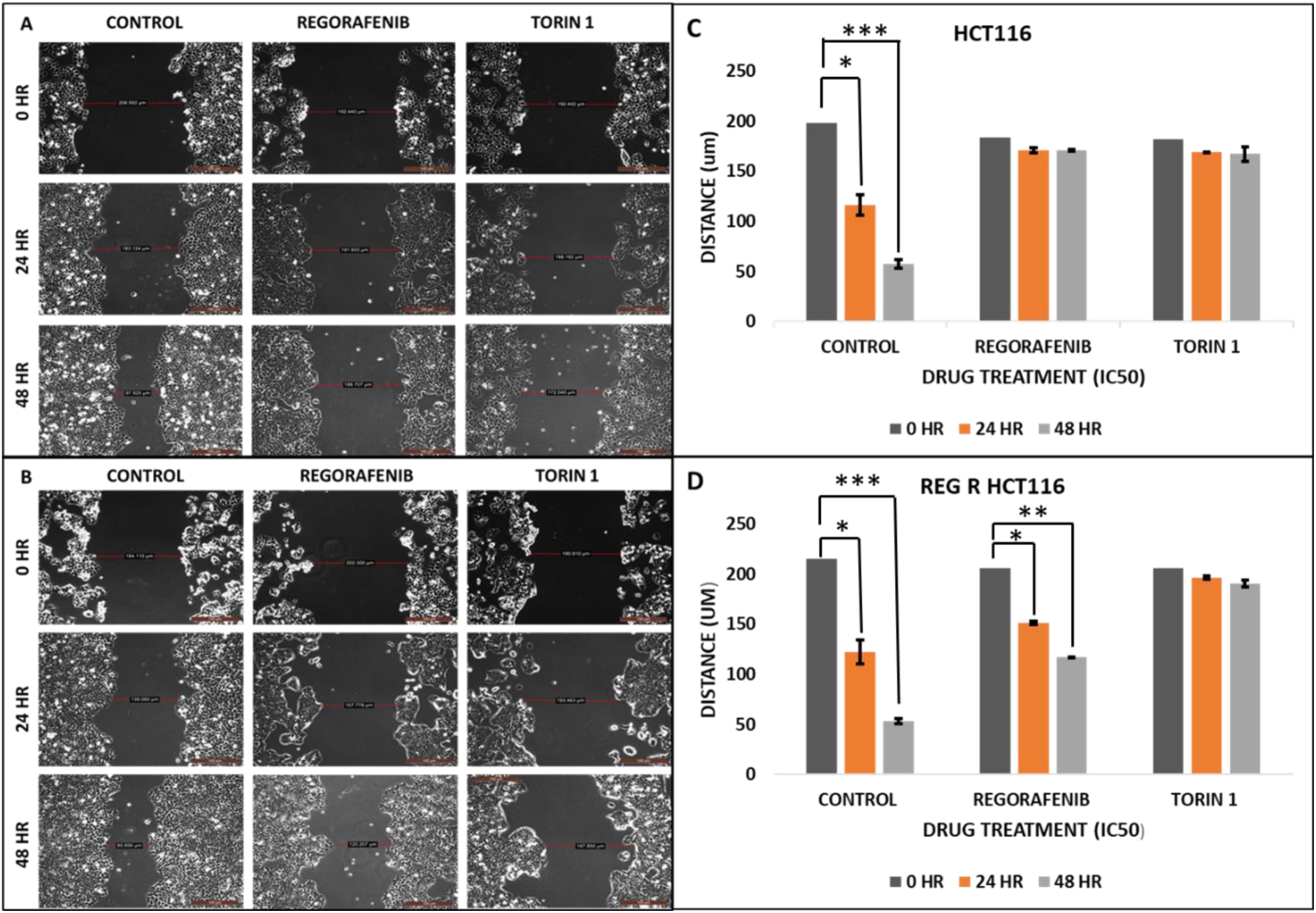
Inhibition of cell migration was assessed using a wound-healing assay after treatment with Regorafenib, Rapamycin, Torin 1, and a combination of Regorafenib and Torin 1 (Reg/Tor-1). (A) HCT116 and (B) REG-R-HCT116 cells were subjected to drug treatment, and wound closure was compared between treated and untreated cells at 24- and 48 hours post-incubation. The graphs illustrate the lengths of scratch wounds (measured in µm) at 0, 24, and 48 hours for both individual and combination treatments. A Bar Graph for the same is placed at the bottom for both cell lines in triplicate. A scale bar of 100 µm is provided for reference.

## 4. Discussion

Acquired resistance to regorafenib, a multikinase inhibitor approved for advanced colorectal cancer, remains a critical barrier to durable therapeutic responses. Although regorafenib targets multiple kinases—including VEGFR, FGFR, PDGFR, and RAF—the emergence of drug-tolerant cell subpopulations limits its long-term efficacy. The present study provides an integrated proteomics and phosphor-proteomics framework for understanding regorafenib resistance in colorectal cancer. By combining global proteome quantification with phosphosite-specific mapping, we reveal a coordinated molecular reprogramming centered on the PI3K/AKT/mTOR–eIF2 signaling axis, extensive Remodelling of ribosomal and translational machinery, and post-translational adaptation involving chromatin and cytoskeletal regulators. Together, these findings delineate a complex adaptive landscape through which tumor cells maintain growth and survival under sustained kinase inhibition.

### 4.1 Proteomic Remodelling reflects translational and metabolic adaptation

The proteomic profile of Reg-R-HCT116 cells revealed broad upregulation of ribosomal proteins (RPS6, RPS18, RPL27) and translation initiation factors (EIF5, EIF3B), strongly implicating enhanced translational capacity. The overrepresentation of ribosomal and eIF2 signaling pathways suggests that resistant cells overcome regorafenib-induced stress by maintaining protein synthesis despite MAPK blockade. Similar adaptive translation reprogramming has been documented in multiple drug-resistant cancers, where elevated ribosome biogenesis supports growth under metabolic or therapeutic stress (Truitt & Ruggero, *Nat Rev Cancer*, 2017).

The increased abundance of ribosomal proteins also implies persistent mTORC1 activation, which drives cap-dependent translation through phosphorylation of 4EBP1 and S6K. In our dataset, this was supported by enhanced expression of downstream translational regulators and the activation of eIF2-related pathways. The coupling of mTORC1-mediated translational activation with eIF2 stress signaling likely allows resistant cells to balance protein synthesis with proteostasis, ensuring survival under chronic drug exposure.

Concurrently, the proteomic data indicated shifts in metabolic pathways, including glycolytic enzymes and mitochondrial proteins, consistent with bioenergetic reprogramming. Drug-resistant cells frequently exhibit metabolic plasticity, switching between oxidative phosphorylation and glycolysis to maintain ATP supply and redox balance (Vasan et al., *Nat Rev Cancer*, 2022). Upregulation of these metabolic and translational modules reflects a survival adaptation that sustains biosynthetic demand despite regorafenib’s cytostatic effects.

### 4.2 Loss of epithelial identity and cytoskeletal reorganization

The downregulation of epithelial markers such as EPCAM and STMN1, together with lamin (LMNA, LMNB1, LMNB2) depletion, points to extensive cytoskeletal and nuclear Remodelling. The altered expression of these structural proteins is consistent with enhanced motility and mesenchymal-like plasticity, typical features of drug-tolerant phenotypes. Cytoskeletal Remodelling not only enables migration and anchorage-independent survival but also modulates receptor trafficking and signaling crosstalk, allowing resistant cells to sustain pro-survival inputs even under kinase inhibition (Sanhueza et al., *Oncogene*, 2021).

Interestingly, lamin downregulation was accompanied by upregulation of histone variants (H2B, H3, H4), suggesting chromatin condensation and apoptotic evasion. The proteomic pattern reflects a nucleus that is transcriptionally active but structurally altered—an adaptive chromatin state that favors survival gene expression and suppresses pro-apoptotic programs. Such chromatin compaction and histone enrichment have been reported as hallmarks of drug-induced quiescence and reversible tolerance in cancer cells (Sharma et al., *Cell*, 2010).

### 4.3 Phospho-proteomics reprogramming reveals selective activation of mTORC2–NDRG1 signaling

Phosphoproteome profiling uncovered distinct shifts in phosphorylation landscapes, revealing that regorafenib resistance is maintained by selective hyperactivation of cytoplasmic and translational signaling combined with dephosphorylation of nuclear and splicing regulators. Among the hyper-phosphorylated proteins, NDRG1 emerged as a major node with multiple upregulated phosphosites (Ser330–Ser366), known targets of SGK1 and mTORC2. NDRG1 phosphorylation promotes cell survival under hypoxia and nutrient deprivation, while non-phosphorylated NDRG1 induces apoptosis (Murakami et al., *Cancer Res*, 2019). Thus, its phosphorylation here signifies sustained mTORC2 activity despite regorafenib exposure.

In parallel, ACINUS, a chromatin-condensation factor cleaved during apoptosis, was also hyper-phosphorylated. Phosphorylation of ACINUS by AKT and SRPK1 prevents its apoptotic cleavage and maintains chromatin stability (Hu et al., *Nat Commun*, 2016). These phospho-proteomics signatures collectively indicate that AKT–mTORC2–NDRG1/ACINUS signaling forms a core anti-apoptotic circuit enabling regorafenib resistance.

This model is consistent with previous reports that chronic RAF inhibition reactivates parallel PI3K/AKT signaling, often through loss of negative regulators like NF1—a mutation we identified in Reg-R-HCT116 exomes. NF1 loss releases RAS inhibition, leading to compensatory PI3K activation and downstream mTORC2 engagement. This supports a dual-activation model: MAPK blockade → PI3K/AKT/mTORC2 hyperactivation → enhanced translation and survival.

### 4.4 Dephosphorylation of nuclear and splicing regulators as a stabilizing adaptation

While cytoplasmic proteins exhibited hyper-phosphorylation, the nuclear phosphoproteome was dominated by dephosphorylation events. Key RNA-processing and chromatin Remodelling factors—including BAZ1A, ARID1A, TRIM28, SAFB, and HNRNPA2B1—were hypo-phosphorylated. Phosphorylation regulates their activity in transcription, splicing, and DNA damage response; its loss may “lock” these regulators into a survival-permissive state. For instance, hypo-phosphorylated ARID1A (a SWI/SNF subunit) and TRIM28 are associated with chromatin condensation and transcriptional repression of pro-apoptotic genes (Zhou et al., *Cell Rep*, 2021). Similarly, decreased phosphorylation of splicing factors like HNRNPM and TRA2B could alter isoform usage to favor oncogenic or anti-apoptotic transcripts (Sebestyén et al., *Genome Biol*, 2016).

This dual pattern—cytoplasmic hyper-phosphorylation and nuclear hypo-phosphorylation—suggests a spatial reorganization of signaling: phosphorylation activity concentrates in growth and stress-response compartments while being suppressed in the nucleus to maintain genomic stability and transcriptional quiescence. This phospho-compartmentalization may be a hallmark of regorafenib tolerance, allowing metabolic activity without triggering DNA-damage-induced apoptosis.

### 4.5 Cytoskeletal and vesicular hyper-phosphorylation reinforces survival and signaling plasticity

Hyper-phosphorylation of CAV1, CTTN, and RANBP2 indicates active membrane trafficking and cytoskeletal Remodelling, processes that sustain receptor recycling and intracellular signaling during drug stress. CAV1 phosphorylation facilitates caveolae-mediated endocytosis and downstream activation of Src and PI3K, bypassing RAF inhibition. CTTN phosphorylation enhances actin polymerization, promoting motility and invasion, while RANBP2 regulates nucleocytoplasmic transport and SUMOylation of signaling proteins, stabilizing survival kinases. Together, these events depict a phospho-proteomic shift toward enhanced vesicular dynamics, enabling resistant cells to continuously modulate receptor availability and sustain autocrine signaling.

### 4.6 Integration of proteome and phosphoproteome: persistent mTOR/eIF2 activation as a central node

Integrative network analysis of both datasets converges on PI3K/AKT/mTOR–eIF2 signaling as the principal adaptive pathway. Upregulation of ribosomal and translational proteins in the proteome coupled with phosphorylation of mTORC2 substrates (NDRG1, RICTOR, AKT Ser473) underscores the persistence of anabolic signaling. eIF2 pathway activation reflects translational stress adaptation, ensuring selective translation of stress-response mRNAs. Such translational plasticity has been implicated in drug tolerance across cancer types, providing a rapid-response mechanism independent of genetic mutation (Leprivier et al., *Nat Rev Cancer*, 2020).

Importantly, both Ingenuity Pathway Analysis and KEGG enrichment pinpointed *Torin1* and *Sirolimus* as predicted upstream inhibitors capable of suppressing these activated nodes. Torin1, a dual mTORC1/2 inhibitor, effectively reversed regorafenib resistance in our functional assays, demonstrating cytotoxic synergy and reactivation of apoptotic markers. This pharmacological validation supports the proteo-signaling model derived from our omics data and positions dual mTOR inhibition as a viable strategy for overcoming regorafenib resistance.

### 4.7 Phospho-signaling balance and potential regulatory phosphatases

The extensive dephosphorylation observed across nuclear proteins suggests that resistance may involve altered phosphatase dynamics, potentially mediated by PP1 and PP2A complexes. We noted hypo-phosphorylation of PPP1R12A, a PP1 regulatory subunit, indicating deregulated phosphatase control. Phosphatase imbalance could stabilize chromatin and RNA-processing machinery by preventing phosphorylation-dependent degradation or turnover. Restoring phospho-homeostasis via phosphatase modulation might therefore re-sensitize resistant cells, an emerging therapeutic avenue supported by recent work on PP2A reactivation in drug-resistant cancers (Perrotti & Neviani, *Nat Rev Cancer*, 2023).

### 4.8 Model of resistance and therapeutic implications

Collectively, our data support a multi-layered model of regorafenib resistance (Fig. 8). Chronic kinase inhibition triggers a compensatory shift from RAF–MEK–ERK dependency to PI3K/AKT/mTOR–eIF2–driven translation and metabolic survival. Concurrent cytoskeletal Remodelling enhances motility and receptor recycling, while chromatin and RNA-processing dephosphorylation stabilize transcriptional programs favoring persistence. This integrated proteo-phospho adaptation collectively redefines the signaling landscape toward a stress-tolerant, apoptosis-resistant phenotype.

From a therapeutic perspective, this rewired network suggests several actionable vulnerabilities:

1. **mTORC1/2 inhibition (Torin1, AZD8055):** Effective in blocking downstream translational activation and reversing regorafenib resistance.
2. **SGK1–NDRG1 axis targeting:** May disrupt cytoprotective signaling linked to mTORC2.
3. **Splicing and phosphatase modulation:** Restoring phosphorylation dynamics of nuclear regulators could re-enable apoptotic gene expression.
4. **Combination strategies:** Co-targeting mTOR and regorafenib-responsive kinases could delay or prevent resistance emergence.

## 5. Future directions

Future work should extend these findings through temporal phospho-proteomics to capture the dynamic evolution of resistance and through *in vivo* xenograft validation to assess translational relevance. Given that NF1 loss and mTOR activation are recurrent in resistant colorectal tumors, stratifying patients based on these molecular signatures could guide personalized regorafenib–mTOR inhibitor combinations. Furthermore, single-cell proteomics may reveal heterogeneity in phospho-states within resistant populations, identifying subclonal drivers of relapse.

## 6. Conclusion

This study presents an integrated proteomic and phospho-proteomics dissection of regorafenib resistance in colorectal cancer, revealing a complex adaptive architecture driven by persistent PI3K/AKT/mTOR–eIF2 signaling and post-translational rewiring. The resistant state is characterized by translational activation, metabolic plasticity, cytoskeletal reorganization, and chromatin stabilization, collectively ensuring survival under kinase inhibition. Pharmacologic inhibition of mTORC1/2 by Torin1 effectively disrupts this network, reinstating apoptotic sensitivity. These findings provide a molecular blueprint for designing rational combination therapies to overcome regorafenib resistance and underscore the importance of phospho-proteomic interrogation in decoding adaptive tumor evolution.

## Author contributions

Dr. Jayadev and Dr. Ashutosh Singh designed the study. Dr. Deepu Sharma and Dr. Rohit Verma performed proteome and phosphor-proteome analysis and contributed to data curation. Dr. Fayyaz Rasool performed the experiments. Dr. Fayyaz Rasool and Dr. Jayadev helped to finalize the manuscript critically.

## Declarations

AI tools such as ChatGPT were used to correct grammar errors and enhance readability.

