## Supplementary Data. for "Cellular mechanisms of acquired drug resistance against regorafenib in colon cancer cells"

**S1:** List of the Primer sequence used in the study.

| S. No. | Primer Name | Primer Sequence |
|---|---|---|
| 1. | ACTA2 FW | TCCCTTGAGAAGAGTTACGA |
| 2. | ACTA2 RV | GATGCTGTTGTAGGTGGTTT |
| 3. | ACTG2 FW | TCAAGGAGAAGCTGTGCTAT |
| 4. | ACTG2 RV | ATAAAGGAAGGCTGGAAGAG |
| 5. | EIF2AK2 FW | GAATCAACATCCACACTTCC |
| 6. | EIF2AK2 RV | GAAACCTGCTGAAAGATCAC |
| 7. | EIF5 FW | GAACTCCTTGCCTGATCTAA |
| 8. | EIF5 RV | AATGGTATAAGCCACTGCTC |
| 9. | RPS13 FW | ACGTGAAGGAGCAGATTTAC |
| 10. | RPS13 RV | TCTCAGGATTACACCGATCT |
| 11. | RPS3A FW | GTAGAGATGGGGTTTCATCA |
| 12. | RPS3A RV | CATTTCCTGTCCTTCAACTC |
| 13. | RPS18 FW | GATATGCTCATGTGGTGTTG |
| 14. | RPS18 RV | CCATCCTTTACATCCTTCTG |
| 15. | RPS15 FW | TACAAGCCCGTAAAGCAT |
| 16. | RPS15 RV | CCATTACTTGAGAGGGATGA |
| 17. | RPS4Y1 FW | AGTTGTGCAAAGTGAGGAAG |
| 18. | RPS4Y1 RV | ATCTTGCCAGTCCCTAAATC |
| 19. | RPL37 FW | CTTGACCTGACCCATGTATT |
| 20. | RPL37 RV | TAGGACTGCACGTTTTACCT |
| 21. | RPL10L FW | CCGGTATTGTAAGAACAAGC |
| 22. | RPL10L RV | TTCTACCCAGGTCAAAGATG |
| 23. | RPL35A FW | AGAGTCATCTGGGGAAAAGT |
| 24. | RPL35A RV | TAAATCCTTGAGGGGTACAG |
| 25. | RPL27 FW | TCGCCAAGAGATCAAAGA |
| 26. | RPL27 RV | AGAGTACCTTGTGGGCATTA |
| 27. | RPL22L1 FW | CTTTGTAGGCACCAGAAACT |
| 28. | RPL22L1 RV | GATTCCAACCGAGTGAACTA |
| 29. | RPL9 FW | GAGAATGGGTCTCTTGTTGA |
| 30. | RPL9 RV | TCTTTCTGGGCTTGAGATAC |
| 31. | BCL2 FW | CGATCTGGAAATCCTCCTAA |
| 32. | BCL2 RV | CCATCAATCTTCAGCACTCT |
| 33. | CASPASE 3 FW | GGCGTGTCATAAAATACCAG |
| 34. | CASPASE 3 RV | ACAAAGCGACTGGATGAA |
| 35. | CASPASE 9 FW | GCTCTTCCTTTGTTCATCTC |
| 36. | CASPASE 9 RV | GTAAGGTTTTCTAGGGTTGG |
| 37. | RICTOR FW | TGTTGTGACTGAGGAGTTCA |
| 38. | RICTOR RV | GCACGATGAGGAAGAATAGT |
| 39. | RAPTOR FW | CACTTCTCGTTTTCATCTGG |
| 40. | RAPTOR RV | AGGCGTAGTTCTTTTCCTCT |
| 41. | PARP FW | GTTTTGTGTTGTGTCTGTGG |
| 42. | PARP RV | GAATCTCTCTCCAGCCTTTT |
| 43. | 18S-FW | GCAATTATTCCCCATGAACG |
| 44. | 18S-RV | GGCCTCACTAAACCATCCAA |
| 45. | DKK1 FW | ACAACTACCAGCCGTACCC |
| 46. | DKK1 RV | TGCAGGCGAGACAGATTTGC |

| | | |
|---|---|---|
| 47. | CRTC2 FW | GCTCTGACTCTGCCCTTCAT |
| 48. | CETC2 RV | CTCAATAGCAGGTACTTTGGGG |
| 49. | VDAC3 FW | CCAAGTCTTGTAGTGGAGTGATGG |
| 50. | VDAC3 RV | GTCAGTTTCAACCCTTCAGCC |
| 51. | TJP FW | GTTTATTTGGGCTGTGGCGTG |
| 52. | TJP RV | TCCTCCATTGCTGTGCTCTTG |
| 53. | RAP1 FW | CTGCCTGAGAGAAGTAGAAGATCG |
| 54. | RAP1 RV | CTTCCCAACGCCTCCTGAAC |
| 55. | FMR1 FW | CCTGAACTCAAGGCTTGGCA |
| 56. | FMR1 RV | TCTCTTCCTCTGTTGGAGCTTTA |
| 57. | NUP107FW | TCAGAAAAGAGTTTTACTTCAGGCA |
| 58. | NUP107RV | GGGGTAACTATAAACAAAAGGTTGG |
| 59. | HECTD1 FW | ATCCCTGCAACGTCGAGC |
| 60. | HECTD1 RV | TTTCGTACTCTTCTTCTTCCTGCT |

**S2:** HCT116 Control and HCT116 Regorafenib Resistance cells were cultured and seed for Immunocytochemistry to check the expression of p-ACINUS.

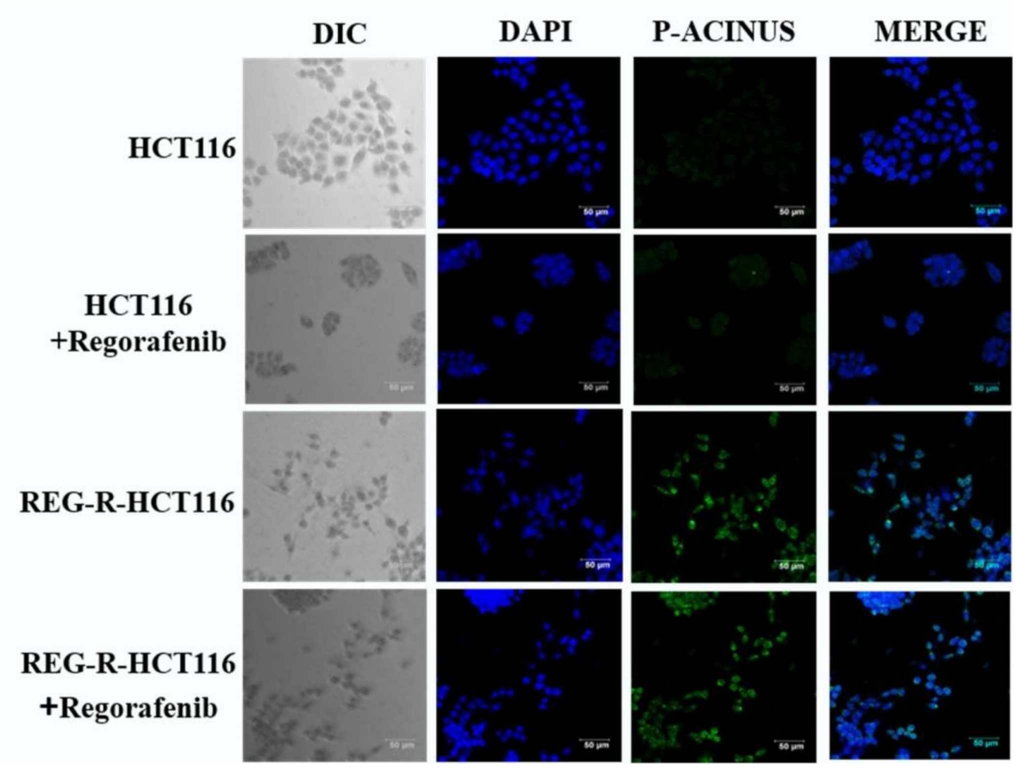

**S3:** Comparison of the quantified protein densitometry analysis using Image J software. The values of the densitometric analysis of two biological replicates were used and standard deviations were calculated.

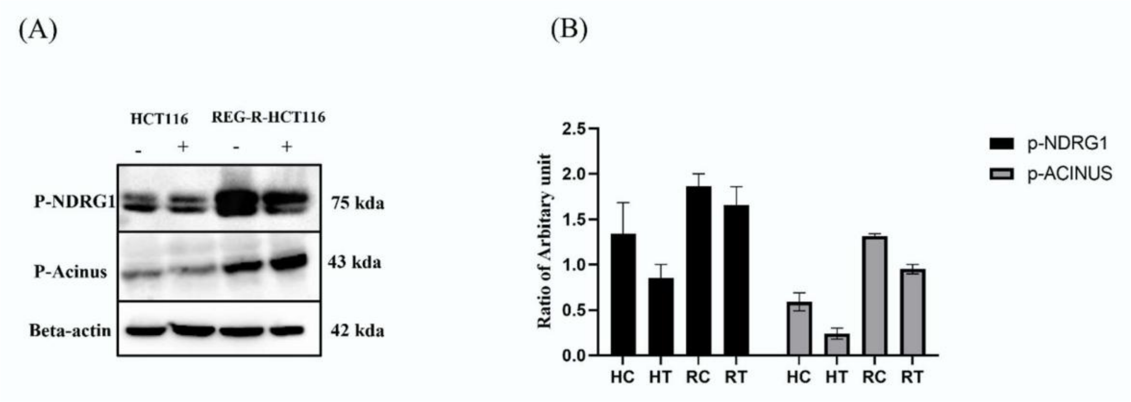


**S4**: List of the genes found upregulated in Reg-R-HCT116 cells as compare to control in proteome data analysis. These genes are selected for validation of proteome data and upregulation of eIF2 signalling.

| **Gene Symbol** | **Description** | **Abundance Ratio (log2): (T, Sample) / (C, Control)** |
|---|---|---|
| ACTA2 | Actin, aortic smooth muscle OS=Homo sapiens GN=ACTA2 PE=1 SV=1 | 1.6702037 |
| ACTG2 | Actin, gamma-enteric smooth muscle OS=Homo sapiens GN=ACTG2 PE=1 SV=1 | 1.6702037 |
| EIF2AK2 | Interferon-induced, double-stranded RNA-activated protein kinase OS=Homo sapiens GN=EIF2AK2 PE=1 SV=2 | 1.8780897 |
| EIF5 | Eukaryotic translation initiation factor 5 OS=Homo sapiens GN=EIF5 PE=1 SV=2 | 1.9735516 |
| RPS13 | 40S ribosomal protein S13 OS=Homo sapiens GN=RPS13 PE=1 SV=2 | 1.5084098 |
| RPS3A | 40S ribosomal protein S3a OS=Homo sapiens GN=RPS3A PE=1 SV=2 | 1.9995339 |
| RPS18 | 40S ribosomal protein S18 OS=Homo sapiens GN=RPS18 PE=1 SV=3 | 2.1524181 |

| | | |
|---|---|---|
| RPS15 | 40S ribosomal protein S15 OS=Homo sapiens GN=RPS15 PE=1 SV=2 | 2.4595619 |
| RPS4Y1 | 40S ribosomal protein S4, Y isoform 1 OS=Homo sapiens GN=RPS4Y1 PE=1 SV=2 | 2.5739876 |
| RPL37 | 60S ribosomal protein L37 OS=Homo sapiens GN=RPL37 PE=1 SV=2 | 1.5368963 |
| RPL10L | 60S ribosomal protein L10-like OS=Homo sapiens GN=RPL10L PE=1 SV=3 | 1.598766 |
| RPL35A | 60S ribosomal protein L35a OS=Homo sapiens GN=RPL35A PE=1 SV=2 | 1.7641291 |
| RPL27 | 60S ribosomal protein L27 OS=Homo sapiens GN=RPL27 PE=1 SV=2 | 1.8034472 |
| RPL22L1 | 60S ribosomal protein L22-like 1 OS=Homo sapiens GN=RPL22L1 PE=1 SV=2 | 1.9852897 |
| RPL9 | 60S ribosomal protein L9 OS=Homo sapiens GN=RPL9 PE=1 SV=1 | 1.9969607 |
